# Accelerating functional gene discovery in osteoarthritis

**DOI:** 10.1101/836221

**Authors:** Natalie C. Butterfield, Katherine F. Curry, Julia Steinberg, Hannah Dewhurst, Davide Komla-Ebri, Naila S. Mannan, Anne-Tounsia Adoum, Victoria D. Leitch, John G. Logan, Julian A. Waung, Elena Ghirardello, Lorraine Southam, Scott E. Youlten, J Mark Wilkinson, Elizabeth A. McAninch, Valerie E. Vancollie, Fiona Kussy, Jacqueline K. White, Christopher J. Lelliott, David J. Adams, Richard Jacques, Antonio C. Bianco, Alan Boyde, Eleftheria Zeggini, Peter I. Croucher, Graham R. Williams, J. H. Duncan Bassett

**Affiliations:** Molecular Endocrinology Laboratory, Department of Metabolism, Digestion and Reproduction, Imperial College London, London W12 0NN, UK; Institute of Translational Genomics, Helmholtz Zentrum München – German Research Center for Environmental Health, 85764 Neuherberg, Germany; Wellcome Trust Sanger Institute, Hinxton, Cambridge CB10 1SA, UK; Cancer Council NSW, Sydney, New South Wales 2000, Australia; The Garvan Institute of Medical Research and St. Vincent’s Clinical School, University of New South Wales Medicine, Sydney, New South Wales 2010, Australia; Department of Oncology and Metabolism, University of Sheffield, Sheffield S10 2RX, UK; Centre for Integrated Research into Musculoskeletal Ageing and Sheffield Healthy Lifespan Institute, University of Sheffield, Sheffield S10 2TN, UK; Division of Endocrinology and Metabolism, Rush University Medical Center, Chicago, IL 60612, USA; The Jackson Laboratory, Bar Harbor, ME 04609, USA; School of Health and Related Research (ScHARR), University of Sheffield, Sheffield S1 4DA, UK; Section of Adult and Pediatric Endocrinology, Diabetes & Metabolism, Department of Medicine, University of Chicago, Chicago, IL 60637, USA; Dental Physical Sciences, Queen Mary University of London, Mile End Road, London E1 4NS, UK

## Abstract

Osteoarthritis causes debilitating pain and disability, resulting in a huge socioeconomic burden, yet no drugs are available that prevent disease onset or progression. Here, we develop, validate and use rapid-throughput imaging techniques to identify abnormal joint phenotypes in unselected mutant mice generated by the International Knockout Mouse Consortium. We identify 14 genes with functional involvement in osteoarthritis pathogenesis, including the homeobox gene *Pitx1*, and functionally characterize 6 candidate human osteoarthritis genes in mouse models. We demonstrate sensitivity of the methods by identifying age-related degenerative joint damage in wild-type mice. Finally, we generate mutant mice with an osteoarthritis-associated polymorphism in the *Dio2* gene by *Crispr/Cas9* genome editing and demonstrate a protective role in disease onset with public health implications. This expanding resource of unselected mutant mice will transform the field by accelerating functional gene discovery in osteoarthritis and offering unanticipated drug discovery opportunities for this common and incapacitating chronic disease.

---

Osteoarthritis is the commonest cause of joint destruction, pain and disability. Joint replacement for end-stage disease remains the only treatment, leading to an escalating healthcare crisis in our obese and ageing society. Osteoarthritis is a complex trait and the 86 reported genome-wide associated loci explain only a small proportion of its heritability, which is estimated between 40-70%^1–3^.

Osteoarthritis is characterized by articular cartilage damage and loss, together with structural abnormalities of subchondral bone and low-grade chronic joint inflammation. It is unknown which of these processes trigger disease or which represent secondary responses to joint destruction^4^. It is also uncertain whether the pathogenesis of osteoarthritis reflects an abnormal response to injury involving defective stem cell recruitment and abnormal cell proliferation, differentiation, metabolism, apoptosis and senescence^5, 6^.

Chondrocytes in healthy articular cartilage are resistant to terminal differentiation whereas they revert to a developmental program following injury, in which they proliferate and undergo hypertrophic differentiation with accelerated cartilage mineralization^7^. Osteoarthritis pathogenesis involves cross-talk between the synovium, articular cartilage and subchondral bone^8^, although the timing of bone remodeling relative to cartilage degradation remains uncertain^9^. Nevertheless, increases in apoptotic and senescent chondrocytes are triggered by processes including endoplasmic reticulum stress^10^. Senescent cells express a secretory phenotype that contributes to inflammation, vascular invasion^11^ and cartilage breakdown via key pathways that stimulate matrix metalloproteases^12^ and aggrecan-specific proteinases^13^.

Despite the profound clinical and socio-economic impacts of osteoarthritis, our understanding of its genetic basis is in its infancy. We hypothesized that accelerating gene discovery in osteoarthritis will increase understanding of joint physiology and disease pathogenesis and facilitate identification of new drug targets that prevent or delay joint destruction. Studies of extreme phenotypes in humans have underpinned identification of the molecular basis of single gene disorders and mechanisms of complex disease, and resulted in revolutionary new treatments^14, 15^. Analogous to our gene discovery studies in osteoporosis^16–20^, we proposed that a joint-specific extreme phenotype screen in mutant mice would accelerate functional gene discovery in osteoarthritis.

Unselected mutant mice are generated at the Sanger Institute as part of the International Mouse Phenotyping Consortium (IMPC). Mice undergo broad phenotyping using the International Mouse Phenotyping Resource of Standardized Screen (IMPReSS) that is completed at 16 weeks of age when tissues are harvested for further analysis. In the Origins of Bone and Cartilage Disease (OBCD) Project we collaborate with IMPC and receive knee joints for analysis. Rapid-throughput phenotyping of the mouse knee requires quantitative imaging; this presents a complex and unsolved challenge that relates to anatomical size, three-dimensional complexity, image resolution and the necessity to maintain joints in their native fully-hydrated state.

Here we present the invention, optimization, validation and application of a rapid-throughput multimodality imaging pipeline to phenotype the mouse knee. We analyzed 50 unselected mouse lines, identifying seven (14%) with markedly abnormal phenotypes. A systematic prioritization strategy identified seven further lines, resulting in 14 genes (28%) with evidence for a functional role in osteoarthritis pathogenesis. The four leading candidates were *Pitx1*, *Bhlhe40*, *Sh3bp4* and *Unk*. We interrogated the database of joint phenotypes from unselected mouse lines with 409 genes differentially expressed in human osteoarthritis cartilage. This resulted in an enriched yield of abnormal joint phenotypes in six (75%) of eight lines for which data were available, including *Unk*. We then applied the novel pipeline to characterize the early features of age-related joint degeneration in one-year old mice and demonstrated its sensitivity to detect disease onset as well as surgically provoked late-stage disease, paving the way for application to analysis of drug intervention studies. Finally, we generated *Crispr/Cas9* mutant mice with a Thr92Ala polymorphism in the *Dio2* gene that is orthologous to the human variant associated with osteoarthritis susceptibility. The Ala92 allele conferred protection against early-onset osteoarthritis, challenging current understanding and providing new implications for public health.

## Results

### Invention and optimization of novel imaging methods

We established a rapid-throughput joint phenotyping pipeline (OBCD joint pipeline), which applies three complementary imaging approaches to characterize features of osteoarthritis in mutant mice that include articular cartilage damage and loss, together with abnormalities of subchondral bone structure and mineralization (Figure 1). Iodine contrast-enhanced micro-computerized tomography (ICEμCT) was developed to determine articular cartilage volume (ACV), median thickness (Median AC Th), maximum thickness (Max AC Th), subchondral bone volume/tissue volume (SC BV/TV), trabecular thickness (SC Tb.Th), trabecular number (SC Tb.N) and bone mineral density (SC BMD). Joint surface replication (JSR) was invented and optimized to quantify articular cartilage surface damage. Subchondral bone X-ray microradiography (scXRM) was developed from previous protocols^21, 22^ to determine subchondral bone mineral content (SC BMC) (Figures 1, 2 and Supplementary Figure 1).

**Figure 1.**
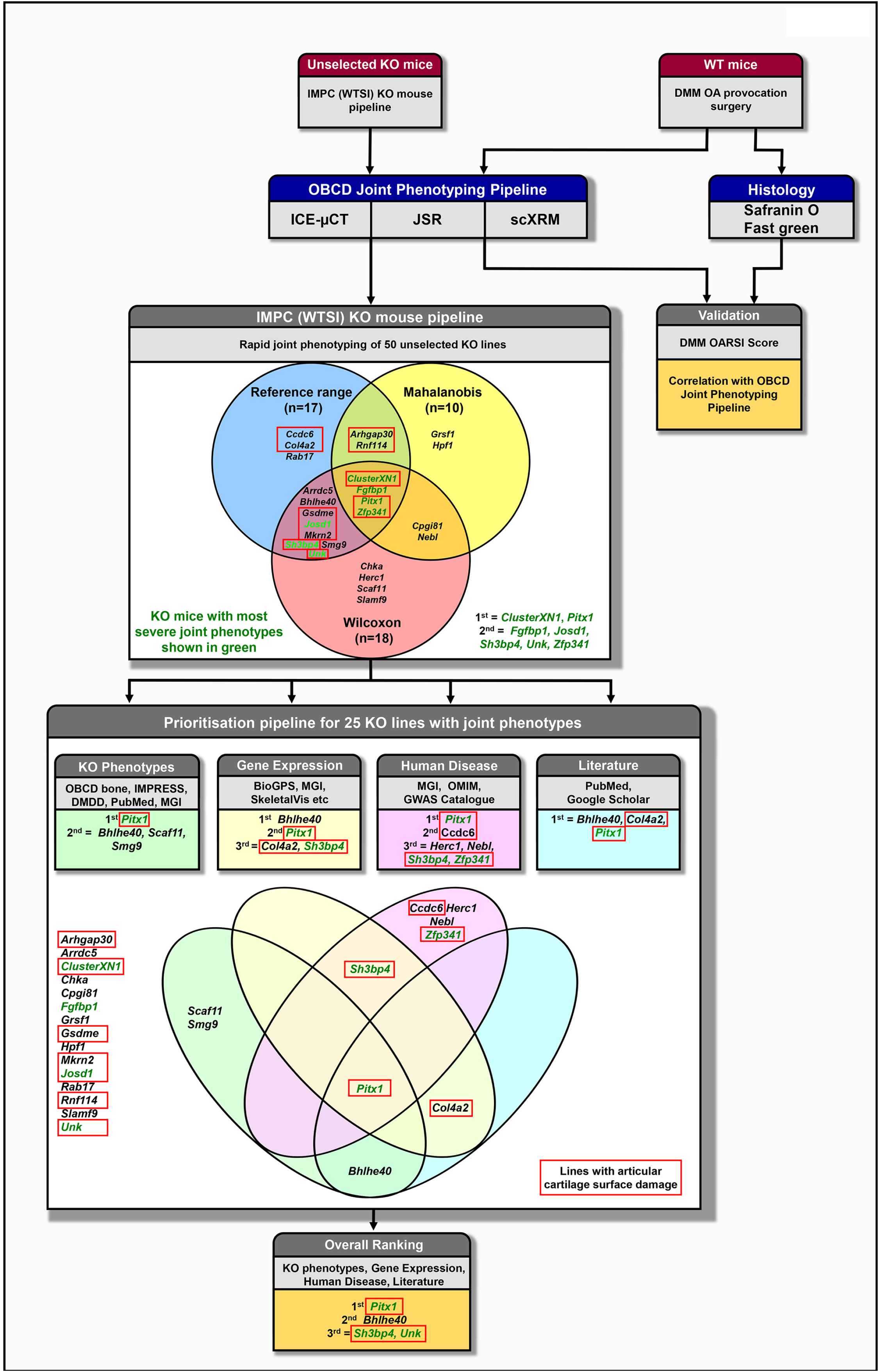
Rapid-throughput joint phenotyping identifies new osteoarthritis genes. Knee joints from unselected mutant mice generated at the Wellcome Trust Sanger Institute (WTSI) for the International Mouse Phenotyping Consortium (IMPC) are analyzed by three novel imaging modalities; iodine contrast-enhanced micro computerized tomography (ICEμCT), joint surface replication (JSR) and subchondral bone X-ray microradiography (scXRM). The methods were validated by comparison with scoring of histological sections (Osteoarthritis Research Society International (OARSI) histology after destabilization of the medial meniscus (DMM) provocation surgery. 25 mouse lines (3-way Venn diagram) had outlier phenotypes relative to wild-type reference data following statistical analyses. Further prioritization was based on additional skeletal abnormalities identified in mutant mice, gene expression, association with human disease and literature searching. Boxes and 4-way Venn diagram show top ranked genes in each category. Pitx1, Bhlhe40, Sh3bp4 and Unk were the top ranked genes. Green text: most severely abnormal joint phenotypes. Red boxes indicate knockout lines with increased articular cartilage surface damage.

**Figure 2.**
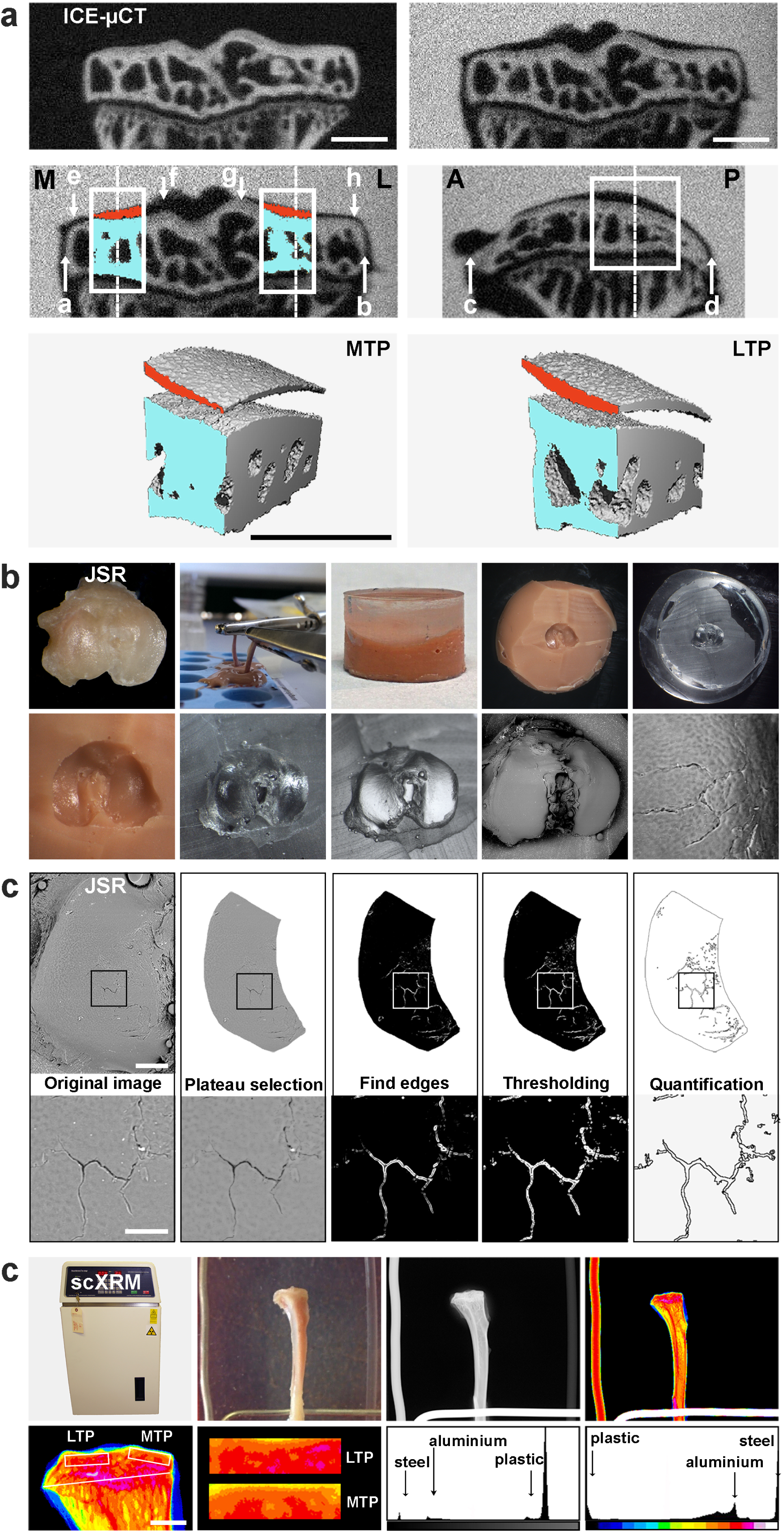
Novel imaging methods. ***a.* Iodine contrast enhanced micro computerized tomography (ICEμCT).** Coronal views of conventional uCT scan (top left) and ICEµCT scan (top right), at 2μm voxel resolution; mineralized tissue is white. In ICEµCT scan, contrast agent is at similar X-ray absorption to mineralized tissue, and soft tissue including articular cartilage is black. Coronal (middle left) and sagittal (middle right) views of ICEµCT scans with VOIs (volumes of interest, white box) scaled according to a-b (medial-lateral) and c-d (anterior-posterior) tibial dimensions, and positioned midway between plateau edges (medial plateau, MTP, e-f, lateral plateau, LTP, g-h, with a 7% shift medially, see Methods). Plateau midpoints: dashed lines. M: medial, L: lateral, A: anterior, P: posterior. ICEµCT VOIs used for quantitative analyses (MTP, bottom left, LTP, bottom right) with subchondral bone in blue and articular cartilage in red. Scale bars = 500μm. ***b.* Joint surface replication (JSR) - Generation of mouse tibia knee joint surface replicas.** Upper images left to right show: surface of disarticulated mouse tibia; limbs immersed in impression medium for moulding; mould (brown) overlaid with casting resin; surface of mould after removal of cast; and surface of cast after removal from mould. Lower images left to right show: detail of mould surface; detail of cast surface; carbon-coated cast; back-scattered electron-scanning electron microscopy (BSE-SEM) image of whole tibia; and high-power BSE-SEM image showing fibrillations in articular cartilage surface induced following DMM surgery. ***c.* Joint surface replication (JSR) - Quantitation of joint surface damage.** Upper images left to right show: BSE-SEM view of lateral tibial plateau; selection of whole plateau region for analysis; automated identification of surface damage edges; thresholding of damage to capture all damage areas and damage detection. Lower images left to right show high power views to demonstrate joint surface damage detail. Scale bars = 500μm (upper), 100μm (lower). ***d.* Subchondral bone X-ray microradiography (scXRM).** Upper images left to right show: Faxitron MX-20; mouse tibia alongside aluminium (left), plastic (right) and steel (bottom) standards; plain X-ray microradiograph grey scale image of tibia and standards; image pseudocoloured by application of 16 colour look-up table in which lower bone mineral content (BMC) is yellow and higher BMC is pink/red. Lower images left to right show: pseudocoloured image of proximal tibia with scaled (to width of tibia, white line) subchondral regions of interest (ROI), white boxes (LTP; lateral tibial plateau, MTP; medial tibial plateau); high power pseudocoloured views of LTP and MTP ROIs; grey scale pixel distribution in relation to standards; grey scale distribution stretched to plastic and steel standards. Scale bar = 1mm.

### Validation of new imaging methods

The OBCD joint pipeline methods were validated by comparison with Osteoarthritis Research Society International (OARSI) histological scoring^23^ of knees from wild-type (WT) mice following destabilization of the medial meniscus (DMM)^13, 24^. One cohort of WT mice (n=16) was phenotyped using ICEμCT, JSR and scXRM in the OBCD joint pipeline, and a second cohort (n=11) was analyzed by OARSI scoring (Figure 1). DMM surgery resulted in decreased ACV and Max AC Th with a marked increase in AC damage. This cartilage destruction was accompanied by increased SC BV/TV, increased SC Tb.Th, decreased SC Tb.N and increased SC BMC. These abnormalities were consistent with extensive cartilage damage demonstrated by OARSI-scored histology (Figure 3 and Supplementary Figure 2). OARSI analysis was also undertaken on three joints that were phenotyped in the OBCD joint pipeline and found to have features indicating mild, intermediate and severe osteoarthritis following DMM surgery. OARSI scoring was concordant with the severity of abnormalities identified by ICEμCT, JSR and scXRM, thus validating the use of these imaging modalities to characterize osteoarthritis (Figure 3, Supplementary Figures 2 and 3).

**Figure 3.**
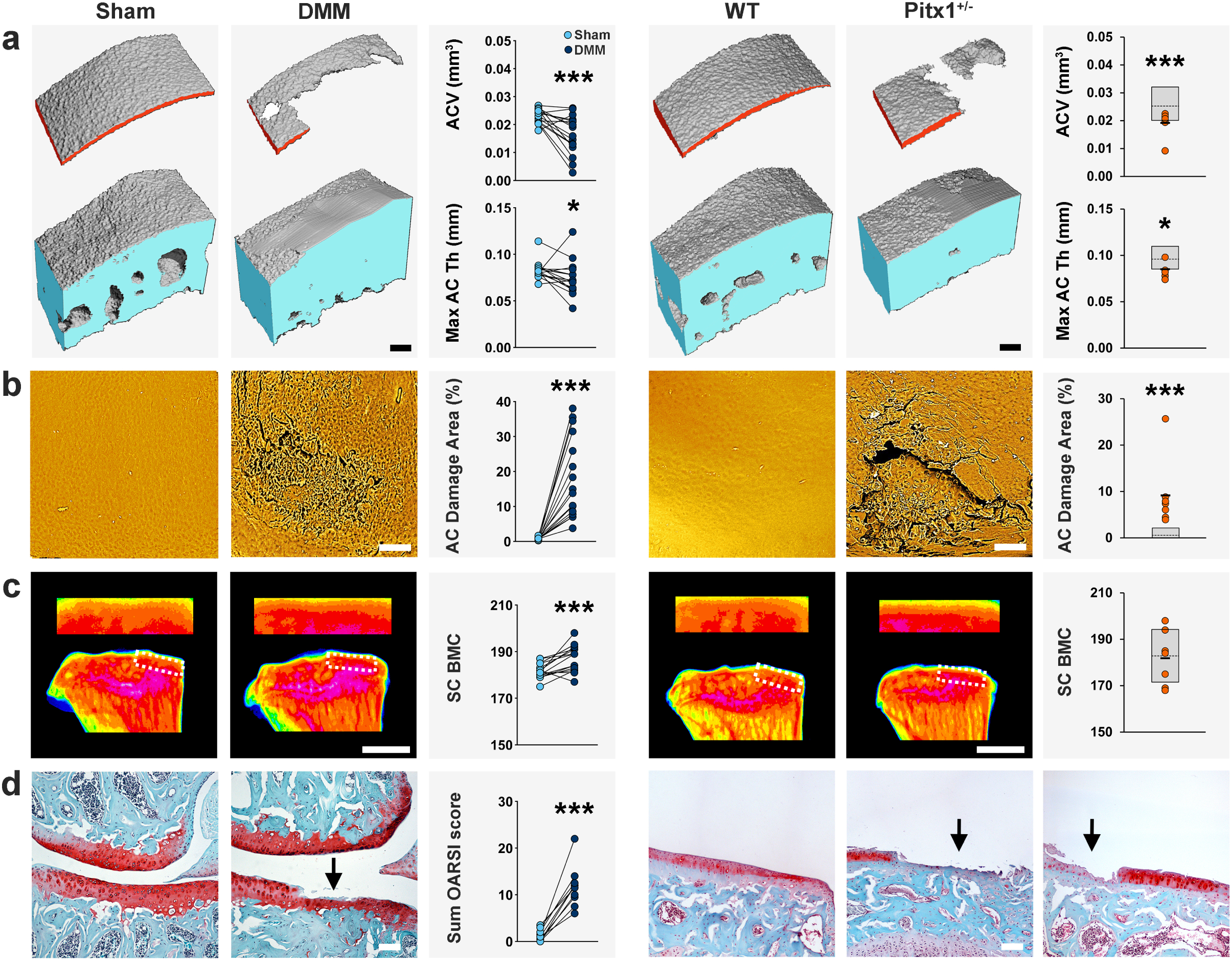
Severe early onset osteoarthritis in heterozygous Pitx1^+/-^ mice is similar to extent of joint damage following osteoarthritis provocation surgery. ***a.*** Iodine contrast enhanced μCT images of medial tibial plateau articular cartilage (red) and subchondral bone (blue) from 22-week-old WT male mice 12 weeks after sham operation or following destabilization of medial meniscus surgery (DMM; n=11) to provoke osteoarthritis, and 16-week-old male WT and heterozygous Pitx1^+/-^ mice (n=7). Graphs show decreased articular cartilage volume (ACV) and maximum articular cartilage thickness (Max AC Th). ***b.*** Back-scattered electron-scanning electron microscopy images of tibial plateau joint surface replicas from sham and DMM operated WT mice, and WT and Pitx1^+/-^ mice. Graphs show increased articular cartilage (AC) damage area. ***c.*** Pseudocoloured X-ray microradiography images of proximal tibia and the medial tibial plateau subchondral bone region of interest (dashed box) from sham and DMM operated WT mice, and WT and Pitx1^+/-^ mice. Low bone mineral content (BMC) is yellow and high BMC is pink. Graphs show increased subchondral BMC (SC BMC; DMM only). ***d.*** Coronal sections of knee joint compartments stained with Safranin-O/Fast green from sham and DMM operated WT mice, and from WT and two examples of Pitx1^+/-^ mice with extensive joint destruction. Graph shows the summed OARSI score on the medial tibial plateau in DMM operated mice (n=11). Arrows indicate areas of cartilage destruction. Grey boxes: reference ranges, dashed lines: population means/medians, orange circles: individual mutant samples and black horizontal lines: sample means. Scale bars = 100μm (A, B, D) and 1mm (C). *P<0.05, ***P<0.001 (DMM), *P<0.00568, ***P<0.0001 (Pitx1^+/-^).

### Definition of WT reference ranges

Phenotype datasets for all parameters were obtained from 100 WTmice. Reference ranges, coefficients of variation, estimates of skewness and kurtosis, normality, repeatability and power calculations were determined (Supplementary Table 1). The reference range for each parameter was defined as either (i) the mean +/- 2.0 standard deviations for normally-distributed data, or (ii) the median and 2.5^th^ −97.5^th^ percentile range for non-normally distributed data (Supplementary Figure 1).

### Rapid-throughput joint phenotyping identifies new osteoarthritis genes

To identify new genes that cause osteoarthritis, we used the OBCD joint pipeline to analyze knees from 16-week-old male mice (n=3-7) from 50 unselected mutant lines generated in an identical *C57BL6/N;C57BL6/NTac* genetic background (Supplementary Table 3).

Rigorous statistical approaches were used to determine which lines displayed robust outlier phenotypes (Figure 1, Supplementary Tables 3 and 4). Lines with outlier phenotypes were identified: (i) when the mean value for an individual parameter was outside the WT reference range; (ii) by using the Wilcoxon rank sum statistical test to analyze all individual phenotype measurements rather than only mean values, with a Bonferroni multiple-testing correction for the effective number of tests (Supplementary Table 5); and (iii) by calculation of Mahalanobis distances to ensure that lines with significantly abnormal phenotypes were not overlooked when they resulted from simultaneous but smaller variances in multiple phenotype parameters. 17 lines (34%) had at least one abnormal phenotype parameter outside the reference range, 12 (24%) of which had increased cartilage surface damage. 18 lines (36%) were outliers after statistical analysis using the Wilcoxon test with Bonferroni correction, and 10 (20%) were outliers following Mahalanobis analysis. Overall, 25 individual lines (50%) had an abnormal joint phenotype based on these criteria (Figure 1).

The effect of gene deletion on phenotype severity was assessed by scoring whether joint abnormalities were (i) reference range outliers, (ii) outliers based on the Bonferroni-corrected Wilcoxon test, (iii) outliers after Mahalanobis analysis, and by considering whether joint pathology included abnormalities of cartilage morphology, cartilage integrity and/or subchondral bone structure (Supplementary Table 6). Of the 25 lines with an outlier phenotype, the top seven ranked lines with the most severe abnormalities were a cluster of 6 microRNAs (*ClusterXN1: miR-106a, miR-18b, miR-19b-2, miR-20b, miR-92-2, miR-363*), paired like homeodomain 1 (*Pitx1*), fibroblast growth factor binding protein 1 (*Fgfbp1*), Josephin domain containing 1 (*Josd1*), SH3 domain binding protein 4 (*Sh3bp4*), unkempt family zinc finger (*Unk*), and zinc finger protein 341 (*Zfp341*) (Figure 1, Supplementary Table 6).

### Prioritization of 25 mouse mutants with outlier joint phenotypes

An informatics strategy was used to investigate biological plausibility and prioritize the 25 candidate genes based on (i) additional skeletal consequences of gene deletion, (ii) gene expression in skeletal cells and tissues, (iii) association with human disease, and (iv) structured literature searching. Each criterion was assigned a score and scores were summed to rank genes in order of priority.

Additional skeletal consequences of gene deletion were investigated by: (i) identifying whether mutant mice had abnormalities of bone structure and strength^16^, (ii) IMPC IMPReSS phenotype screening, (iii) analysis of the skeleton in embryos from lines in which homozygous gene deletion was lethal or resulted in sub-viability^25^, and (iv) determining whether mutations of the same gene had skeletal abnormalities identified in Mouse Genome Informatics (MGI) databases or the published literature. The top ranked genes with major effects on skeletal phenotype were *Pitx1*, basic helix-loop-helix family member e40 (*Bhlhe40*), SR-related CTD associated factor 11 (*Scaf11*) and SMG9 nonsense mediated mRNA decay factor (*Smg9*) (Figure 1, Supplementary Table 6).

Gene expression in skeletal cells and tissues was investigated by interrogation of (i) MGI and BioGPS^26^ databases, and (ii) transcriptome datasets from human chondrocytes and cartilage^27, 28^, mouse and human osteoblasts^26, 28^, mouse osteocytes^19^, and mouse and human osteoclasts^26, 28^. The top ranked genes with expression enriched in cartilage and bone compared to non-skeletal tissues were *Bhlhe40*, *Pitx1*, collagen type IV alpha 2 chain (*Col4a2*) and *Sh3bp4* (Figure 1, Supplementary Table 6).

Association with human disease was investigated by searching MGI and Online Mendelian Inheritance in Man (OMIM) databases to determine whether mutations in any of the 25 genes resulted in monogenic diseases affecting the skeleton. To determine whether any gene loci were associated with arthritis or other skeletal phenotypes, we interrogated the European Bioinformatics Institute GWAS catalogue. The top ranked genes associated with arthritis and skeletal disease in humans were *Pitx1*, coiled-coil domain containing 6 (*Ccdc6*), HECT and RLD domain containing E3 ubiquitin protein ligase family member 1 (*Herc1*), nebulette (*Nebl*)*, Sh3bp4* and *Zfp341* (Figure 1, Supplementary Table 6).

Structured literature searching of PubMed and Google Scholar databases identified *Bhlhe40*, *Col4a2* and *Pitx1* as the top ranked candidate genes with publications related to arthritis or skeletal cell biology (Figure 1, Supplementary Table 6).

Together, consideration of additional skeletal phenotypes, gene expression, association with human disease and the published literature prioritized 10 lines (*Pitx1*, *Bhlhe40*, *Scaf11*, *Smg9*, *Col4a2*, *Sh3bp4*, *Ccdc6*, *Herc1, Nebl* and *Zfp341*). 11 (44%) of the 25 unselected candidates with outlier phenotypes (*Arhgap30*, *Arrdc5*, *ClusterXN1*, *Cpgi81*, *Gsdme*, *Hpf1*, *Josd1*, *Mkrn2*, *Scaf11*, *Slamf9*, *Smg9*) had no prior links to osteoarthritis or skeletal biology based on association with human disease and structured literature searching, whereas only two (20%) of the 10 prioritized candidates (*Scaf11*, *Smg9*) had no prior association (Figure 1, Supplementary Table 6).

Combining findings from informatics prioritization with the seven lines with the most severe joint phenotypes (*ClusterXN1*, *Pitx1*, *Fgfbp1*, *Josd1*, *Sh3bp4*, *Unk*, *Zfp341*) identified 14 genes with strong evidence from independent sources for a functional role in osteoarthritis pathogenesis (*Pitx1*, *Bhlhe40*, *Scaf11*, *Smg9*, *Col4a2*, *Sh3bp4*, *Ccdc6*, *Herc1, Nebl*, *Zfp341, ClusterXN1*, *Fgfbp1*, *Josd1*, and *Unk*). The four genes with the strongest overall evidence were *Pitx1*, *Bhlhe40*, *Sh3bp4* and *Unk* (Figure 1, Supplementary Table 6).

### Severe early onset osteoarthritis in heterozygous Pitx1^+/-^ mice

*Pitx1* is a homeobox transcription factor required for patterning, and specification of hindlimb morphology^29, 30^. Homozygous *Pitx1^-/-^* mutations cause post-natal lethality with gross morphological abnormalities affecting the hindlimb skeleton^29, 30^. Genomic rearrangements at the human *PITX1* locus result in homeotic arm-to-leg transformations in Liebenberg syndrome (OMIM 186550), and misexpression of *Pitx1* in the mouse forelimb recapitulates this phenotype^31^. Haploinsufficiency for *Pitx1* in mice and humans results in clubfoot and other leg malformations, demonstrating hindlimb development is sensitive to *Pitx1* gene dosage^32^.

16-week-old heterozygous *Pitx1^+/-^* mice had the most severe phenotype observed in this study, with extensive joint damage affecting both compartments of the knee including decreased articular cartilage volume and thickness, and increased areas of complete loss or severe erosion of articular cartilage. The *Pitx1^+/-^* phenotype was as severe as the extensive joint damage observed in 22-week-old WT mice 12 weeks after DMM surgery (Figure 3 and Supplementary Figure 2A and B). Accordingly, histology confirmed severe osteoarthritis with extensive vertical clefts in articular cartilage and areas of erosion over 50-75% of the articular surface to the calcified cartilage beneath (OARSI grade 5) (Figure 3, Supplementary Figure 2C and Supplementary Table 3).

### Early onset osteoarthritis in Bhlhe40^-/-^ and Sh3pb4^-/-^ mutant mice

*Bhlhe40^-/-^* mice had decreased median and maximum AC Th affecting both compartments of the knee. Further evidence of spontaneous damage included vertical clefts in the articular cartilage surface extending down to the layer of chondrocytes immediately below the superficial layer, together with some loss of surface lamina (OARSI grade 2) (Figure 4, Supplementary Figure 4 and Supplementary Table 3). *Bhlhe40* is a widely expressed transcription factor involved in regulation of cell proliferation, differentiation, apoptosis and senescence^33^. *Bhlhe40* expression is enriched in skeletal tissues (Supplementary Table 6), specifically in proliferating and differentiating chondrocytes during endochondral ossification^34^, and is elevated in response to hypoxia^35^, bone morphogenetic protein-2 (BMP2) and transforming growth factor-β (TGFβ) but suppressed by parathyroid hormone^36^. *Bhlhe40* stimulates terminal chondrocyte and osteoblast differentiation^36, 37^, and has been implicated in bone loss in chronic periodontitis^38^. Deletion of *Bhlhe40* in mice results in increased bone mineral content and density (Supplementary Table 6), but a role in osteoarthritis pathogenesis has not been postulated and *BHLHE40* has not been associated with osteoarthritis in GWAS. Overall, young adult *Bhlhe40^-/-^* mice display thinning of articular cartilage, suggesting a novel role in disease onset that involves the key hypoxia, BMP2, TGFβ and PTH signalling pathways^39–42^.

**Figure 4.**
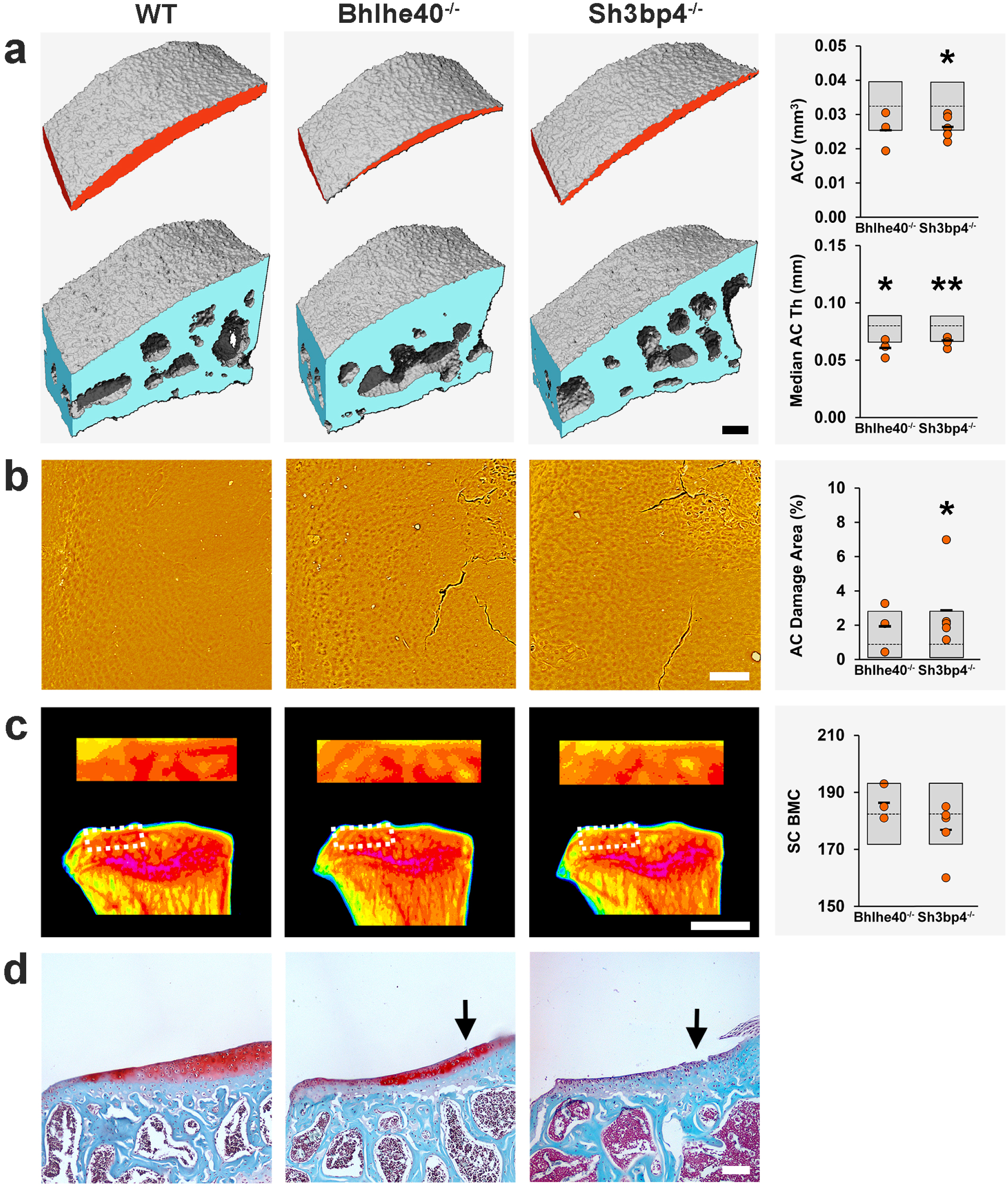
Early onset *osteoarthritis in Bhlhe40^-/-^ and Sh3pb4^-/-^ mutant mice*. ***a.*** Iodine contrast enhanced μCT images of lateral tibial plateau articular cartilage and subchondral bone from WT and homozygous *Bhlhe40^-/-^* and *Sh3bp4^-/-^* mice. Graphs show articular cartilage volume (ACV) and median articular cartilage thickness (Median AC Th) in *Bhlhe40^-/-^* (n=3) and *Sh3bp4^-/-^* (n=5) mice. ***b.*** Back-scattered electron-scanning electron microscopy images of lateral tibial plateau joint surface replicas from WT, *Bhlhe40^-/-^* and *Sh3bp4^-/-^* mice. Graph shows articular cartilage damage. ***c.*** Pseudocoloured X-ray microradiography images of proximal tibia and the lateral tibial plateau subchondral bone region of interest (dashed box) from WT, *Bhlhe40^-/-^* and *Sh3bp4^-/-^* mice. Low bone mineral content (BMC) is yellow and high BMC is pink. Graph shows no change in subchondral BMC (SC BMC). ***d.*** Coronal sections of lateral knee joint compartment stained with Safranin-O/Fast green from WT, *Bhlhe40^-/-^* and *Sh3bp4^-/-^* mice. Arrows indicate areas of cartilage damage. Grey boxes, reference ranges; dashed lines, population means/medians; orange circles, individual mutant samples; and black horizontal lines, sample means. Scale bars = 100μm (A, B, D) and 1mm (C). \**P*<0.00568, \*\**P*<0.001.

*Sh3bp4^-/-^* mice had decreased ACV and median and maximum AC Th, together with increased articular cartilage surface damage affecting the lateral compartment. Histology revealed moderate osteoarthritis with articular cartilage fibrillations and vertical clefts over <25% of the articular surface extending to the calcified cartilage beneath (OARSI grade 3) (Figure 4, Supplementary Figure 4 and Supplementary Table 3). *Sh3bp4* is a poorly characterized SH3 domain binding protein involved in transferrin receptor (TfR) internalization^43^, fibroblast growth factor receptor (FGFR) trafficking^44^, mammalian target of rapamycin (mTOR) signalling^45^ and inhibition of the Wnt pathway^46^. *SH3BP4* has not been associated with osteoarthritis in GWAS.

In summary, *Sh3bp4^-/-^* mice display loss of articular cartilage and moderate cartilage damage. *Sh3bp4* thus represents a new osteoarthritis susceptibility gene, the deletion of which accelerates joint damage. Its role in disease pathogenesis may involve the key TfR, FGFR, mTOR and Wnt signalling pathways, which regulate bone and cartilage homeostasis and tissue repair^47–50^.

### Additional applications of the OBCD joint phenotyping pipeline

The development of new joint phenotyping methods to investigate unselected mutant mice demonstrates the power of unbiased functional genomics in osteoarthritis gene discovery. Open access availability of IMPC mouse lines provides a rich resource for investigation of disease mechanisms and the identification and testing of new preventagoldtive or disease modifying drugs. Here we demonstrate how the OBCD joint phenotype database can be leveraged to add value to studies of human osteoarthritis and how the new imaging techniques can be applied to address additional hypotheses.

### Application 1: Early onset osteoarthritis in mice with deletion of genes differentially expressed in human osteoarthritis cartilage

Phenotyping genetically modified mouse models is a powerful method to functionally annotate potential disease susceptibility genes identified in human studies. We interrogated the OBCD database of joint phenotypes from unselected mouse lines to annotate the function of 409 genes differentially expressed in low-versus high-grade articular cartilage in osteoarthritis patients^51^. Phenotype data were available from eight differentially expressed genes (*Unk*, *Josd1,* gasdermin E (*Gsdme*), Rho GTPase activating protein 30 (*Arhgap30*)*, Ccdc6, Col4a2,* methyl-CpG binding domain protein 1 (*Mbd1*) and staufen double-stranded RNA binding protein 2 (*Stau2*)). Mutation of six (75%) differentially expressed genes in mice resulted in joint abnormalities whereas deletion of the other two (25%) (*Mbd1*, *Stau2*) had no effect (Supplementary Tables 3 and 4). This compares with 14 out of 50 unselected lines (28%) with strong evidence for a functional role in osteoarthritis following prioritization in this study (*P*=0.01582, Fisher’s exact test). This enrichment of signal supports interrogation of the OBCD phenotype database to accelerate functional gene discovery in osteoarthritis. Combining complementary mouse and human gene discovery approaches demonstrates synergy, especially as none of the genes identified have been associated with osteoarthritis in GWAS.

*Unk^-/-^* mice had decreased articular cartilage volume with increased surface damage affecting the MTP and decreased maximum cartilage thickness in the LTP (Figure 5, Supplementary Figure 5, Supplementary Tables 3, 4 and 6). UNK is an RNA binding zinc finger protein expressed in developing brain that controls neuronal morphology^52^. *UNK* is also expressed in the skeleton (Supplementary Table 6). *Unk^-/-^* mice display articular cartilage loss with early cartilage damage indicating *Unk* is a protective osteoarthritis susceptibility gene. Downregulation of *UNK* at the RNA and protein levels in high-grade osteoarthritis cartilage in humans suggests a role in disease pathogenesis.

**Figure 5.**
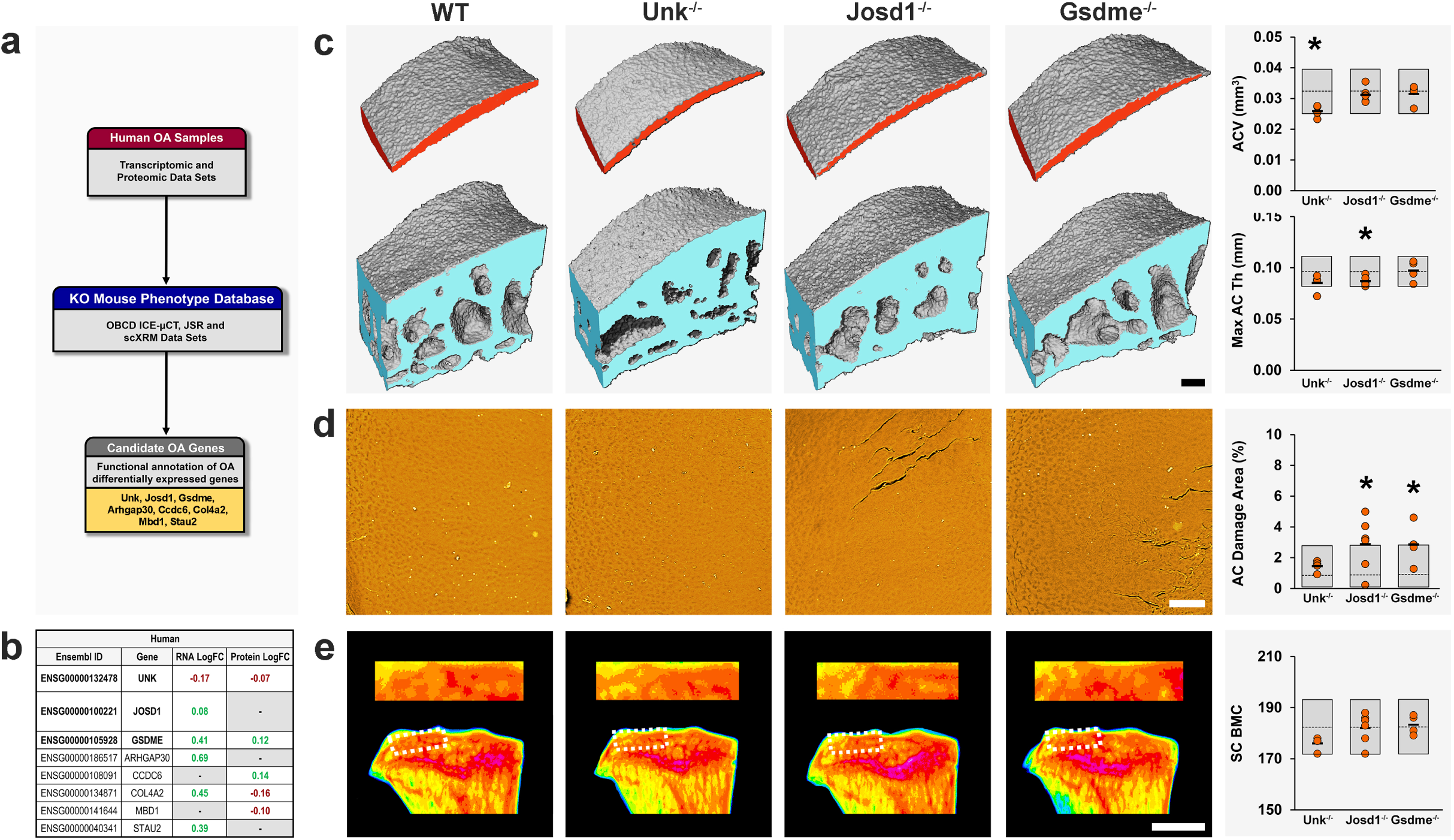
Application 1: Early onset osteoarthritis in mice with deletion of genes differentially expressed in human osteoarthritis cartilage. ***a.*** Mouse lines with deletion of candidate genes differentially expressed in osteoarthritic human articular cartilage (Unk, Josd1, Gsdme, Arhgap30, Ccdc6, Col4a2, Mdb1 and Stau2) were investigated for osteoarthritis phenotypes. ***b.*** Differential expression (log fold change, logFC) of human orthologues of candidate genes in low-grade versus high-grade osteoarthritis in human articular cartilage samples. Genes with significantly increased expression in high-grade osteoarthritis cartilage shown in green and those with decreased expression in red. ***c.*** Iodine contrast enhanced μCT images of lateral tibial plateau articular cartilage and subchondral bone from WT, Unk^-/-^, Josd1^-/-^ and Gsdme^-/-^ mice. Graphs show articular cartilage volume (ACV) and maximum articular cartilage thickness (AC Max Th) in Unk ^-/-^ (n=4), Josd1 ^-/-^ (n=6) and Gsdme^-/-^ (n=4) mice. ***d.*** Back-scattered electron-scanning electron microscopy images of lateral tibial plateau joint surface replicas from WT, Unk^-/-^, Josd1^-/-^ and Gsdme^-/-^ mice. Graph shows articular cartilage damage. ***e.*** Pseudocoloured X-ray microradiography images of proximal tibia and the lateral tibial plateau subchondral bone region of interest (dashed box) from WT, Unk^-/-^, Josd1^-/-^ and Gsdme^-/-^ mice. Low bone mineral content (BMC) is yellow and high BMC is pink. Graph shows subchondral BMC (SC BMC). Grey boxes, reference ranges; dashed lines, population means/medians; orange circles, individual mutant samples; and black horizontal lines, sample means. Scale bars = 100μm (C, D) and 1mm (E). *P<0.00568.

*Josd1^-/-^* mice had decreased median and maximum cartilage thickness affecting both joint compartments, and increased articular cartilage surface damage affecting the lateral compartment (Figure 5, Supplementary Figure 5, Supplementary Tables 3, 4 and 6). JOSD1 is a widely expressed deubiquitinating enzyme that may play a role in regulation of cell membrane dynamics^53^. It stabilizes SOCS1, an important negative regulator of cytokine signalling^54, 55^, and *JOSD1* mRNA is upregulated in high-grade osteoarthritis cartilage (Figure 5), suggesting a protective role for JOSD1 in osteoarthritis.

*Gsdme^-/-^* mice had increased articular cartilage surface damage affecting the lateral compartment (Figure 5, Supplementary Figure 5, Supplementary Tables 3, 4 and 6). Gasdermin E is a member of a family of proteins that facilitate necrotic programmed cell death (pyroptosis) following cleavage by caspase-3^56^. Pyroptosis has recently been shown to promote knee osteoarthritis^57, 58^, and upregulation of *Gsdme* in high-grade osteoarthritis cartilage (Figure 5), supports a role for gasdermin E in disease progression.

*Arhgap30^-/-^* mice had increased articular cartilage surface damage affecting the lateral compartment and a significant outlier phenotype following Mahalanobis analysis (Supplementary Tables 3, 4 and 6). *Arhgap30* encodes a Rho GTPase implicated in cell proliferation, migration and invasion acting via inhibition of Wnt^59^. Its putative role and increased expression in high-grade osteoarthritis cartilage, suggests involvement of ARHGAP30 in cartilage repair mechanisms during osteoarthritis pathogenesis.

*Ccdc6^-/-^* mice had increased articular cartilage surface damage affecting the medial compartment (Supplementary Tables 3, 4 and 6). *Ccdc6* is a cell cycle checkpoint regulator that facilitates cell survival^60^. *CCDC6* was associated with heel BMD in a large GWAS^19^ and *Ccdc6^-/-^* mice have decreased bone strength (Supplementary Table 6). Concordant with upregulation of CCDC6 in high-grade osteoarthritic cartilage (Figure 5), *CCDC6* was differentially expressed in a meta-analysis of gene expression profiling in synovial tissue from osteoarthritis cases and controls^61^. Overall, *CCDC6* represents a new osteoarthritis susceptibility gene, deletion of which may contribute to development of osteoarthritis by actions in both cartilage and the synovium.

*Col4a2^-/-^* mice had increased articular cartilage surface damage affecting the lateral compartment (Supplementary Table 3). The collagen type IV alpha 2 chain is a major component of vascular basement membrane^62^, but is also present in skeletal basement membranes due to expression in bone cells (Supplementary Table 6). *COL4A2* was differentially expressed in osteoarthritis synovium^61^, in knee joints following DMM surgery^63^, in a time course analysis following DMM surgery^64^, and in articular cartilage biopsies from osteoarthritis patients^65^. The *COL4A2* locus was differentially methylated in a genome-wide analysis of hip compared to knee osteoarthritis cartilage^66^. In summary, several lines of evidence suggest a role for *Col4a2* in the pathogenesis of osteoarthritis and the articular cartilage damage in *Col4a2^-/-^* mice is consistent with this hypothesis (Figure 1 and Supplementary Table 3).

### Application 2: Age-related joint degeneration

We next studied 4- and 12-month-old WT mice to investigate the effect of ageing. Joints from 12-month-old mice had features of osteoarthritis compared to 4-month-old juvenile mice. Even though articular volume and thickness did not change with age, the 12-month-old mice had significantly increased areas of cartilage damage in both lateral and medial compartments. Histology revealed an increased maximum OARSI score on the LTP in 12-month-old compared with 4-month-old mice. These changes were accompanied by loss of subchondral bone (decreased SC BV/TV, SC Tb.Th and SC Tb.N) in the lateral compartment and increased SC BMC in the medial compartment (Figure 6 and Supplementary Figure 6).

**Figure 6.**
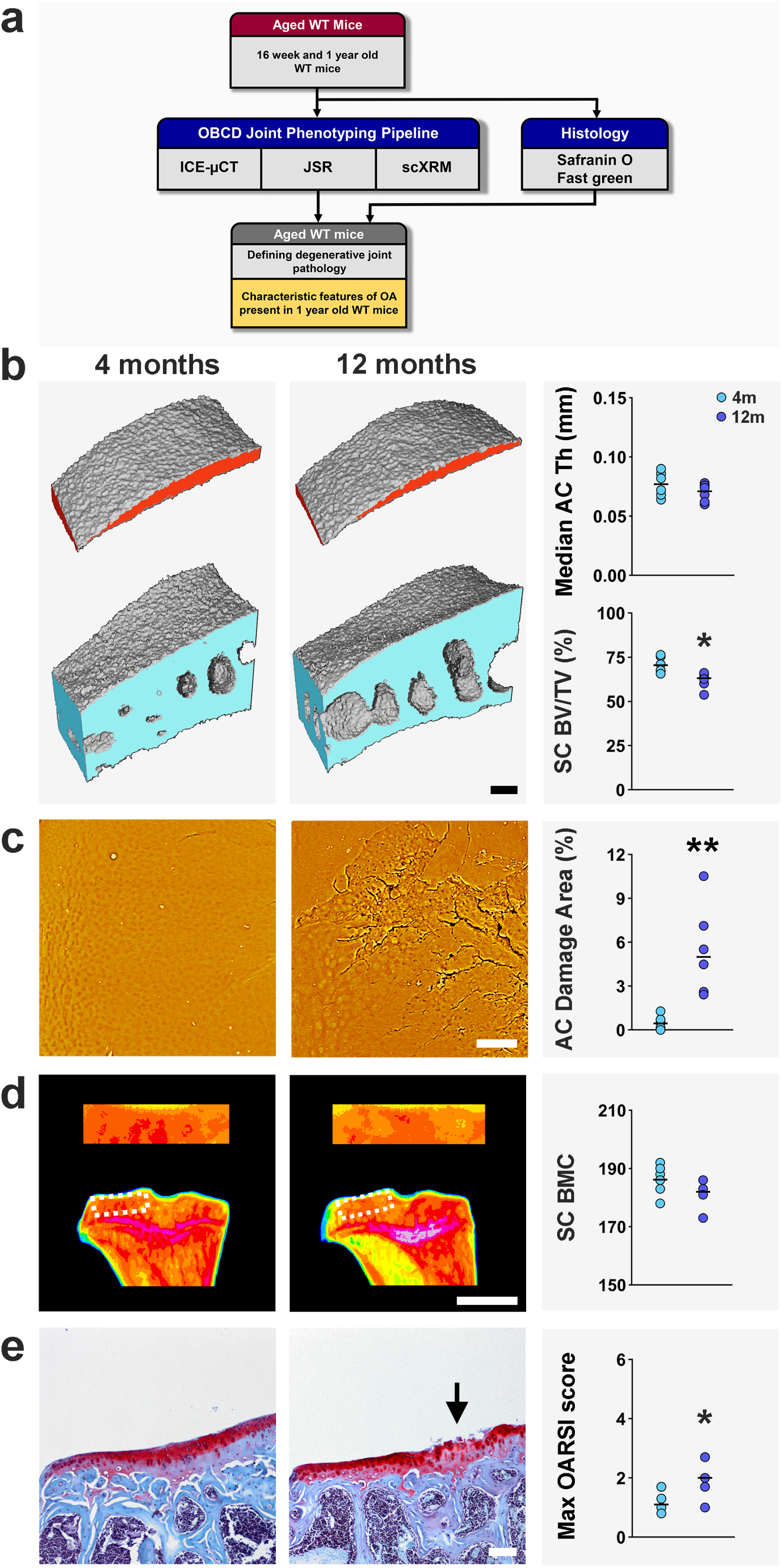
Application *2: Age-related joint degeneration*. ***a.*** Cohorts (n=6) of young adult (4 months) and aged (12 months) WT mice were analyzed by iodine contrast enhanced μCT, (ICEμCT), joint surface replication (JSR) and subchondral X-ray microradiography (scXRM). ***b.*** ICEμCT images of lateral tibial plateau articular cartilage and subchondral bone. Graphs show median articular cartilage thickness (Median AC Th) and subchondral bone BV/TV (SC BV/TV). ***c.*** Back-scattered electron-scanning electron microscopy images of lateral tibial plateau JSRs. Graph shows articular cartilage damage. ***d.*** Pseudocoloured scXRM images of proximal tibia and the lateral tibial plateau subchondral bone region of interest (dashed box). Low bone mineral content (BMC) is yellow and high BMC is pink. Graph shows subchondral BMC (SC BMC). ***e.*** Coronal sections of lateral knee joint compartment stained with Safranin-O/Fast green. Graph shows maximum OARSI histological scores. Arrow indicates cartilage damage. Scale bars = 100μm (B, C, E) and 1mm (D). \**P*<0.05, \*\**P*<0.01.

### Application 3: Mice with a Dio2^Ala92^ polymorphism are protected from osteoarthritis

As proof-of-concept for functional investigation of signals arising from human genetic association studies, we studied the effect of rs225014, a polymorphism in the human *DIO2* gene. rs225014 results in a substitution at amino acid 92 (Thr92Ala) in the DIO2 enzyme that activates thyroid hormones in target cells. The minor allele (Ala92) frequency is estimated at 40%^67^. rs225014 was positively associated with osteoarthritis in a genome-wide linkage study that included replication in a separate cohort^68^, but this association was not reproduced in the Rotterdam study ^69^ or a later meta-analysis^70^. Nevertheless, a polymorphism in the *DIO2* promoter (rs12885300) has been associated with hip geometry in genome-wide linkage studies. While minor alleles at rs225014 and rs12885300 were associated with opposite osteoarthritis outcomes, a haplotype combining both was associated with increased disease susceptibility^71^. Overall, these findings implicate regulation of local thyroid hormone availability in the pathogenesis of osteoarthritis and are consistent with well-known actions of thyroid hormones, which stimulate hypertrophic chondrocyte differentiation^72^ and expression of cartilage matrix degrading enzymes^73^. Nevertheless, *DIO2* has not been associated with osteoarthritis in GWAS, and the role of the *DIO2* rs225014 polymorphism remains controversial, attracting much debate even beyond the osteoarthritis field^74–76^.

*In vivo* studies have demonstrated increased subchondral bone but intact articular cartilage in *Dio2^-/-^* knockout mice^77^. *Dio2^-/-^* mice have decreased calreticulin expression in articular cartilage^78^, a gene implicated in cartilage thinning in response to mechanical loading^79^, and are protected from cartilage damage following forced exercise. By contrast, increased *DIO2* expression resulted in negative effects on chondrocyte function and homeostasis *in vitro*^80^ and *DIO2* expression was increased in articular cartilage from osteoarthritis patients^81^. Furthermore, cartilage-specific overexpression of *Dio2* in transgenic rats resulted in cartilage destruction^82^. Together, these studies suggest that decreased DIO2 expression and reduced thyroid hormone availability in the joint may protect against osteoarthritis, whereas increased DIO2 expression may increase susceptibility.

To test this hypothesis, we used the CRISPR/Cas9 system to generate *Dio2^Thr92^* and *Dio2^Ala92^* mutant mice. The Thr92Ala polymorphism results in decreased enzyme activity and impaired conversion of the prohormone thyroxine (T4) to the active hormone triiodothyronine (T3) reducing local thyroid hormone signalling^74^. Analysis of joints from 16-week-old male mice demonstrated features of early onset osteoarthritis in *Dio2^Thr92^* mice compared to *Dio2^Ala92^* mice. *Dio2^Thr92^* mice had decreased cartilage volume and median thickness with increased articular cartilage damage (Figure 7 and Supplementary Figure 7). By contrast, *Dio2^Ala92^* mutants had no signs of osteoarthritis, indicating a protective role for the Ala92 polymorphism and providing the first functional evidence of a role for this candidate *DIO2* polymorphism *in vivo*. The data provide powerful evidence that decreased thyroid hormone signalling is protective against osteoarthritis.

**Figure 7.**
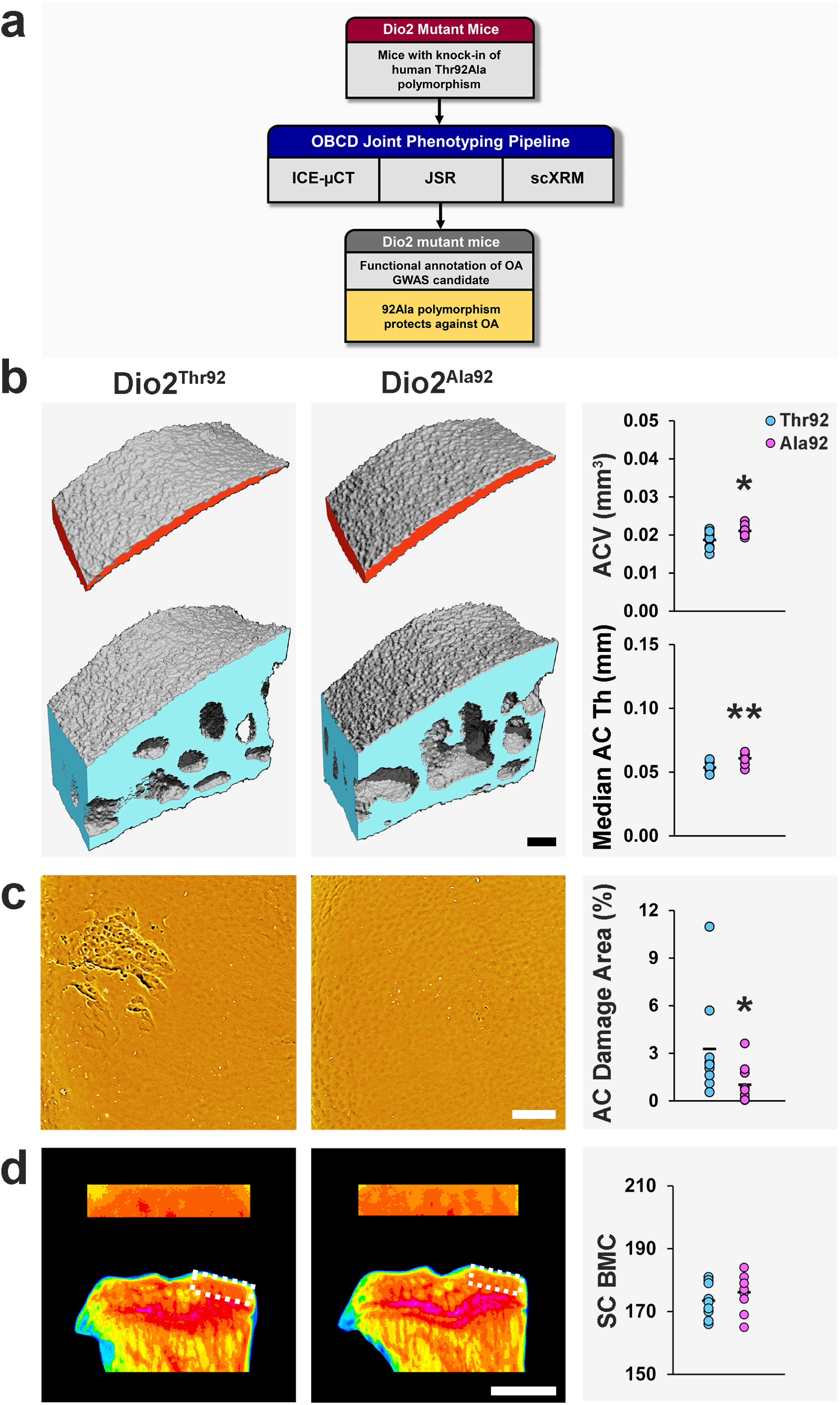
Application 3: Mice with a Dio2^Ala92^ polymorphism are protected from osteoarthritis. ***a.*** Mice with *Dio2* polymorphisms homologous to the Thr92Ala polymorphism in human *DIO2* (*Dio2^Thr92^,* n=9 and *Dio2^Ala92^*, n=10) were analyzed by iodine contrast enhanced μCT, (ICEμCT), joint surface replication (JSR) and subchondral X-ray microradiography (scXRM). ***b.*** ICEμCT images of medial tibial plateau articular cartilage and subchondral bone. Graphs show articular cartilage volume (ACV) and median articular cartilage thickness (Median AC Th). ***c.*** Back-scattered electron-scanning electron microscopy images of medial tibial plateau JSRs. Graph shows articular cartilage damage. ***d.*** Pseudocoloured scXRM images of proximal tibia and the medial tibial plateau subchondral bone region of interest (dashed box). Low bone mineral content (BMC) is yellow and high BMC is pink. Graph shows subchondral BMC (SC BMC). Scale bars = 100μm (B, C) and 1mm (D). \**P*<0.05, \*\**P*<0.01.

## Discussion

The molecular mechanisms that initiate and drive osteoarthritis progression and their genetic origins are largely unknown. We developed rapid-throughput joint phenotyping methods to identify abnormal joint phenotypes in mutant mice, identifying 14 genes with strong functional evidence for involvement in osteoarthritis pathogenesis.

*Pitx1^+/-^* mice had the most severe and extensive joint destruction in these studies, with severe erosions and complete loss of articular cartilage. Decreased *PITX1* mRNA expression in primary human articular chondrocytes, reduced PITX1 protein in histological sections of human osteoarthritic cartilage, and increased subchondral bone thickening in preliminary studies of ageing *Pitx1^+/-^* mice are all consistent with an important role for PITX1 in osteoarthritis pathogenesis^83^. Overall, we show that heterozygous deletion of *Pitx1,* a critical developmental gene, results in severe early onset joint damage. *Pitx1* is thus a major new osteoarthritis gene with a key role in disease onset and progression that likely involves its effects on joint morphology and the developmental programmes that are re-initiated during disease progression.

We exploited our joint phenotype database to characterize to functions of six candidate genes differentially expressed in human osteoarthritis cartilage. These studies identify novel roles for *Unk*, *Josd1*, *Gsdme*, *Arhgap30, Ccdc6* and *Col4a2* in the pathogenesis of osteoarthritis. *Unk^-/-^* mice were also prioritized as one of four lines with the strongest combined evidence for a key functional role in osteoarthritis pathogenesis, while *Josd1^-/-^* mice were one of seven lines with the most severely abnormal joint phenotype (Figure 1 and Supplementary Table 6). Overall, these findings demonstrate the value of leveraging the combined power of mouse and human experimental pipelines to enhance understanding of complex disease.

Previous histological studies have only demonstrated the onset of osteoarthritis at 15 months of age in wild type *C57BL6* mice^84^. The current studies, however, clearly define early features of age-related joint degeneration in one-year old animals. Thus, our new phenotyping methods are more economical and have improved sensitivity to detect early osteoarthritis compared to current labour- and resource-intensive histological techniques. In addition to identifying genes that provoke early onset of osteoarthritis when deleted, these new methods now provide efficient opportunities to analyze genes that protect joints from age-related degeneration when inhibited or deleted, or for analysis of drug intervention studies.

Finally, we identified a novel protective role for a minor allele of the *DIO2* gene (Ala92). The Ala92 variant impairs conversion of the prohormone T4 to the active hormone T3 and is controversially associated with osteoarthritis. Furthermore, it has been suggested that expression of this minor allele underlies the continuing psychological and metabolic symptoms experienced by some hypothyroid patients despite restoration of normal serum T4 levels following levothyroxine replacement^76, 85^. As a result, genetic testing and treatment of hypothyroid patients with the Ala92 variant using increased doses of levothyroxine (T4) or a combination of levothyroxine and liothyronine (T3) has been proposed. Importantly, this therapeutic approach, which aims to overcome impaired conversion of T4 to T3 by the Ala92 variant, may actually have detrimental long-term consequences as our data indicate that expression of the Ala92 variant may actually help to maintain cartilage integrity. Thus, the allelic imbalance that has been reported to increases expression of the Ala92 variant in human osteoarthritis cartilage^81^, may in fact represent a counter-regulatory response in damaged articular cartilage to protect against osteoarthritis progression. Our findings underscore the importance of caution and further research in this controversial area, and have important public health implications, as levothyroxine is the most commonly prescribed drug in the USA and third most commonly prescribed in the UK^86^.

In summary, we have shown our novel joint phenotyping methods have broad applications in osteoarthritis research that will accelerate functional gene discovery, transform understanding of disease pathogenesis and identify unanticipated drug targets for this debilitating chronic disease.

## Acknowledgements

We thank Jayashree Bagchi-Chakraborty and Mahrokh Nodani for technical assistance. We thank members of Sanger Mouse Pipelines (Mouse Informatics, Molecular Technologies, Genome Engineering Technologies, Mouse Production Team, Mouse Phenotyping) and the Research Support Facility for provision and management of mice. This work was funded by a Wellcome Trust Strategic Award (101123). Generation of mutant mice was funded by the Wellcome Trust (098051). G.R.W., J.H.D.B. were funded by a Wellcome Trust Joint Investigator Award (110140, 110141) and European Commission Horizon 2020 Grant (666869), E.Z. by the Wellcome Trust (206194), and PIC by Mrs. Janice Gibson and the Ernest Heine Family Foundation.

## Author Contributions

AB, EZ, PIC, GRW and JHDB conceived and designed experiments, NCB, VDL, JGL, JAW, AB, GRW and JHDB developed methods, NCB, JGL and JHDB wrote software code, NCB, JS, RJ, GRW and JHDB performed statistical analyses, NCB, KFC, HD, DKE, NSM, AA, VDL, JGL, JAW, EG, LS and EAM conducted experiments, JS, LS, SEY, JMW, VEV, FK, JKW, DJA, CJL, ACB, PIC, GRW and JHDB provided experimental resources, VDL curated data, AB, EZ, PIC, GRW and JHDB acquired funding, NCB, GRW and JHDB produced figures, NCB, GRW and JHDB wrote the manuscript, and all authors reviewed and edited the manuscript.

## Methods

### Experimental models

Mouse lines generated in this study have been deposited in the European Mutant Mouse Archive (EMMA; https://www.infrafrontier.eu/) or are available on request. Animal experiments were performed and reported in accordance with ARRIVE guidelines^87^. All animal experiments were conducted under licence at Imperial College (project licence PPL70/8785) and the Wellcome Trust Sanger Institute (project licence P77453634 and PPL80/2485) in accordance with the Animals (Scientific Procedures) Act 1986 and recommendations of the Weatherall report. Animal experiments were approved by the Sanger or Imperial College Hammersmith Campus Animal Welfare Ethical Review Bodies (AWERB) as appropriate. Studies performed at the University of Chicago were approved by the Institutional Animal Care and Use Committee (IACUC) at Rush University Medical Center (16-077 and 15-033).

#### IMPC wild-type and mutant mouse strains

Samples from 16-week old male wild-type (WT) and genetically-modified mice designed with deletion alleles on the *C57BL/6N Taconic;C57BL/6N* background were generated as part of the Wellcome Trust Sanger Institute’s (WTSI) Mouse Genetics Project, part of the International Mouse Phenotyping Consortium (IMPC; http://www.mousephenotype.org). Details on back-crossing status, weight, health status, administered drug and procedures, husbandry and specific conditions (including housing, food, temperature and cage conditions) have been reported previously^88, 89^ with the exception that the diet used was Mouse Breeder Diet 5021 (Labdiet, London, UK) and the IMPReSS screen had 4 fewer tests (hair phenotyping, open field, hot plate and stress induced hypothermia). Further details available on request. Mouse strains, genotypes and Research Resource Identifiers (RRIDs) are included in Supplementary Table 7.

#### Wild-type male mice for provocation studies

Mice were housed at Imperial College London in individually ventilated cages (Techniplast, UK) on a 12h light/dark cycle at 22°C. Mice were housed in sibling groups and given access to chow diet and water *ad libitum*. Mice were weighed weekly and were within a range of 25– 35g. Housing density was between 1–5 animals per cage (up to 25g) or between 1-4 animals per cage (over 25g). Health status was monitored by screening sentinel mice every 3 months, and health reports are available on request. No previous drugs, tests or procedures were administered except those specified, and no adverse events occurred. Mice were euthanized by exposure to increasing concentrations ofCO_2_ followed by cervical dislocation.

For surgical provocation of osteoarthritis, WT *C57BL/6* virgin male mice were purchased from Charles River Laboratories (Margate, Kent, UK; n=32). Mice had been maintained as homozygous WT and back-crossed more than 20 generations (https://www.criver.com/sites/default/files/Technical%20Resources/C57BL_6%20Mouse%20Model%20Information%20Sheet.pdf). Mice were housed 4 per cage in sibling groups with identical enrichment and allocated to surgical groups based on cage number. Weight was monitored before and for 48 hours after surgery. Order of surgery was based on cage number and numerical mouse ID, and is available on request. Surgery was performed at 10 weeks of age.

For provocation of osteoarthritis by aging, WT *129/SV/C57BL/6J* male mice (16 weeks and 1 year of age, 12 total, 6 mice/group) were generated from a long-standing colony originally obtained from Jacques Samarut (École Normale Supérieure, Lyon, France)^90^. WT mice were maintained and back-crossed for more than 20 generations.

#### Type 2 iodothyronine deiodinase (Dio2) mutant mice

Homozygous 16-week-old F3 male mice (19 mice total) harboring alanine/alanine or threonine/threonine polymorphisms for the deiodinase 2 gene at codon 92 (*Dio2^Ala92^*, n=9 and *Dio2^Thr92^, n=10*) were generated by Applied Stemcell Inc. (Milpitas, CA, USA) and housed as described^74^. No previous drugs, tests or procedures were administered. Information regarding back-crossing status, weight, health status, husbandry and specific conditions (including housing and cage conditions) is available on request.

#### Human sample collection

Tissue samples were collected from 115 patients undergoing total joint replacement surgery (102 knee and 13 hip osteoarthritis, see Steinberg *et al*.,^51^ accompanying related manuscript) for ethical compliance and details of the committee that approved the study design,. All patients provided written, informed consent prior to participation. Matched low-grade (healthy tissue or low-grade degeneration) and high-grade (high-grade degeneration) cartilage samples were collected from each patient.

### Rapid-throughput joint phenotyping

#### Sample preparation

Hindlimbs from WT and mutant mice were skinned and fixed for 24-30h in 10% neutral buffered-formalin, rinsed twice and stored in 70% ethanol at 4°C. Samples were anonymized and randomly assigned to batches for rapid-throughput analysis in an unselected fashion. Prior to analysis, limbs were rehydrated in PBS + 0.02% sodium azide for >24h. Soft tissue was removed and the knee joint was disarticulated under a Leica MZ9 dissecting microscope (Leica Microsystems, UK) with the aid of fine forceps (Dumont #5, Cat#11252-20; Fine Science Tools, Germany) and 5mm spring scissors (Vannas Tubingen, Cat#15003-08, Fine Science Tools, Germany).

#### Iodine contrast-enhanced µCT (ICEµCT)

This method was developed to detect signs of OA, including cartilage damage and subchondral bone sclerosis or bone loss. The joint is immersed in an iodinated contrast agent with similar X-ray absorption to bone and imaged (Figure 2). This allows segmentation of both the articular cartilage and underlying subchondral bone in the same volume of interest. The technique can also be used to analyze the condyles and head of the femur.

Rehydrated tibiae were blotted on low-linting tissue (Kimtech, UK) to remove all liquid. Single tibiae were placed into 6mm sample holders (Scanco Medical AG, Brüttisellen, Switzerland) in 180mg/mL ethiodized oil (Lipiodol Ultra, Guerbet Laboratories, Solihull, UK) mixed with sunflower oil. The tibial epiphysis (1-1.5mm) was scanned using a Scanco µCT-50 micro-computed tomography scanner (µCT) at 70 kVA, 2µm voxel resolution with 1500ms integration time and 1x averaging.

Analysis was performed using Xming 6.9.0.31 (©2005-2007 Colin Harrison) software implemented with the pUTTy client program (release 0.62 ©1997-2011 Simon Tatham). Tomographs were rotated twice to place the limb in a standard position. First, the tibia was rotated clockwise to point the anterior-most patellar surface vertically. Secondly, the tibia was rotated 90° to place the medial-lateral axis of the tibia facing the viewer (Figure 2A, coronal views). Medial and lateral tibial plateaux were analyzed separately. Equally sized medial and lateral VOIs were scaled according to the size of the tibia (Figure 2A, medial-lateral dimensions, a-b; anterior-posterior dimensions, c-d). The width of each VOI was equal to 14% of the medial-lateral axis, which equated to 59% of the medial plateau width and 53% of the lateral plateau width. The anterior-posterior dimension of each VOI was 33% of the size of each tibia. VOIs were located at the same positions on each limb by determining the plateau edges (Figure 2A, e, f, g, h). Each VOI was then centered on the medial-lateral midpoint of the plateau (Figure 2A, dashed line), and the lateral plateau VOI was further offset by a distance equivalent to 7% of the medial-lateral axis (Figure 2A). The medial VOI midpoint was 61.5%, and the lateral VOI midpoint 68.5% (7% posterior shift) along the anterior-posterior axis. This accounted for the different shapes of the medial and lateral plateau, based on standard measurements determined from mean values obtained from 10 wild-type mice to capture the thickest part of each plateau (data not shown).

For articular cartilage, all tissue below a density threshold of 200 mg/hydroxyapatite/cm^3^ (mg HA/cm^3^) was analyzed (Figure 2B-C). The volume and thickness (median and maximum) were determined. For subchondral bone, all tissue between the articular cartilage and the growth plate above a density threshold of 300 mg/HAcm^3^ was analyzed. Subchondral bone volume/tissue volume (BV/TV), bone mineral density (BMD), trabecular thickness (Tb.Th) and trabecular number (Tb.N) were determined.

#### Joint surface replication (JSR)

This method was developed to determine articular cartilage surface damage. The technique enables visualization of joint surface morphology at high-resolution in the natural hydrated condition by using a resin cast as a surrogate (Figure 2). The method can also be used to analyze the condyles and head of the femur.

#### Joint surface replicas: Moulding

Disarticulated, rehydrated tibiae were positioned with the articular surfaces parallel to the moulding surface using Draper Helping Hand brackets with articulated arms (31324Hh, Bamford Trading, Ross-on-Wye, UK). A multi-aperture moulding template of depth >0.5 cm was custom-made. Moisture was removed to preserve surface detail by blotting with low-linting tissue (Kimtech). Samples were immersed immediately in Virtual Light Body dental impression medium (Ivoclar-Vivadent, Leicester, UK) for 4 minutes prior to de-moulding and immediate return to storage medium. Moulds were visually inspected for defects or bubbles and either cast immediately or stored for a maximum of 2 weeks according to manufacturer’s advice.

#### Joint surface replicas: Casting

Joint surface replicas were cast using Crystal Clear 202 acrylic resin (Smooth-On, Bentley Advanced Materials, London, UK) according to manufacturer’s instructions (Figure 2). Pre-warmed (37°C) parts A and B (1:9 ratio) were mixed thoroughly and vacuum de-gassed. Resin was poured into joint surface moulds by gravity flow using a fine pipette tip with the aid of a dissecting microscope (MZ9, Leica Microsystems, UK) and bubbles were removed from mould-resin interfaces. Casts were set for 16h at 18-23°C, post-cured for 6h at 65°C and removed from moulds after 7 days at room temperature, according to manufacturer’s instructions. Each cast was secured to a custom-produced 10×10×0.3cm aluminum raft with double-sided carbon tape (Agar Scientific, Stansted, UK). Rafts of samples were coated with >20nm carbon using a high-vacuum bench-top carbon coater (Agar Scientific,).

#### Joint surface replicas: Imaging

Coated samples were analyzed using a Vega3 XMU scanning electron microscope (Tescan UK, Cambridge) at high vacuum with a 4-quadrant back-scattered electron detector (Deben, Bury St. Edmunds, UK). Tibial plateaux were imaged at 1536×1536 pixel image size with a scan speed of 5 and 3x frame-corrected averaging over a view field of 1800×1800µm (Figure 2). Beam voltage was 20kV and working distance was 17mm. ImageSnapper semi-automated imaging software (V1.0, Tescan UK) was used to acquire images.

#### Joint surface replicas: Damage quantitation

A quantitation pipeline was developed to determine areas of damage on the joint surfaces of the medial and lateral plateau accurately, while excluding extraneous surface debris (Figure 2). Macros (ImageJ1.44) were produced for automated analysis (see below; Supplementary Data 1). Samples were analyzed in batches by a blinded operator. After each step, macros included batch-saving of modified images, unless otherwise stated. Particle sizes are based on pixel size of 1.37µm^2^. Steps for semi-automated damage quantitation were:

1. Select plateau (*free select* tool; Fig. 2C, 2^nd^ panel and close-up). Isolated plateaux were batch-saved with a modified file name (*Macro 1*)
2. Exclude bright outlier particles <4 pixels (5.5µm^2^; *Macro 2*)
3. Detect damage using the *find edges* tool (*Macro 3,* Figure 2C, 3^rd^ panel and close-up)
4. Manually erase debris (soft tissue, bubbles in moulded surfaces) with close comparison to the original image
5. Adjust the threshold to capture all damage accurately with close comparison to the original image (*Macro 5,* Figure 2C, 4^th^ panel and close-up)
6. Quantify damage area (*analyze particles* tool; *Macro 6,* Figure 2C, 5^th^ panel and close-up) above a particle size >20 pixels (27.5µm^2^), with a circularity 0**–**0.5 (0; line; 1; perfect circle). Excluding circular objects further refines the damage selection as circular objects are not likely to be genuine damage

Damage area was recorded and represented as percentage of the total plateau area. The whole plateau area was determined by performing steps 1–2 above, and then:

3) Set threshold to minimum (*Macro 5)*
4) Determine whole plateau area *(analyze particles* tool; *Macro 6;* plateau outline in Figure 2C, 5^th^ panel) above a particle size >100 pixels (188µm^2^), with a circularity of 0**–**1

#### Subchondral X-ray microradiography (scXRM)

Quantitative subchondral X-ray microradiography (scXRM) was performed at 10µm pixel resolution using a Faxitron MX20 variable kV point-projection X-ray source and digital image system (Qados, Cross Technologies plc, Berkshire, UK) with modifications to previously published protocols^21, 22^.

Disarticulated tibiae were imaged flat with the posterior surface facing downwards (Figure 2). Samples were imaged at 26 kV for 15 s with the sample tray raised 6mm above the 5X magnification slot to maximize analysis area. Quantitation was performed in ImageJ1.44 by stretching pixel information between density values for plastic (minimum) and steel (maximum) standards and assigning to one of 256 bins using Macro 7 (Supplementary Data 1). The width of each tibia at the growth plate was determined and used to scale the regions of interest (ROIs; height: 9%, width: 34%, of the growth plate width). These dimensions were based on an optimal ROI size determined from 10 wild-type tibiae that was sufficiently wide to include the whole subchondral plate and shallow enough to avoid calcified cartilage. To analyze mice harboring polymorphisms in the *Dio2* gene, the MTP ROI was truncated by 10% of its height to avoid the calcified cartilage. The ROI positions were parallel to each plateau and included the topmost yellow pixel representing the bone directly beneath the tibial plateau. For each ROI, the number of pixels in each bin was determined using the *Custom Histogram* macro (http://rsb.info.nih.gov/ij/). The median grey level (density bin at which the cumulative frequency of pixels reaches 50%) defines the relative bone mineral content (BMC).

### Repeatability

To determine the repeatability of each method, 7 samples were selected that were evenly distributed across the reference range. Samples were blinded and analyzed 5 times in a random order, and in a different random order for each method. Mean, standard error of the mean, and coefficients of variation were calculated for each sample across 5 measurements (Supplementary Figure 1 and Supplementary Table 2).

### Surgical provocation of osteoarthritis

Mice underwent destabilization of the medial meniscus (DMM) ^13, 24^ osteoarthritis provocation surgery at 10 weeks-of-age. To minimize recovery time, mice were anaesthetized using inhaled isofluorane/oxygen mix (1-1.5L/min), and breathing rate was monitored visually throughout surgery. The medial menisco-tibial ligament (MMTL) of the right knee was transected using a 5mm microsurgical blade (World Precision Instruments, Hitchin, UK, Cat# 500249). Sham surgery was performed on the left knee, in which skin and joint capsule were opened but the MMTL was left intact. Each joint capsule was closed with a single suture, and the skin was closed using intradermal sutures. Temperature was maintained using a heat-pad and monitored by rectal thermometer. Post-operative buprenorphine (0.1mg/kg in saline, Vetergesic; Ceva United Kingdom, Amersham, UK), and carprofen (5mg/kg in saline, Rimadyl; Zoetis, Leatherhead, UK) were administered subcutaneously post-surgery while under anesthetic, and daily for 48h. Dosage was calculated based on individual weight. Physical pain indicators (including mobility, piloerection and hunching) were monitored post-surgery and daily for 48h. Surgeries were performed between 10am and 4pm in a designated surgical suite. Mice were returned to the home cage for recovery. Mice were housed 4 per cage in sibling groups and sacrificed 12 weeks post-surgery.

Left and right hindlimbs from half of the experimental cohort (16 mice; 2 mice/cage) were phenotyped by rapid-throughput joint phenotyping methods described above. The limbs of the other half of the surgical cohort (2 mice/cage) were sectioned and scored for osteoarthritis using previously-accepted gold-standard protocols. Numbers of mice in each group were based on group numbers in published guidelines^24^.

### Histological scoring of DMM surgical samples

Articulated left (sham) and right (DMM) hindlimbs were decalcified for 7-10 days in 10% EDTA and embedded in paraffin wax in a 90° flexed physiological position. Coronal 4µm sections were cut at 80µm intervals through the knee joint and stained with Safranin O/Fast Green and Weigert’s haematoxylin counterstain^77, 91^. Sections from 5 levels within the articulating zone centered around the midpoint were scored for each limb. All sections were scored by 3 blinded observers using the Osteoarthritis Research Society International (OARSI) scoring system^23^. Summed and maximum scores were calculated for each compartment (medial tibial plateau; MTP, medial femoral condyle; MFC, lateral tibial plateau; LTP, lateral femoral condyle; LFC) in a single limb from 5 sections. In cases when 5 sections were not available, scores were weighted according to the following equation:

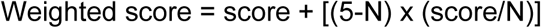

where N was the number of scoreable sections. Weighted scores were rounded to the nearest 0.5. Sum scores represent the sum of scores from 5 levels. The maximum score is the highest score (out of a maximum of 6). The total joint sum score is the sum of scores for all compartments across 5 levels. The total joint maximum score represents the maximum score from all compartments across 5 levels. Presence of osteophytes and displacement of the medial meniscus (indicating successful surgery) were also scored and noted separately. Lack of displacement of the medial meniscus was grounds for exclusion (n=1).

### Co-registration with previously accepted gold-standard histology

To co-register data obtained from rapid-throughput joint phenotype analysis directly with OARSI histological scoring, we performed further histology and OARSI scoring of samples from 3 mice that had previously been phenotyped in the rapid-throughput pipeline and found to have mild, intermediate and severe degrees of joint damage (Supplementary Figure 3). 4µm coronal sections were cut at 50µm intervals through the disarticulated tibiae and femora. Five sections centered around the midpoint of each femur and tibia were identified for OARSI scoring. Each limb was scored by 3 independent scorers, and the mean summed and maximum scores determined.

### Functional genomics analysis of human cartilage

We isolated chondrocytes from each cartilage sample as described^92^. Assays and analyses were performed as described^51^ and RNA sequencing was performed on the Illumina HiSeq2000 or Hiseq4000 (75bp paired-ends), with quality control including FastQC 0.11.5 (http://www.bioinformatics.babraham.ac.uk/projects/fastqc). For raw RNA sequencing data details see Data Availability Statement. For protein extracts, we carried out digestion, 6-plex or 10-plex tandem mass tag labelling and peptide fractionation. For samples from 12 knee osteoarthritis patients, we applied a liquid chromatography mass spectrometry (LC-MS) analysis using the Dionex Ultimate 3000 ultra-high-performance liquid chromatography (UHPLC) system coupled with the high-resolution LTQ Orbitrap Velos mass spectrometer (Thermo Fisher Scientific GmbH, Dreieich, Germany). For all remaining samples, LC-MS analysis was performed on the Dionex Ultimate 3000 UHPLC system coupled with the Orbitrap Fusion Tribrid Mass Spectrometer (Thermo Fisher Scientific). For proteomics data details see Data Availability Statement.

### Skeletal phenotyping

Rapid-throughput skeletal phenotyping was performed as previously described^16, 17, 19^, on female 16-week old mice. A minimum of three animals were phenotyped per line.

### Prioritization pipeline

A prioritization pipeline was developed based on the severity of the mutant mouse phenotype and i) additional skeletal abnormalities identified in mutant mice, ii) gene expression in skeletal cells and tissues, iii) gene association with human monogenic and complex diseases, and iv) structured literature searching (Supplementary Table 6).

Genes were allocated overall priority scores out of a total maximum of 21 as follows:

A maximum score of 6 for rapid-throughput joint phenotyping (outlier according to criteria of reference range, Wilcoxon test followed by Bonferroni adjustment and Mahalanobis analysis, 1 point each; abnormal articular cartilage morphology, articular cartilage damage, subchondral bone morphology, 1 point each).

A maximum score of 5 for abnormal skeletal phenotypes reported in genetically modified mice by other studies. The following resources were searched: 1) OBCD bone phenotyping pipeline for abnormal skeletal phenotypes of structure and strength (www.boneandcartilage.com), 2) International Mouse Phenotyping Resource of Standardized Screens (IMPReSS; https://www.mousephenotype.org/impress), 3) Deciphering the Mechanisms of Developmental Disorders database (DMDD; https://dmdd.org.uk/), 4) PubMed (https://www.ncbi.nlm.nih.gov/pubmed/) using the search criteria “GENE NAME AND (knockout OR deletion OR mutation)”, and 5) Mouse Genome Informatics database (http://www.informatics.jax.org/). The presence of an abnormal skeletal phenotype was assigned 1 point per database.

A maximum score of 4 for expression in skeletal tissues and cells (the skeleton, chondrocytes, osteoblasts/osteocytes and osteoclasts; Supplementary Table 6). Expression in the skeleton (1 point) was determined by searching MGI (http://www.informatics.jax.org/) and BioGPS (http://biogps.org/#goto=welcome). An expression level greater than the median was considered to indicate tissue or cell type expression. Expression in chondrocytes (1 point) was determined by searching published transcriptome data^27^ and SkeletalVis^28^ (http://phenome.manchester.ac.uk/). Search was limited by species (human) in both datasets. In Kean et al.^27^ a baseMean expression greater than the median was considered to indicate cell type expression. In SkeletalVis searches, a fold-change greater than 2 (P<0.001) was considered to indicate cell type expression. Expression in osteoblasts and/or osteocytes (1 point) was determined by searching BioGPS, SkeletalVis, and osteocyte RNAseq data^19^. Expression in osteoclasts (1 point) was determined by searching BioGPS, human osteoclast RNAseq datasets, and SkeletalVis. A baseMean expression greater than the median was considered to indicate cell type expression.

A maximum score of 4 for association with monogenic and polygenic skeletal disease (Supplementary Table 6). Monogenic disease association was determined by searching MGI for human-mouse disease association, and Online Mendelian Inheritance in Man (OMIM, https://www.omim.org/) by gene name (1 point). Polygenic disease association was assessed by searching the European Bioinformatics Institute Genome-wide Association Studies database (EBI GWAS Catalogue, https://www.ebi.ac.uk/gwas/) for association with arthritis and skeletal diseases (1 point each).

A maximum score of 2 for publications related to skeletal tissues and cells (Supplementary Table 6). Systematic structured searches of PubMed and Google Scholar (https://scholar.google.co.uk/) were performed using the search string “GENE NAME AND (arthritis OR cartilage OR skeleton OR chondrocyte OR osteoblast OR osteocyte OR osteoclast)”. For PubMed results, the following scores were assigned: 0 (<1 publication), 0.5 (1–24 publications), or 1 (>25 publications). For Google Scholar results, the following scores were assigned: 0 (<100 results), 0.5 (100-999 results), or 1 (>1000 results).

### Quantitation and Statistical Analysis

Statistical details and results of experiments are included in the relevant results section, figure legends, Reporting Summary and below. For all experiments, n refers to biological replicates (number of mice or histological sections). Measurements were taken from distinct samples unless otherwise indicated. Exact *P*-values are reported in Supplementary Tables 4 and 8.

#### Rapid-throughput joint phenotyping

Reference range data was calculated from 100 wild-type mice for each parameter, and the mean, standard deviation, median and percentiles determined (Microsoft Excel 2016 and GraphPad Prism 8, Supplementary Table 1). Frequency distribution of datasets was assessed by the Shapiro-Wilk normality test. Reference ranges 2 standard deviation above or below the mean (normally-distributed parameters), or between the 2.5^th^-97.5^th^ percentiles (non-normally distributed parameters), were determined. An outlier was defined if the mean value of a phenotype parameter was outside the defined reference range.

Normally distributed parameters were articular cartilage volume (LTP), maximum articular cartilage thickness (LTP), subchondral bone volume/tissue volume (LTP), subchondral trabecular thickness (LTP, MTP), subchondral trabecular number (LTP), subchondral bone mineral density (LTP), and subchondral bone mineral content (LTP, MTP). Non-normally distributed parameters were articular cartilage volume (MTP), median articular cartilage thickness (LTP, MTP), maximum articular cartilage thickness (MTP), articular cartilage surface damage (LTP, MTP), subchondral bone volume/tissue volume (MTP), subchondral trabecular number (MTP) and subchondral bone mineral density (MTP).

To determine the statistical significance of outlier parameters identified relative to the reference range, we performed non-parametric 2-tailed Wilcoxon rank sum tests (Supplementary Table 4, *wilcox.test* function in R). For each parameter, this analysis calculated whether the distribution parameters in mutant samples was significantly different to the distribution of parameters in the 100-sample WT reference range. To account for multiple testing when defining statistical significance, we calculated the effective number of tests and then applied a Bonferroni correction as follows. Because several of the 18 phenotype parameters determined in the 100 WT mice used to calculate reference data were correlated (Supplementary Table 5, \**P*<0.05), we first calculated the number of effective tests using WT data in R. We calculated the pairwise correlations between all 18 parameters and obtained eigenvalues of the correlation matrix using the *eigen* function in R. We then calculated the number of effective tests (N_eff_) as:

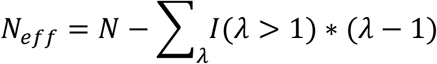

where N=18 is the number of parameters, and *λ* denotes the eigenvalues, resulting in an effective N_eff_≤=8.8. Thus, applying a Bonferroni correction for the effective number of tests, statistical significance for the differences between WT reference data and results from each mutant mouse line was P<0.00568, with a second more stringent threshold of P<0.0001 also correcting for the number of mouse lines.. For each of 18 parameters across the 50 mouse lines, differences between mutant and WT values were determined by 2-tailed Wilcoxon rank sum tests.

Differences between young and old wild-type mice, and between *Dio2^Thr92^* and *Dio2^Ala92^* mice, were determined using unpaired 2-tailed *t*-tests (Microsoft Excel 2016, normally-distributed parameters) or 2-tailed Wilcoxon rank sum tests (GraphPad Prism 8, non-normally distributed parameters).

Differences between sham and DMM-operated limbs from the same mouse were determined using paired 2-tailed *t*-tests (Microsoft Excel 2016, normally distributed parameters), or 2-tailed Wilcoxon matched pairs signed rank tests (GraphPad Prism 8, non-normally distributed parameters).

Differences between OARSI histology scores in sections from sham and DMM-operated knee joints were determined using paired 2-tailed Wilcoxon matched-pairs signed rank tests (GraphPad Prism 8). Differences between OARSI histology scores of sections from young and aged wild-type mice were determined using unpaired 2-tailed Wilcoxon rank sum tests (GraphPad Prism 8).

#### Power calculations

To determine the sample size required, we determined coefficients of variation (CV, standard deviation/mean) for normally distributed parameters, and percentage median absolute deviation from the median (median absolute deviation from the median (MAD)/median) for non-normally distributed parameters (Supplementary Table 1). The effect size (d) was set at twice the standard deviation (σ) or the MAD. To detect this effect size with 80% power, the number of mice per group (N) was determined by

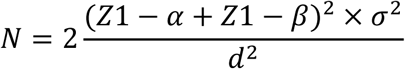

where power index (Z_1-α_+Z_1-β_)^2^ is 6.2 for a power of 80%. Thus, 4 mice are required to detect an effect size of 2 standard deviations at 80% power, and 6 mice at 90% power, for all parameters. For non-parametric data, 1-4 mice are required, depending on the parameter, to detect an outlier phenotype lying outside the 95% confidence intervals of the reference range (Supplementary Table 1).

#### Mahalanobis analysis

Robust Mahalanobis distances^93, 94^ were determined to identify outliers in multivariate data. These distances measure how far each observation is from the center of a data cluster, taking into account the variances of the variables and the covariances of pairs of variables. Robust Mahalanobis distances (MD_i_)

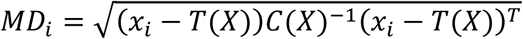

were calculated for each sample (represented by a vector of multivariate observations *x_i_*). T(X) is a robust (i.e. relative unaffected by outliers) estimate of the mean vector and C(X) is a robust estimate of the covariance matrix. Under the assumption of multivariate normality, the distribution of 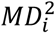 is approximately chi-squared with p degrees of freedom (where p is the number of variables). This means that any observation *x_i_* with 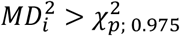 can be considered an outlier. The assumption of multivariate normality was checked by plotting the ordered distances against the corresponding quantiles of an appropriate chi-squared distribution. Robust estimates of the mean and covariance matrix are used so that potential outliers are not masked. The masking effect, by which outliers do not necessarily have a large Mahalanobis distance, can be caused by a cluster of outliers that attract the mean and inflate the covariance in its direct. By using a robust estimate of the sample mean and covariance, the influence of these outliers is removed and the Mahalanobis distance is able to expose all outliers. The minimum volume ellipsoid method was used to calculate robust estimates of the mean and covariance matrix. Given *n* observations and *p* variables, the minimum volume ellipsoid method seeks an ellipsoid containing

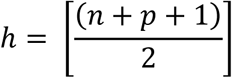

*h* points of minimum volume. All multivariate analysis was conducted in R (R Project for Statistical Computing). Overall, a mouse line was defined to have an abnormal joint phenotype using this method only when 50% or more individual samples from that line were identified as outliers following Mahalanobis analysis.

### Analysis of RNA and protein in human tissue

Full details of this analysis are described^51^ in (Steinberg *et al*., accompanying related manuscript). Briefly, gene-level RNA quantification was carried out using Salmon 0.8.2^95^ and tximport ^96^. After quality control, we retained expression estimates for 15,249 genes with counts per million of 1 or higher in at least 40 cartilage samples (combining low-grade and high-grade), and matched low-grade and high-grade cartilage samples from 83 patients. To identify gene expression differences between high-grade and low-grade cartilage, we carried out multiple analyses using limma^97, 98^, DESeq2^99^ and edgeR^100^ with 5 different designs to account for potential confounders. This yielded 2557 genes with significant differential expression between low-grade and high-grade cartilage at 5% false discovery rate (FDR) across all 5 analysis designs and all 5 testing methods.

To carry out protein identification and quantification, we submitted the mass spectra to SequestHT search in Proteome Discoverer 2.1, and searched all spectra against a UniProt fasta file that contained 20,165 reviewed human entries. After quality control, we retained 4801 proteins that were quantified in at least 30% of samples, and matched low-grade and high-grade cartilage samples from 99 patients. To account for protein loading, abundance values were normalized by the sum of all protein abundances in a given sample, then log2-transformed and quantile normalised. An analysis for differential abundance was carried out using limma^97^ with 2016 proteins showing significant differences at 5% FDR.

We used Ensembl38p10 to identify human orthologues for 50 mouse genes phenotyped by the OBCD joint phenotyping pipeline. We identified high-confidence one-to-one orthologues for 43/50 genes.

### Data Availability Statement

All datasets generated and/or analyzed during the current study are included in Supplemental Tables, and available from the corresponding authors on reasonable request or as detailed below. All figures have raw data associated with them. Raw data to support the findings for Figures 1-7 and Supplementary Figures 1-7 are available from the corresponding authors on reasonable request. Additionally, raw data to support the findings for Figure 5 are available as specified below. Raw RNA sequencing data is available in Steinberg et al.,^51, 101^ and has been deposited in the European Genome-Phenome Archive [EGA; https://www.ebi.ac.uk/ega/home) with the identifiers EGAS00001002255, EGAD00001003355 (n=17), EGAD00001003354 (n=9), EGAD00001001331 (n=12). Proteomics data is available in Steinberg et al.,^51, 101^and has been deposited in the PRoteomics IDEntifications database (PRIDE; https://www.ebi.ac.uk/pride/archive/) with the identifiers PXD00202014 and PDX014666, username, password: AoeRBm3e).

### Code availability

Code generated during this project is available as Supplementary Data 1.

