## Supplementary figures tables and data for "Accelerating functional gene discovery in osteoarthritis"

**Supplementary Information**

***Supplementary Figures***

***Supplementary Figure 1***

***Reference range data for joint phenotype parameters***

***a.*** Graphs showing articular cartilage (articular cartilage volume (ACV), median articular cartilage thickness (Median AC Th), maximum articular cartilage thickness (Max AC Th), AC damage area) and subchondral bone (subchondral bone volume per tissue volume (SC BV/TV), SC trabecular thickness (SC Tb.Th), SC trabecular number (SC Tb.N), SC bone mineral density (SC BMD), SC bone mineral content (SC BMC)) parameters in the lateral and medial tibial plateaux (LTP, MTP) of 16-week-old male WT mice (n=100). Values from individual mice are shown as black dots with reference ranges defined as mean +/-2.0 standard deviations (normally distributed parameters), or median and 2.5$^{th}$-97.5th percentiles (non-normally distributed parameters) and shown in grey boxes with the median or mean values shown as horizontal lines.

***b.*** Repeatability of measurement for each parameter determined by five repeat analyses of seven samples obtained from individual mice (x-axis). Black symbols are repeat measures for each sample. Reference ranges are shown in grey boxes, the median/mean as horizontal lines and mean ± standard error of the mean is reported for each sample.

***Supplementary Figure 2***

***Joint abnormalities following osteoarthritis provocation surgery and severe early onset osteoarthritis in Pitx1$^{+/-}$ mice***

***a.*** Graphs showing articular cartilage (articular cartilage volume (ACV), median articular cartilage thickness (Median AC Th), maximum articular cartilage thickness (Max AC Th), AC damage area) and subchondral bone (subchondral bone volume per tissue volume (SC BV/TV), SC trabecular thickness (SC Tb.Th), SC trabecular number (SC Tb.N), SC bone mineral density (SC BMD), SC bone mineral content (SC BMC)) parameters in the lateral and medial tibial plateaux (LTP, MTP) of sham and DMM operated knees of WT mice (n=16). Paired data (sham versus DMM) are shown for each mouse.

***b.*** Graphs showing Osteoarthritis Research Society International (OARSI) histological scores in sham and DMM operated knees of WT mice (n=11). Parameters include the summed and maximum scores on the LTP, lateral femoral condyle (LFC), MTP, medial femoral condyle (MFC), and combined values for the total joint.

***c.*** Graphs showing articular cartilage and subchondral bone parameters in the LTP and MTP of *Pitx1*$^{+/-}$ mice (n=7). **P*<0.05, ***P*<0.01, ****P*<0.001 (DMM), **P*<0.00568, ***P*<0.001, ****P*<0.0001 (*Pitx1*$^{+/-}$).

***Supplementary Figure 3***

***Validation of novel imaging methods***

***a.*** Iodine contrast enhanced μCT images of medial tibial plateau articular cartilage (red volumes) and subchondral bone (blue volumes) from 22-week-old WT male mice 12 weeks after sham operation or following destabilization of medial meniscus surgery (DMM) that resulted in either mild, intermediate or severe osteoarthritis.

***b.*** Back-scattered electron-scanning electron microscopy images of medial tibial plateau joint surface replicas from sham and DMM operated WT mice with mild, intermediate or severe osteoarthritis.

***c.*** Pseudocoloured X-ray microradiography images of proximal tibia and the medial tibial plateau subchondral bone region of interest (dashed box) from sham and DMM operated WT mice with mild, intermediate or severe osteoarthritis. Low bone mineral content (BMC) is yellow and high BMC is pink.

***d.*** Coronal sections of medial tibial plateaux stained with Safranin-O/Fast green from three sham and DMM operated WT mice with mild, intermediate or severe osteoarthritis. Arrows indicate areas of cartilage damage. Scale bars = 100μm (A, B, D) and 1mm (C).

***e.*** Graphs showing articular cartilage (articular cartilage volume (ACV), median articular cartilage thickness (Median AC Th), maximum articular cartilage thickness (Max AC Th) and AC damage area) and subchondral bone (subchondral bone volume per tissue volume (SC BV/TV), SC trabecular thickness (SC Tb.Th), SC trabecular number (SC Tb.N), SC bone mineral density (SC BMD), SC bone mineral content (SC BMC)) parameters together with summed and maximum OARSI histological scores on the medial tibial plateaux of sham and DMM operated knees from three mice with mild, intermediate and severe OA. Results from entire cohort in grey, with mild, intermediate and severe examples coloured according to key.

***Supplementary Figure 4***

***Early onset osteoarthritis in Bhlhe40$^{-/-}$ and Sh3pb4$^{-/-}$ mutant mice***

***a.*** Graphs showing articular cartilage (articular cartilage volume (ACV), median articular cartilage thickness (Median AC Th), maximum articular cartilage thickness (Max AC Th) and AC damage area) and subchondral bone (subchondral bone volume per tissue volume (SC BV/TV), SC trabecular thickness (SC Tb.Th), SC trabecular number (SC Tb.N), SC bone mineral density (SC BMD), SC bone mineral content (SC BMC)) parameters in the lateral and medial tibial plateaux (LTP, MTP) from *Bhlhe40*$^{-/-}$ mice (n=3).

***b.*** Graphs showing articular cartilage and subchondral bone parameters in the LTP and MTP from *Sh3pb4*$^{-/-}$ mice (n=5). Data from individual mice are shown, mean values represented by horizontal bars, reference ranges as grey boxes with the median or mean values shown as dotted lines. *$P$<0.00568, **$P$<0.001.

***Supplementary Figure 5***

***Application 1: Early onset osteoarthritis in mice with deletion of genes differentially expressed in human osteoarthritis cartilage***

***a.*** Graphs showing articular cartilage (articular cartilage volume (ACV), median articular cartilage thickness (Median AC Th), maximum AC Th (Max AC Th), AC damage area) and subchondral bone (subchondral bone volume per tissue volume (SC BV/TV), SC trabecular thickness (SC Tb.Th), SC trabecular number (SC Tb.N), SC bone mineral density (SC BMD), SC bone mineral content (SC BMC)) parameters in the lateral and medial tibial plateaux (LTP, MTP) of *Unk* $^{-/-}$ mice (n=4).

***b.*** Graphs showing articular cartilage and subchondral bone parameters in the LTP and MTP of *Josd1*$^{-/-}$ mice (n=6).

***c.*** Graphs showing articular cartilage and subchondral bone parameters in the LTP and MTP of *Gsdme*$^{-/-}$ mice (n=4). Data from individual mice are shown, mean values represented by horizontal bars, reference ranges as grey boxes with the median or mean values shown as dotted lines. *$P$<0.00568.

***Supplementary Figure 6***

***Application 2: Age-related joint degeneration***

Graphs showing articular cartilage (articular cartilage volume (ACV), median articular cartilage thickness (Median AC Th), maximum AC Th (Max AC Th), AC damage area) and subchondral bone (subchondral bone volume per tissue volume (SC BV/TV), SC trabecular thickness (SC Tb.Th), SC trabecular number (SC Tb.N), SC bone mineral density (SC BMD), SC bone mineral content (SC BMC)) parameters in the lateral and medial tibial plateaux (LTP, MTP) of 4-month-old (n=6) and 12-month-old (n=6) WT mice. *$P$<0.05, **P<0.01.

***Supplementary Figure 7***

***Application 3: Mice with a Dio2$^{Ala92}$ polymorphism are protected from osteoarthritis***

Graphs showing articular cartilage (articular cartilage volume (ACV), median articular cartilage thickness (Median AC Th), maximum articular cartilage thickness (Max AC Th), AC damage area) and subchondral bone (subchondral bone volume per tissue volume (SC BV/TV), SC trabecular thickness (SC Tb.Th), SC trabecular number (SC Tb.N), SC bone mineral density (SC BMD), SC bone mineral content (SC BMC)) parameters in the lateral and medial tibial plateaux (LTP, MTP) from *Dio2*$^{Thr92}$ (n=9) and *Dio2*$^{Ala92}$ (n=10) mice. *$P$<0.05, **$P$<0.01.

***Supplementary Tables***

***Supplementary Table 1***

Datasets for all joint phenotyping pipeline parameters in 100 wild-type mice.

***Supplementary Table 2***

Repeatability of measurement of joint phenotyping parameters.

***Supplementary Table 3***

Joint phenotyping pipeline data from 50 unselected mouse lines.

***Supplementary Table 4***

Statistical analysis of joint phenotyping pipeline data from 50 unselected mouse lines.

***Supplementary Table 5***

Spearman correlation matrix for statistically significant correlation coefficients (R-values, $P<0.05$) between joint phenotyping parameters.

***Supplementary Table 6***

Prioritization analysis of 25 mouse lines with abnormal joint phenotypes.

***Supplementary Table 7***

Animal species, strain, source, sex and unique identifier or Research Resource Identifier (RRID).

***Supplementary Table 8***

Actual *P*-values for validation of new methods with surgical provocation of osteoarthritis by DMM surgery, Application 2 and Application 3.

***Supplementary Data***

***Supplementary Data 1***

ImageJ macros for automated joint surface replication quantitation and subchondral X-ray microradiography.

Supplementary Figure 1

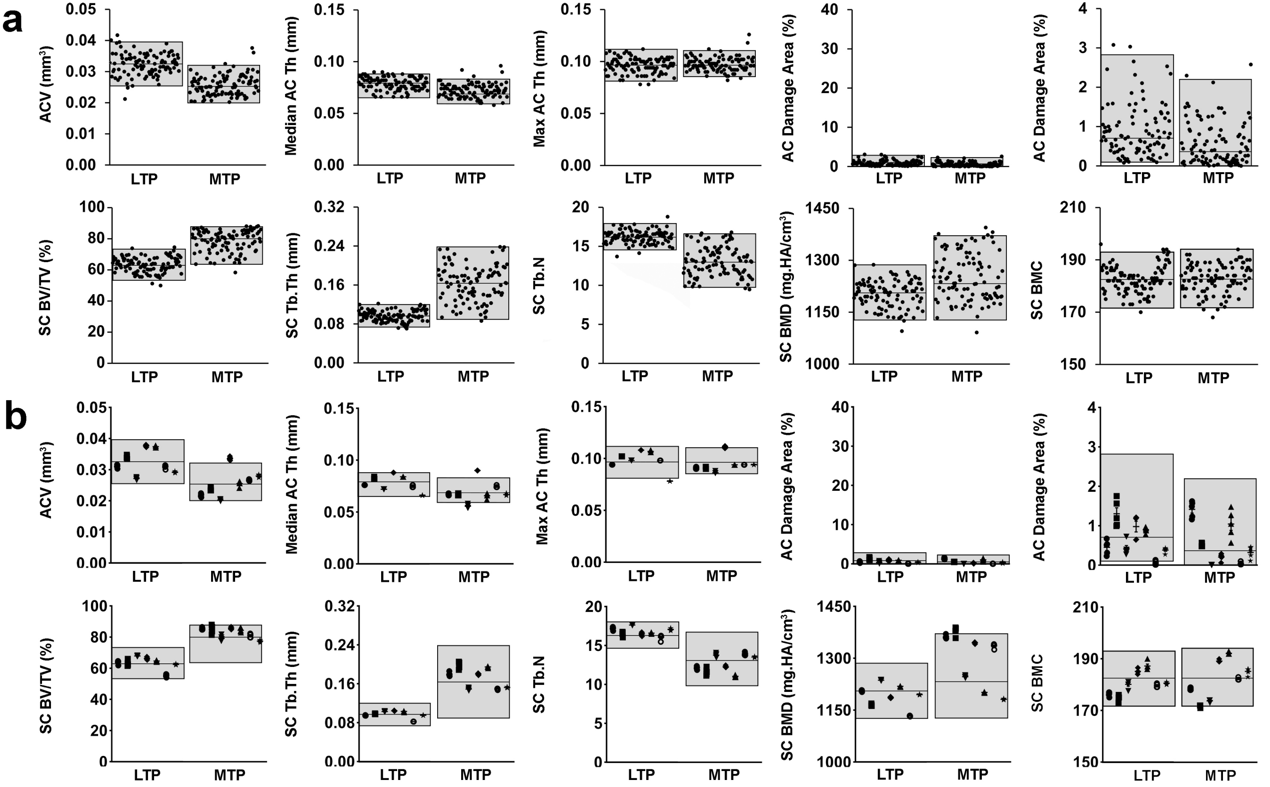

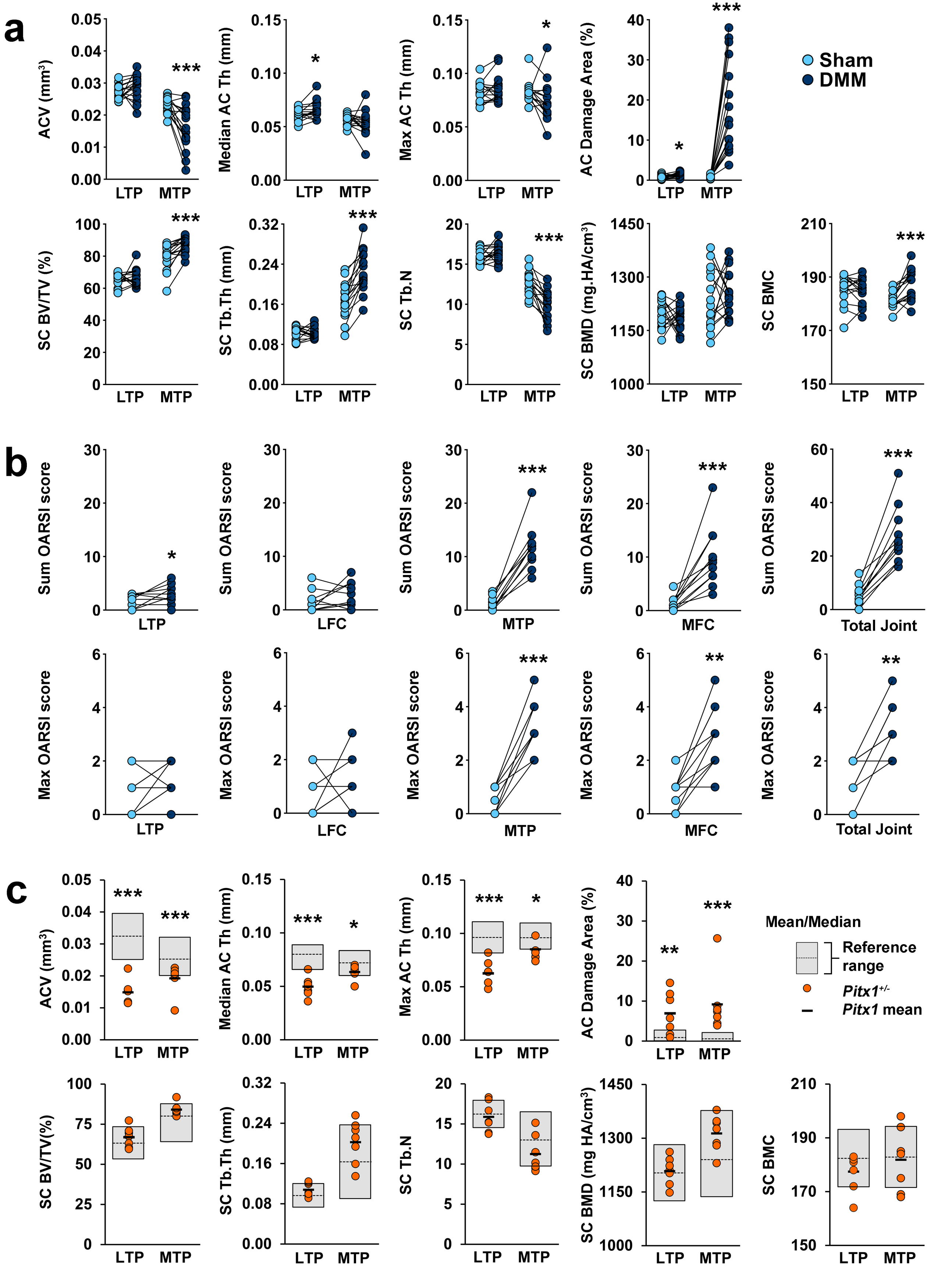
a
ACV (mm³)
Median AC Th (mm)
Max AC Th (mm)
AC Damage Area (%)
Sham
DMM
SC BV/TV (%)
SC Tb.Th (mm)
SC Tb.N
SC BMD (mg.HA/cm³)
SC BMC
LTP
MTP
b
Sum OARSI score
Max OARSI score
LTP
LFC
MTP
MFC
Total Joint
c
ACV (mm³)
Median AC Th (mm)
Max AC Th (mm)
AC Damage Area (%)
Mean/Median
Reference range
Pitx1+/-
Pitx1 mean
SC BV/TV(%)
SC Tb.Th (mm)
SC Tb.N
SC BMD (mg HA/cm³)
SC BMC

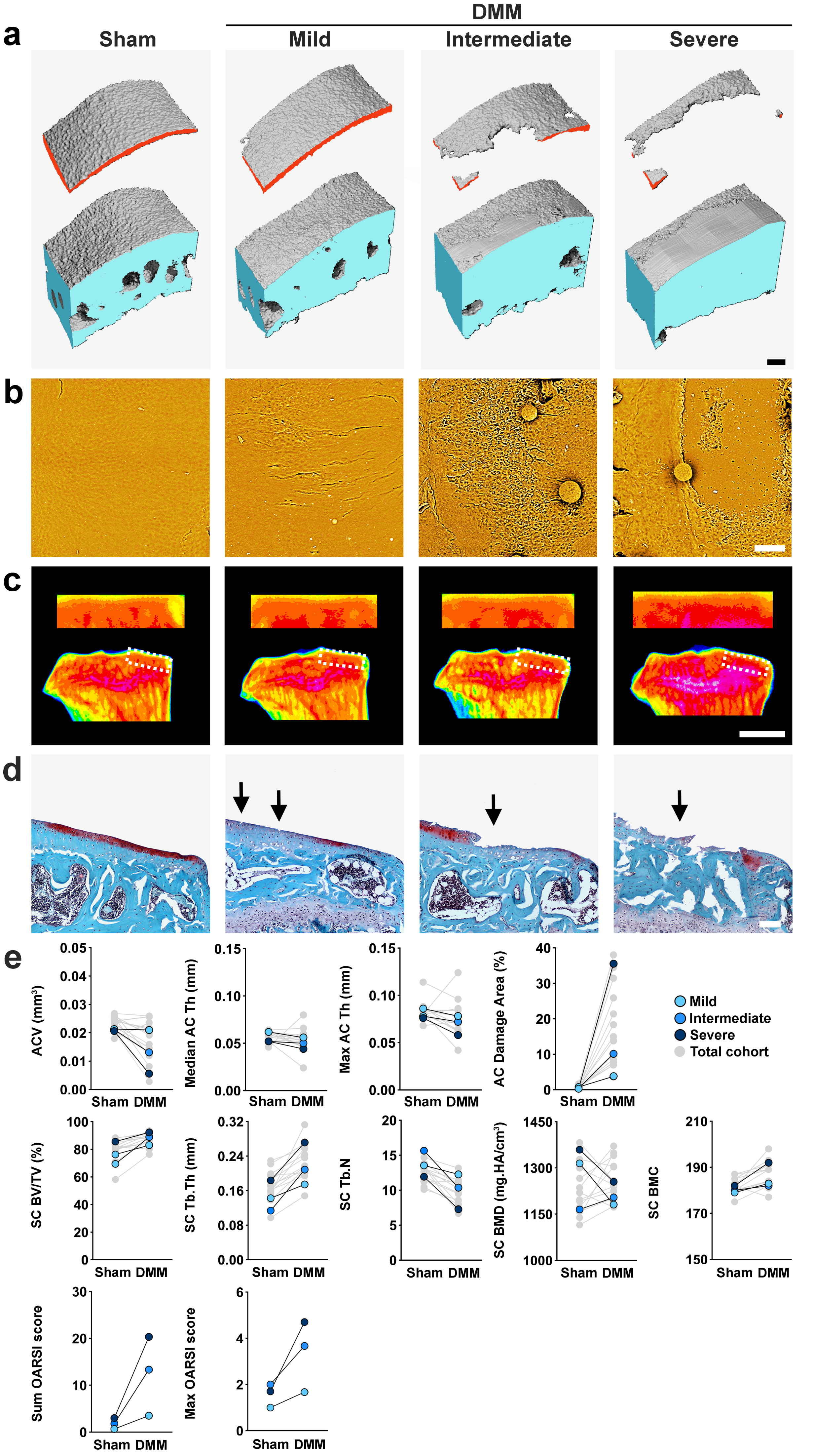
DMM
a
Sham
Mild
Intermediate
Severe
b
c
d
e
ACV (mm³)
Median AC Th (mm)
Max AC Th (mm)
AC Damage Area (%)
Mild
Intermediate
Severe
Total cohort
SC BV/TV (%)
SC Tb.Th (mm)
SC Tb.N
SC BMD (mg.HA/cm³)
SC BMC
Sum OARSI score
Max OARSI score
Sham DMM

Supplementary Figure 4

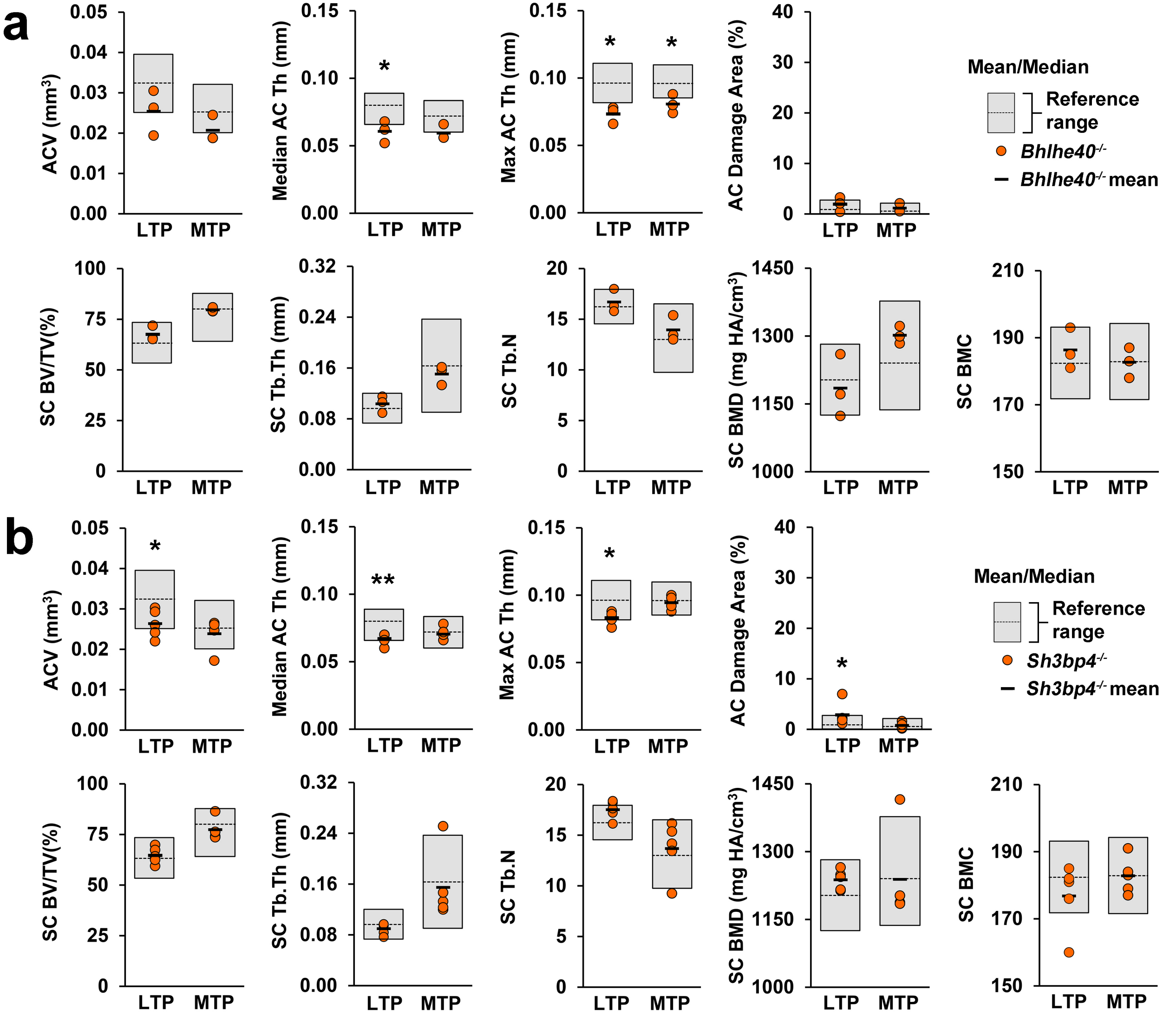

Supplementary Figure 5

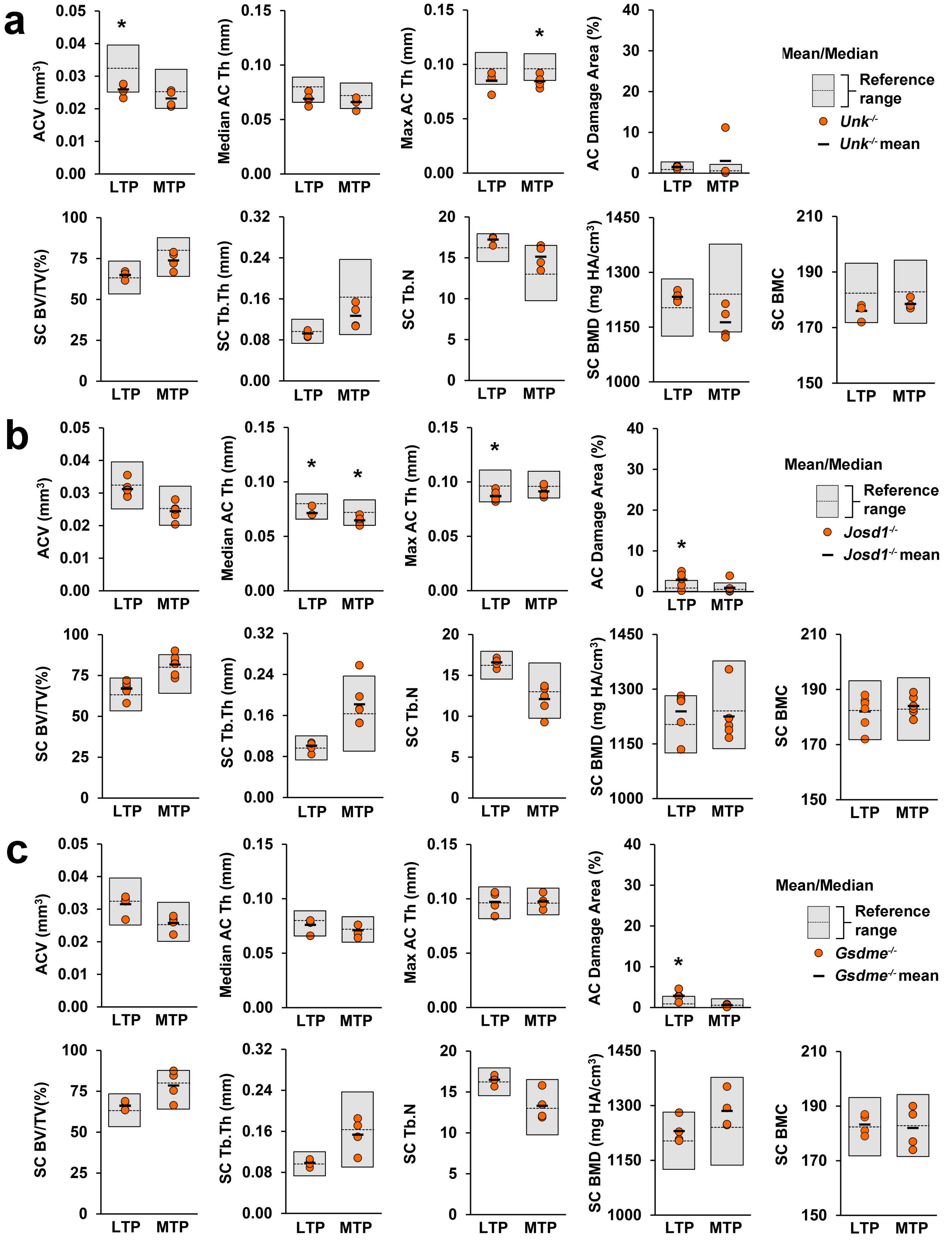

### Supplementary Figure 6

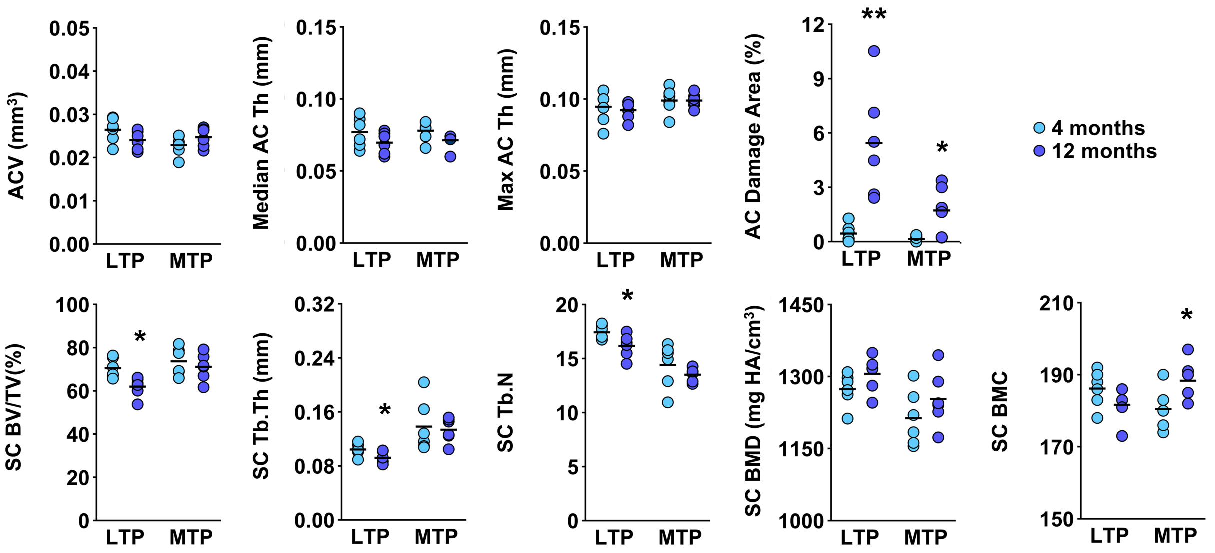

**Supplementary Figure 7**

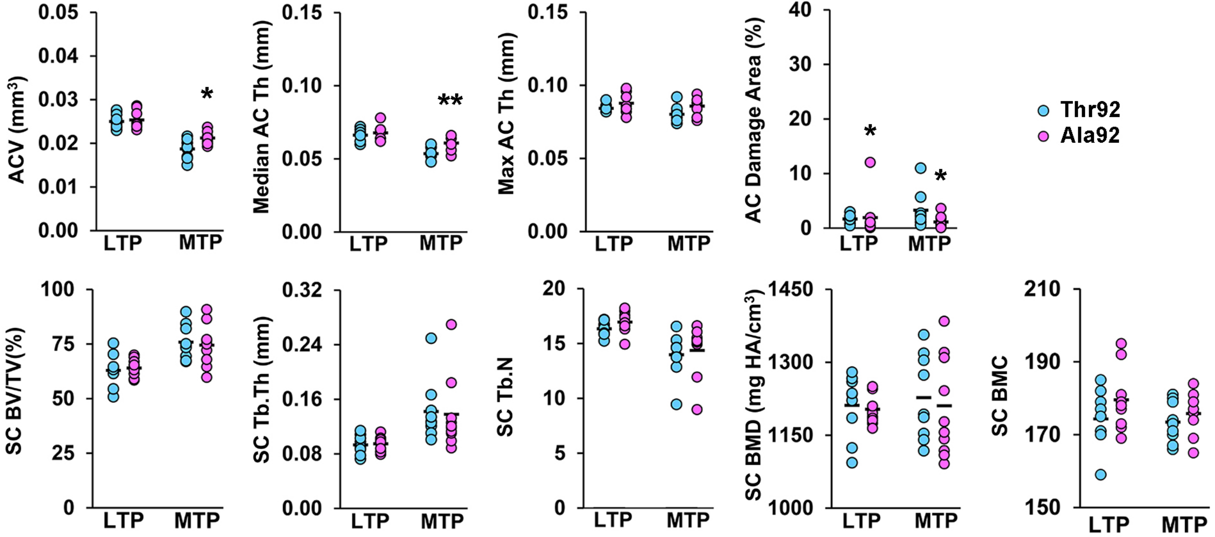

Datasets for all joint phenotyping pipeline parameters in 100 wild-type mice.

| Tissue compartment: | Articular cartilage | | | | | | | | Subchondral bone | | | | | | | | | |
|---|---|---|---|---|---|---|---|---|---|---|---|---|---|---|---|---|---|---|
| Parameter: | ACV | | Median AC Th | | Maximum AC Th | | AC Damage Area | | SC BV/TV | | SC Tb.Th | | SC Tb.N | | SC BMD | | SC BMC | |
| Unit: | (mm$^3$) | | (mm) | | (mm) | | (%) | | (%) | | (mm) | | | | (mg.HA/cm$^3$) | | (median grey level) | |
| Technique: | ICE-μCT | | ICE-μCT | | ICE-μCT | | JSR | | ICE-μCT | | ICE-μCT | | ICE-μCT | | ICE-μCT | | scXRM | |
| Reference range: | mean ± 2SD | 2.5–97.5% | 2.5–97.5% | 2.5–97.5% | mean ± 2SD | 2.5–97.5% | 2.5–97.5% | 2.5–97.5% | mean ± 2SD | 2.5–97.5% | mean ± 2SD | mean ± 2SD | mean ± 2SD | 2.5-97.5% | mean ± 2SD | 2.5–97.5% | mean ± 2SD | mean ± 2SD |
| ID of 100 WT mice | LTP | MTP | LTP | MTP | LTP | MTP | LTP | MTP | LTP | MTP | LTP | MTP | LTP | MTP | LTP | MTP | LTP | MTP |
| M02287278 | 0.033 | 0.023 | 0.080 | 0.066 | 0.092 | 0.096 | 0.64 | 0.58 | 64.47 | 82.81 | 0.107 | 0.191 | 15.3 | 11.6 | 1191 | 1252 | 196 | 184 |
| M02287310 | 0.040 | 0.031 | 0.086 | 0.082 | 0.108 | 0.106 | 1.47 | 0.08 | 68.37 | 80.79 | 0.120 | 0.180 | 15.4 | 12.2 | 1286 | 1270 | 187 | 179 |
| M02287113 | 0.028 | 0.021 | 0.078 | 0.068 | 0.086 | 0.090 | 1.05 | 0.44 | 66.31 | 86.49 | 0.100 | 0.233 | 16.7 | 9.9 | 1201 | 1363 | 186 | 180 |
| M02287250 | 0.036 | 0.027 | 0.086 | 0.082 | 0.098 | 0.104 | 0.74 | 0.11 | 58.26 | 70.59 | 0.086 | 0.134 | 15.6 | 13.4 | 1201 | 1187 | 179 | 179 |
| M02287252 | 0.030 | 0.026 | 0.078 | 0.074 | 0.086 | 0.096 | 0.86 | 0.02 | 67.19 | 80.42 | 0.112 | 0.171 | 15.3 | 13.0 | 1231 | 1294 | 188 | 182 |
| M02295226 | 0.031 | 0.027 | 0.084 | 0.078 | 0.096 | 0.098 | 0.98 | 1.49 | 61.04 | 63.71 | 0.087 | 0.094 | 16.4 | 16.4 | 1209 | 1134 | 177 | 176 |
| M02295227 | 0.033 | 0.028 | 0.088 | 0.078 | 0.110 | 0.100 | 0.65 | 1.28 | 64.07 | 76.11 | 0.100 | 0.143 | 16.4 | 14.0 | 1220 | 1178 | 178 | 185 |
| M02289441 | 0.028 | 0.020 | 0.076 | 0.068 | 0.094 | 0.092 | 0.88 | 2.30 | 65.19 | 86.79 | 0.094 | 0.187 | 16.8 | 11.9 | 1211 | 1376 | 175 | 179 |
| M02286957 | 0.036 | 0.027 | 0.082 | 0.074 | 0.094 | 0.108 | 2.46 | 1.59 | 65.54 | 86.22 | 0.103 | 0.217 | 16.6 | 10.4 | 1234 | 1221 | 187 | 192 |
| M02302186 | 0.039 | 0.024 | 0.088 | 0.070 | 0.098 | 0.100 | 1.57 | 1.10 | 63.48 | 83.09 | 0.099 | 0.189 | 16.7 | 11.5 | 1190 | 1252 | 185 | 187 |
| M02302177 | 0.042 | 0.029 | 0.086 | 0.076 | 0.100 | 0.106 | 1.00 | 1.52 | 62.92 | 84.46 | 0.102 | 0.203 | 15.4 | 11.2 | 1230 | 1238 | 182 | 190 |
| M02295224 | 0.027 | 0.024 | 0.072 | 0.066 | 0.092 | 0.094 | 0.89 | 0.93 | 63.25 | 71.61 | 0.092 | 0.118 | 16.8 | 16.2 | 1184 | 1270 | 182 | 182 |
| M02295228 | 0.031 | 0.026 | 0.082 | 0.074 | 0.102 | 0.096 | 0.92 | 0.89 | 61.54 | 81.99 | 0.096 | 0.165 | 15.8 | 13.3 | 1174 | 1340 | 182 | 192 |
| M02302185 | 0.040 | 0.027 | 0.086 | 0.074 | 0.108 | 0.100 | 0.39 | 0.09 | 69.80 | 87.27 | 0.111 | 0.234 | 16.1 | 9.8 | 1243 | 1278 | 189 | 188 |
| M02303582 | 0.028 | 0.025 | 0.076 | 0.078 | 0.088 | 0.104 | 0.56 | 0.42 | 68.91 | 79.24 | 0.102 | 0.156 | 17.8 | 13.2 | 1233 | 1212 | 186 | 186 |
| M02302193 | 0.038 | 0.025 | 0.084 | 0.064 | 0.102 | 0.096 | 0.49 | 0.31 | 67.61 | 86.60 | 0.105 | 0.217 | 16.1 | 10.2 | 1228 | 1303 | 187 | 189 |
| M02303581 | 0.035 | 0.028 | 0.080 | 0.078 | 0.096 | 0.102 | 1.59 | 0.14 | 65.94 | 84.26 | 0.106 | 0.212 | 16.4 | 10.9 | 1195 | 1345 | 186 | 187 |
| M02303583 | 0.034 | 0.025 | 0.084 | 0.072 | 0.100 | 0.094 | 3.08 | 0.49 | 67.09 | 83.99 | 0.107 | 0.177 | 16.0 | 12.4 | 1181 | 1332 | 190 | 183 |
| M02299990 | 0.033 | 0.024 | 0.076 | 0.068 | 0.106 | 0.092 | 0.34 | 0.15 | 61.76 | 79.06 | 0.095 | 0.162 | 16.3 | 13.0 | 1150 | 1247 | 182 | 179 |
| M02299993 | 0.033 | 0.020 | 0.082 | 0.064 | 0.098 | 0.090 | 0.35 | 0.64 | 59.24 | 85.03 | 0.102 | 0.203 | 13.7 | 10.7 | 1195 | 1341 | 186 | 183 |
| M02303128 | 0.030 | 0.022 | 0.082 | 0.070 | 0.104 | 0.096 | 0.67 | 0.54 | 69.42 | 85.76 | 0.110 | 0.202 | 16.1 | 11.0 | 1238 | 1251 | 183 | 181 |
| M02299995 | 0.021 | 0.021 | 0.068 | 0.074 | 0.082 | 0.100 | 0.52 | 0.24 | 62.55 | 67.31 | 0.085 | 0.105 | 16.4 | 15.6 | 1226 | 1190 | 180 | 183 |
| M02303126 | 0.036 | 0.028 | 0.090 | 0.076 | 0.104 | 0.106 | 1.03 | 1.40 | 66.03 | 77.76 | 0.101 | 0.178 | 16.5 | 12.1 | 1248 | 1219 | 187 | 188 |
| M02303124 | 0.034 | 0.025 | 0.082 | 0.070 | 0.096 | 0.096 | 0.37 | 1.29 | 69.71 | 84.29 | 0.107 | 0.224 | 17.2 | 9.9 | 1227 | 1263 | 188 | 189 |
| M02311469 | 0.026 | 0.025 | 0.066 | 0.076 | 0.086 | 0.098 | 0.18 | 0.15 | 63.54 | 73.00 | 0.090 | 0.129 | 17.2 | 14.4 | 1214 | 1264 | 182 | 176 |
| M02311470 | 0.025 | 0.023 | 0.072 | 0.074 | 0.082 | 0.096 | 1.11 | 0.81 | 60.86 | 70.78 | 0.083 | 0.116 | 17.8 | 16.5 | 1193 | 1166 | 179 | 173 |
| M02312125 | 0.029 | 0.023 | 0.078 | 0.074 | 0.094 | 0.096 | 0.42 | 1.26 | 65.57 | 64.45 | 0.087 | 0.100 | 17.7 | 16.7 | 1207 | 1159 | 177 | 173 |
| M02315513 | 0.034 | 0.023 | 0.084 | 0.068 | 0.108 | 0.094 | 0.14 | 0.39 | 66.04 | 79.30 | 0.097 | 0.176 | 16.7 | 12.0 | 1222 | 1221 | 182 | 186 |
| M02315514 | 0.036 | 0.025 | 0.082 | 0.068 | 0.096 | 0.094 | 0.77 | 1.17 | 73.87 | 80.04 | 0.122 | 0.166 | 15.9 | 13.0 | 1287 | 1222 | 185 | 182 |
| M02312139 | 0.034 | 0.022 | 0.080 | 0.064 | 0.092 | 0.090 | 0.76 | 0.11 | 64.19 | 74.87 | 0.095 | 0.148 | 16.3 | 13.5 | 1212 | 1233 | 184 | 184 |
| M02330859 | 0.034 | 0.025 | 0.080 | 0.074 | 0.110 | 0.100 | 0.25 | 0.23 | 57.75 | 76.67 | 0.091 | 0.158 | 15.7 | 13.1 | 1146 | 1226 | 185 | 178 |
| M02325311 | 0.028 | 0.021 | 0.072 | 0.064 | 0.098 | 0.086 | 0.43 | 0.05 | 67.82 | 75.77 | 0.099 | 0.138 | 17.3 | 14.8 | 1238 | 1234 | 181 | 174 |
| M02318534 | 0.025 | 0.023 | 0.066 | 0.066 | 0.086 | 0.090 | 0.08 | 0.34 | 55.82 | 64.11 | 0.081 | 0.103 | 15.3 | 15.9 | 1184 | 1165 | 173 | 171 |
| M02318532 | 0.031 | 0.027 | 0.082 | 0.072 | 0.096 | 0.100 | 0.19 | 0.54 | 58.41 | 75.78 | 0.087 | 0.135 | 15.9 | 14.8 | 1198 | 1215 | 176 | 178 |
| M02318531 | 0.038 | 0.031 | 0.090 | 0.092 | 0.108 | 0.112 | 0.69 | 0.09 | 67.09 | 82.78 | 0.106 | 0.179 | 15.8 | 12.3 | 1193 | 1346 | 184 | 189 |
| M02320072 | 0.035 | 0.023 | 0.086 | 0.070 | 0.108 | 0.098 | 0.13 | 0.18 | 56.78 | 78.55 | 0.082 | 0.147 | 16.3 | 14.4 | 1143 | 1333 | 178 | 176 |
| M02337798 | 0.032 | 0.029 | 0.082 | 0.076 | 0.102 | 0.102 | 0.15 | 0.63 | 59.26 | 70.68 | 0.092 | 0.122 | 14.8 | 16.1 | 1163 | 1289 | 179 | 175 |
| M02333893 | 0.033 | 0.022 | 0.086 | 0.066 | 0.106 | 0.096 | 0.17 | 0.06 | 64.10 | 81.03 | 0.096 | 0.169 | 17.5 | 12.4 | 1222 | 1254 | 177 | 179 |
| M02325309 | 0.033 | 0.024 | 0.078 | 0.064 | 0.092 | 0.098 | 0.73 | 1.25 | 64.96 | 83.47 | 0.093 | 0.179 | 17.3 | 11.9 | 1249 | 1243 | 179 | 184 |
| M02333894 | 0.032 | 0.025 | 0.082 | 0.076 | 0.098 | 0.094 | 0.10 | 0.01 | 55.59 | 79.16 | 0.082 | 0.154 | 15.8 | 13.8 | 1145 | 1336 | 179 | 182 |
| M02318535 | 0.033 | 0.026 | 0.076 | 0.074 | 0.090 | 0.098 | 1.55 | 0.94 | 61.57 | 86.25 | 0.092 | 0.212 | 17.0 | 11.1 | 1158 | 1342 | 183 | 186 |
| M02330870 | 0.029 | 0.023 | 0.070 | 0.066 | 0.090 | 0.092 | 3.03 | 0.13 | 70.19 | 84.23 | 0.111 | 0.194 | 16.4 | 11.1 | 1239 | 1196 | 184 | 189 |
| M02337795 | 0.034 | 0.025 | 0.088 | 0.074 | 0.102 | 0.098 | 0.58 | 0.96 | 55.52 | 72.48 | 0.084 | 0.131 | 15.1 | 14.2 | 1183 | 1160 | 173 | 182 |
| M02337796 | 0.036 | 0.030 | 0.086 | 0.082 | 0.104 | 0.106 | 1.85 | 0.39 | 62.58 | 81.13 | 0.101 | 0.174 | 15.6 | 12.6 | 1218 | 1288 | 184 | 183 |
| M02337797 | 0.032 | 0.027 | 0.078 | 0.078 | 0.098 | 0.096 | 1.25 | 1.27 | 63.38 | 75.63 | 0.088 | 0.144 | 17.8 | 13.9 | 1208 | 1217 | 181 | 181 |
| M02320165 | 0.031 | 0.025 | 0.074 | 0.070 | 0.102 | 0.098 | 1.07 | 2.12 | 59.08 | 66.35 | 0.092 | 0.112 | 15.7 | 14.9 | 1214 | 1171 | 173 | 168 |
| M02320166 | 0.031 | 0.023 | 0.074 | 0.064 | 0.086 | 0.090 | 2.66 | 0.01 | 61.84 | 80.80 | 0.090 | 0.164 | 17.0 | 13.1 | 1154 | 1358 | 183 | 176 |
| M02312126 | 0.033 | 0.023 | 0.078 | 0.064 | 0.094 | 0.092 | 1.40 | 0.40 | 69.48 | 80.11 | 0.113 | 0.166 | 16.1 | 12.7 | 1208 | 1322 | 184 | 182 |
| M02339247 | 0.032 | 0.029 | 0.076 | 0.080 | 0.092 | 0.100 | 2.32 | 0.21 | 63.42 | 77.60 | 0.088 | 0.146 | 16.9 | 14.5 | 1178 | 1309 | 181 | 180 |
| M02344734 | 0.036 | 0.028 | 0.088 | 0.078 | 0.102 | 0.104 | 1.58 | 0.62 | 57.54 | 74.59 | 0.087 | 0.140 | 16.4 | 14.2 | 1136 | 1218 | 180 | 184 |
| M02344820 | 0.028 | 0.027 | 0.066 | 0.068 | 0.078 | 0.094 | 0.33 | 0.27 | 60.30 | 77.27 | 0.096 | 0.153 | 16.1 | 13.6 | 1199 | 1182 | 178 | 183 |
| M02344814 | 0.035 | 0.033 | 0.086 | 0.086 | 0.100 | 0.110 | 0.44 | 0.03 | 56.49 | 72.75 | 0.098 | 0.138 | 14.1 | 14.7 | 1168 | 1292 | 181 | 186 |
| M02340377 | 0.028 | 0.023 | 0.070 | 0.070 | 0.086 | 0.090 | 0.40 | 0.04 | 60.11 | 77.28 | 0.087 | 0.146 | 16.8 | 14.2 | 1207 | 1322 | 180 | 171 |
| M02340376 | 0.036 | 0.028 | 0.082 | 0.066 | 0.100 | 0.096 | 0.38 | 1.47 | 66.45 | 82.52 | 0.114 | 0.193 | 15.2 | 11.4 | 1266 | 1166 | 184 | 190 |
| M02333896 | 0.035 | 0.029 | 0.078 | 0.076 | 0.112 | 0.106 | 0.43 | 0.77 | 56.14 | 68.24 | 0.085 | 0.123 | 16.3 | 15.0 | 1167 | 1185 | 177 | 183 |
| M02344819 | 0.033 | 0.030 | 0.080 | 0.080 | 0.100 | 0.102 | 0.65 | 0.22 | 60.16 | 81.44 | 0.088 | 0.173 | 15.7 | 12.9 | 1167 | 1364 | 182 | 189 |
| M02344818 | 0.030 | 0.023 | 0.072 | 0.062 | 0.086 | 0.092 | 0.63 | 0.91 | 63.54 | 80.77 | 0.099 | 0.169 | 15.6 | 12.4 | 1260 | 1201 | 174 | 183 |
| M02344733 | 0.029 | 0.020 | 0.074 | 0.064 | 0.098 | 0.088 | 0.16 | 1.47 | 51.22 | 70.29 | 0.072 | 0.113 | 15.9 | 16.2 | 1135 | 1305 | 170 | 173 |
| M02339248 | 0.037 | 0.030 | 0.080 | 0.080 | 0.094 | 0.096 | 2.10 | 0.68 | 64.05 | 81.55 | 0.094 | 0.183 | 16.4 | 11.7 | 1225 | 1291 | 177 | 188 |
| M02339246 | 0.029 | 0.023 | 0.072 | 0.076 | 0.086 | 0.096 | 1.90 | 0.17 | 60.77 | 80.06 | 0.098 | 0.178 | 15.0 | 12.1 | 1245 | 1231 | 180 | 183 |
| M02340378 | 0.031 | 0.022 | 0.074 | 0.060 | 0.100 | 0.086 | 1.70 | 0.69 | 67.59 | 84.77 | 0.106 | 0.184 | 16.5 | 11.5 | 1259 | 1357 | 183 | 187 |
| M02344815 | 0.030 | 0.023 | 0.076 | 0.062 | 0.094 | 0.082 | 0.35 | 1.35 | 64.01 | 64.55 | 0.093 | 0.099 | 16.8 | 16.4 | 1257 | 1232 | 179 | 176 |
| M02342848 | 0.028 | 0.025 | 0.066 | 0.060 | 0.078 | 0.086 | 0.60 | 0.16 | 56.53 | 58.34 | 0.080 | 0.086 | 16.4 | 15.6 | 1186 | 1091 | 175 | 172 |
| M02342849 | 0.035 | 0.032 | 0.088 | 0.074 | 0.102 | 0.102 | 1.40 | 0.61 | 56.09 | 66.87 | 0.085 | 0.116 | 15.4 | 14.9 | 1199 | 1137 | 179 | 187 |
| M02342855 | 0.033 | 0.026 | 0.080 | 0.072 | 0.094 | 0.094 | 0.89 | 0.11 | 67.81 | 82.27 | 0.098 | 0.190 | 17.3 | 11.5 | 1240 | 1273 | 182 | 174 |
| M02342846 | 0.031 | 0.025 | 0.076 | 0.078 | 0.102 | 0.104 | 1.05 | 0.22 | 55.52 | 76.30 | 0.078 | 0.140 | 15.8 | 14.7 | 1133 | 1320 | 175 | 178 |
| M02342857 | 0.029 | 0.025 | 0.076 | 0.076 | 0.098 | 0.104 | 0.47 | 0.28 | 56.40 | 73.57 | 0.080 | 0.121 | 15.2 | 15.8 | 1161 | 1294 | 175 | 184 |
| M02350234 | 0.035 | 0.023 | 0.082 | 0.066 | 0.094 | 0.092 | 0.20 | 0.09 | 55.62 | 66.43 | 0.074 | 0.103 | 16.2 | 16.4 | 1135 | 1290 | 174 | 180 |
| M02350237 | 0.027 | 0.022 | 0.074 | 0.070 | 0.088 | 0.100 | 0.82 | 0.02 | 54.88 | 64.43 | 0.074 | 0.094 | 15.9 | 16.2 | 1137 | 1258 | 176 | 176 |
| M02345222 | 0.029 | 0.022 | 0.070 | 0.070 | 0.094 | 0.088 | 0.39 | 0.07 | 49.95 | 68.89 | 0.070 | 0.106 | 15.2 | 16.8 | 1095 | 1280 | 174 | 176 |
| M02350233 | 0.026 | 0.024 | 0.066 | 0.072 | 0.082 | 0.092 | 0.53 | 0.01 | 59.84 | 70.13 | 0.078 | 0.117 | 17.6 | 16.0 | 1179 | 1161 | 181 | 180 |
| M02345220 | 0.030 | 0.027 | 0.068 | 0.068 | 0.090 | 0.088 | 0.17 | 0.01 | 61.50 | 73.84 | 0.095 | 0.138 | 16.6 | 13.9 | 1219 | 1201 | 179 | 189 |
| M02350231 | 0.032 | 0.023 | 0.080 | 0.068 | 0.092 | 0.092 | 0.17 | 0.13 | 56.77 | 85.17 | 0.083 | 0.176 | 15.3 | 12.5 | 1125 | 1366 | 179 | 182 |
| M02345221 | 0.036 | 0.028 | 0.086 | 0.080 | 0.102 | 0.102 | 0.77 | 0.09 | 63.65 | 82.85 | 0.099 | 0.179 | 15.7 | 12.0 | 1220 | 1237 | 183 | 187 |
| M02341339 | 0.033 | 0.029 | 0.080 | 0.076 | 0.094 | 0.106 | 0.28 | 0.19 | 61.35 | 76.71 | 0.088 | 0.138 | 16.6 | 14.2 | 1224 | 1191 | 182 | 181 |
| M02350236 | 0.034 | 0.023 | 0.086 | 0.070 | 0.100 | 0.092 | 0.13 | 0.17 | 57.96 | 85.68 | 0.087 | 0.191 | 15.1 | 12.0 | 1143 | 1395 | 176 | 184 |
| M02341338 | 0.032 | 0.029 | 0.078 | 0.076 | 0.100 | 0.102 | 0.55 | 0.11 | 60.12 | 67.08 | 0.087 | 0.123 | 17.5 | 14.6 | 1211 | 1203 | 181 | 182 |
| M02351498 | 0.032 | 0.027 | 0.074 | 0.070 | 0.086 | 0.100 | 1.21 | 0.05 | 70.74 | 83.59 | 0.111 | 0.179 | 16.7 | 12.0 | 1266 | 1188 | 183 | 184 |
| M02340379 | 0.037 | 0.029 | 0.086 | 0.078 | 0.102 | 0.106 | 1.11 | 0.26 | 61.19 | 88.05 | 0.097 | 0.228 | 15.5 | 10.2 | 1157 | 1373 | 185 | 190 |
| M02364046 | 0.033 | 0.022 | 0.074 | 0.066 | 0.088 | 0.096 | 0.09 | 0.15 | 69.91 | 85.53 | 0.105 | 0.184 | 16.6 | 11.7 | 1258 | 1165 | 182 | 190 |
| M02364047 | 0.031 | 0.021 | 0.076 | 0.058 | 0.090 | 0.084 | 0.60 | 0.07 | 64.16 | 81.44 | 0.096 | 0.151 | 17.1 | 13.5 | 1235 | 1281 | 183 | 181 |
| M02380072 | 0.039 | 0.028 | 0.084 | 0.072 | 0.104 | 0.102 | 0.89 | 0.75 | 69.08 | 83.97 | 0.109 | 0.186 | 16.4 | 11.5 | 1237 | 1197 | 191 | 194 |
| M02382493 | 0.031 | 0.022 | 0.072 | 0.068 | 0.088 | 0.094 | 0.57 | 0.02 | 61.61 | 75.06 | 0.094 | 0.145 | 16.7 | 13.2 | 1233 | 1193 | 187 | 175 |
| M02382494 | 0.031 | 0.030 | 0.076 | 0.080 | 0.096 | 0.102 | 0.72 | 0.80 | 70.98 | 85.87 | 0.107 | 0.204 | 16.7 | 10.8 | 1244 | 1190 | 190 | 191 |
| M02376923 | 0.035 | 0.031 | 0.082 | 0.076 | 0.100 | 0.102 | 0.21 | 0.35 | 68.61 | 87.75 | 0.107 | 0.198 | 16.3 | 11.1 | 1212 | 1368 | 182 | 191 |
| M02376932 | 0.036 | 0.026 | 0.082 | 0.072 | 0.102 | 0.092 | 1.37 | 0.00 | 67.90 | 86.10 | 0.110 | 0.205 | 15.3 | 10.8 | 1189 | 1381 | 181 | 181 |
| M02406881 | 0.031 | 0.027 | 0.076 | 0.074 | 0.096 | 0.094 | 1.33 | 0.56 | 61.89 | 85.32 | 0.093 | 0.185 | 16.5 | 11.7 | 1134 | 1352 | 186 | 191 |
| M02487371 | 0.030 | 0.026 | 0.072 | 0.070 | 0.088 | 0.092 | 1.61 | 0.15 | 61.82 | 76.98 | 0.096 | 0.145 | 16.2 | 13.6 | 1211 | 1176 | 182 | 187 |
| M02311619 | 0.037 | 0.027 | 0.086 | 0.072 | 0.102 | 0.094 | 0.48 | 1.00 | 72.28 | 81.86 | 0.117 | 0.181 | 16.8 | 12.0 | 1252 | 1205 | 185 | 189 |
| M02311621 | 0.037 | 0.028 | 0.084 | 0.066 | 0.104 | 0.098 | 0.52 | 0.72 | 74.54 | 86.64 | 0.120 | 0.238 | 15.8 | 9.6 | 1250 | 1199 | 194 | 189 |
| M02334479 | 0.033 | 0.038 | 0.074 | 0.096 | 0.094 | 0.118 | 0.85 | 1.09 | 67.27 | 78.18 | 0.108 | 0.160 | 16.7 | 13.1 | 1219 | 1198 | 193 | 184 |
| M02319155 | 0.035 | 0.031 | 0.086 | 0.078 | 0.102 | 0.104 | 1.21 | 0.26 | 65.30 | 71.94 | 0.095 | 0.140 | 17.6 | 13.7 | 1197 | 1221 | 187 | 179 |
| M02319158 | 0.031 | 0.036 | 0.074 | 0.090 | 0.104 | 0.126 | 1.55 | 0.48 | 63.60 | 72.57 | 0.087 | 0.143 | 18.8 | 13.9 | 1162 | 1193 | 188 | 182 |
| M02334480 | 0.034 | 0.023 | 0.078 | 0.060 | 0.096 | 0.088 | 0.29 | 0.10 | 66.92 | 87.73 | 0.111 | 0.232 | 15.6 | 10.1 | 1233 | 1251 | 190 | 186 |
| M02355075 | 0.032 | 0.027 | 0.078 | 0.070 | 0.098 | 0.096 | 2.53 | 0.42 | 69.58 | 83.35 | 0.112 | 0.198 | 16.1 | 11.3 | 1247 | 1224 | 194 | 190 |
| M02364713 | 0.033 | 0.025 | 0.076 | 0.072 | 0.098 | 0.096 | 1.35 | 0.26 | 65.11 | 88.23 | 0.117 | 0.238 | 14.9 | 9.5 | 1181 | 1330 | 192 | 187 |
| M02364715 | 0.032 | 0.025 | 0.080 | 0.072 | 0.104 | 0.102 | 0.72 | 1.35 | 69.16 | 84.51 | 0.107 | 0.196 | 16.2 | 11.0 | 1232 | 1174 | 190 | 187 |
| M02374611 | 0.035 | 0.027 | 0.084 | 0.074 | 0.096 | 0.100 | 1.14 | 0.64 | 62.49 | 80.07 | 0.101 | 0.165 | 16.0 | 13.1 | 1190 | 1195 | 193 | 183 |
| M02374613 | 0.034 | 0.030 | 0.080 | 0.080 | 0.098 | 0.104 | 0.82 | 1.22 | 68.09 | 78.15 | 0.110 | 0.164 | 16.6 | 13.0 | 1217 | 1196 | 189 | 190 |
| M02374615 | 0.030 | 0.024 | 0.072 | 0.066 | 0.084 | 0.096 | 2.34 | 2.58 | 57.50 | 86.01 | 0.086 | 0.203 | 14.8 | 10.8 | 1153 | 1269 | 181 | 186 |
| Skewness | -0.13 | 0.75 | -0.28 | 0.62 | -0.28 | 0.81 | 1.28 | 1.31 | -0.17 | -0.71 | 0.03 | -0.02 | -0.01 | 0.22 | -0.32 | 0.24 | 0.23 | -0.37 |
| Kurtosis | 0.37 | 0.94 | -0.63 | 1.12 | -0.40 | 2.32 | 1.43 | 1.51 | -0.17 | -0.71 | -0.55 | -0.68 | 0.63 | -0.82 | -0.41 | -0.84 | -0.19 | -0.47 |
| Shapiro-Wilk normality test | | | | | | | | | | | | | | | | | | |
| W | 0.9944 | 0.9587 | 0.9662 | 0.9655 | 0.9783 | 0.9560 | 0.8846 | 0.8537 | 0.9878 | 0.9374 | 0.9883 | 0.9829 | 0.9897 | 0.9705 | 0.9815 | 0.9660 | 0.9881 | 0.9750 |
| P value | 0.9565 | 0.0033 | 0.0115 | 0.0102 | 0.0987 | 0.0021 | <0.0001 | <0.0001 | 0.4964 | 0.0001 | 0.5286 | 0.2203 | 0.6428 | 0.0241 | 0.1730 | 0.0111 | 0.5150 | 0.0539 |
| Passed normality test (alpha=0.05)? | Yes | No | No | No | Yes | No | No | No | Yes | No | Yes | Yes | Yes | No | Yes | No | Yes | Yes |
| P value summary | ns | ** | * | * | ns | ** | **** | **** | ns | *** | ns | ns | ns | * | ns | * | ns | ns |
| Distribution: | Normal | Non-normal | Non-normal | Non-normal | Normal | Non-normal | Non-normal | Non-normal | Normal | Non-normal | Normal | Normal | Normal | Non-normal | Normal | Non-normal | Normal | Normal |
| Normally-distributed parameters | | | | | | | | | | | | | | | | | | |
| Mean | 0.032 | | | | 0.096 | | | | 63.20 | | 0.096 | 0.163 | 16.2 | | 1203 | | 182.4 | 182.9 |
| Std. Deviation | 0.004 | | | | 0.007 | | | | 5.01 | | 0.01 | 0.037 | 0.841 | | 39 | | 5.3 | 5.6 |
| Lower 95% CI of mean | 0.032 | | | | 0.095 | | | | 62.2 | | 0.094 | 0.156 | 16.1 | | 1195 | | 181.3 | 181.7 |
| Upper 95% CI of mean | 0.033 | | | | 0.098 | | | | 64.2 | | 0.099 | 0.171 | 16.4 | | 1211 | | 183.4 | 184 |
| mean -2SD | 0.025 | | | | 0.081 | | | | 53.2 | | 0.073 | 0.089 | 14.5 | | 1124 | | 171.7 | 171.6 |
| mean + S2D | 0.040 | | | | 0.111 | | | | 73.2 | | 0.120 | 0.238 | 17.9 | | 1282 | | 193.1 | 194.1 |
| Coefficient of variation | 11.0% | | | | 7.8% | | | | 7.9% | | 12.0% | 22.8% | 5.2% | | 3.3% | | 2.9% | 3.1% |
| n for 80% power to detect a 2SD delta | 4 | | | | 4 | | | | 4 | | 4 | 4 | 4 | | 4 | | 4 | 4 |
| n for 90% power to detect a 2SD delta | 6 | | | | 6 | | | | 6 | | 6 | 6 | 6 | | 6 | | 6 | 6 |
| Non-normally-distributed parameters | | | | | | | | | | | | | | | | | | |
| Median | | 0.0253 | 0.0800 | 0.0720 | | 0.0960 | 0.7197 | 0.3683 | | 80.1 | | | | 13.00 | | 1240 | | |
| Lower limit of 95% CI of median | | 0.0248 | 0.0780 | 0.0700 | | 0.0960 | 0.5700 | 0.2200 | | 77.6 | | | | 12.20 | | 1221 | | |
| Upper limit of 95% CI of median | | 0.0266 | 0.0820 | 0.0740 | | 0.0980 | 0.8800 | 0.5600 | | 81.6 | | | | 13.50 | | 1269 | | |
| 2.5% Percentile | | 0.0204 | 0.0660 | 0.0600 | | 0.0851 | 0.0953 | 0.0100 | | 63.9 | | | | 9.71 | | 1136 | | |
| 97.5% Percentile | | 0.0342 | 0.0890 | 0.0910 | | 0.1149 | 2.8360 | 2.2060 | | 87.9 | | | | 16.60 | | 1378 | | |
| Precision | | 0.0069 | 0.0115 | 0.0155 | | 0.0149 | 1.3704 | 1.0980 | | 12.02 | | | | 3.4475 | | 121.0000 | | |
| Coefficient of variation | | 27.3% | 14.3% | 21.5% | | 15.5% | 190.4% | 298.1% | | 15.0% | | | | 26.5% | | 9.8% | | |
| Median of absolute deviations from the median (MAD) | | **0.0024** | **0.0040** | **0.0040** | | **0.0040** | **0.3733** | **0.2788** | | **4.5750** | | | | **1.4429** | | **49.7910** | | |
| Percentage MAD (MAD/median)% | | **9.50%** | **5.00%** | **5.56%** | | **4.17%** | **51.87%** | **75.71%** | | **5.71%** | | | | **11.10%** | | **4.01%** | | |
| n to detect outside 95% CI | | **3** | **3** | **1** | | **2** | **2** | **1** | | **3** | | | | **4** | | **4** | | |

**Repeatability of measurement of joint phenotyping parameters.**

| Tissue | Parameter | Unit | Method | Plateau | Sample 1 | | | Sample 2 | | | Sample 3 | | | Sample 4 | | | Sample 5 | | | Sample 6 | | | Sample 7 | | |
|---|---|---|---|---|---|---|---|---|---|---|---|---|---|---|---|---|---|---|---|---|---|---|---|---|---|
| | | | | | Mean | SEM | CV | Mean | SEM | CV | Mean | SEM | CV | Mean | SEM | CV | Mean | SEM | CV | Mean | SEM | CV | Mean | SEM | CV |
| Articular cartilage | ACV | ($mm^3$) | ICE-µCT | LTP | 0.031 | 0.0002 | 1.7% | 0.034 | 0.0002 | 1.6% | 0.027 | 0.0002 | 1.4% | 0.038 | 0.0001 | 0.9% | 0.037 | 0.0001 | 0.8% | 0.031 | 0.0002 | 1.8% | 0.029 | 0.0001 | 0.8% |
| | | | | MTP | 0.022 | 0.0002 | 2.4% | 0.024 | 0.0002 | 1.9% | 0.020 | 0.0001 | 1.3% | 0.034 | 0.0002 | 1.6% | 0.025 | 0.0004 | 3.1% | 0.027 | 0.0001 | 0.9% | 0.028 | 0.0002 | 1.6% |
| | Median AC Th | (mm) | ICE-µCT | LTP | 0.076 | 0.0000 | 0.0% | 0.082 | 0.0004 | 1.1% | 0.072 | 0.0000 | 0.0% | 0.088 | 0.0000 | 0.0% | 0.084 | 0.0000 | 0.0% | 0.075 | 0.0005 | 1.5% | 0.066 | 0.0000 | 0.0% |
| | | | | MTP | 0.067 | 0.0005 | 1.6% | 0.067 | 0.0005 | 1.6% | 0.056 | 0.0007 | 3.0% | 0.090 | 0.0000 | 0.0% | 0.066 | 0.0010 | 3.3% | 0.074 | 0.0004 | 1.2% | 0.067 | 0.0005 | 1.6% |
| | Maximum AC Th | (mm) | ICE-µCT | LTP | 0.094 | 0.0000 | 0.0% | 0.102 | 0.0000 | 0.0% | 0.098 | 0.0000 | 0.0% | 0.108 | 0.0000 | 0.0% | 0.107 | 0.0005 | 1.0% | 0.098 | 0.0000 | 0.0% | 0.078 | 0.0000 | 0.0% |
| | | | | MTP | 0.091 | 0.0005 | 1.2% | 0.091 | 0.0005 | 1.2% | 0.086 | 0.0004 | 1.0% | 0.111 | 0.0005 | 1.0% | 0.094 | 0.0000 | 0.0% | 0.094 | 0.0000 | 0.0% | 0.094 | 0.0000 | 0.0% |
| | AC Damage Area | (%) | JSR | LTP | 0.45 | 0.0795 | 39.5% | 1.302 | 0.1511 | 26.0% | 0.416 | 0.0790 | 42.5% | 0.976 | 0.1352 | 31.0% | 0.896 | 0.0327 | 8.1% | 0.056 | 0.0236 | 94.2% | 0.38 | 0.0283 | 16.6% |
| | | | | MTP | 1.414 | 0.0924 | 14.6% | 0.52 | 0.0182 | 7.8% | 0.008 | 0.0020 | 55.9% | 0.206 | 0.0393 | 42.7% | 1.042 | 0.1593 | 34.2% | 0.04 | 0.0155 | 86.6% | 0.322 | 0.0648 | 45.0% |
| Subchondral bone | SC BV/TV | (%) | ICE-µCT | LTP | 63.8 | 0.5128 | 1.8% | 63.02 | 0.8034 | 2.9% | 67.88 | 0.1356 | 0.5% | 66.28 | 0.3513 | 1.2% | 64.64 | 0.2159 | 0.8% | 55.38 | 0.4140 | 1.7% | 62.44 | 0.1536 | 0.6% |
| | | | | MTP | 85.4 | 0.3421 | 0.9% | 84.92 | 1.1440 | 3.0% | 79.12 | 0.7749 | 2.2% | 85.76 | 0.3614 | 0.9% | 84.36 | 0.5446 | 1.4% | 81.08 | 0.5544 | 1.5% | 77.46 | 0.3326 | 1.0% |
| | SC Tb.Th | (mm) | ICE-µCT | LTP | 0.095 | 0.0002 | 0.6% | 0.098 | 0.0003 | 0.7% | 0.103 | 0.0004 | 0.8% | 0.104 | 0.0002 | 0.5% | 0.103 | 0.0005 | 1.0% | 0.082 | 0.0001 | 0.2% | 0.095 | 0.0001 | 0.3% |
| | | | | MTP | 0.180 | 0.0017 | 2.1% | 0.198 | 0.0031 | 3.5% | 0.150 | 0.0012 | 1.8% | 0.181 | 0.0007 | 0.9% | 0.195 | 0.0007 | 0.8% | 0.150 | 0.0006 | 0.9% | 0.153 | 0.0005 | 0.7% |
| | SC Tb.N | | ICE-µCT | LTP | 17.2 | 0.0859 | 1.1% | 16.5 | 0.1246 | 1.7% | 17.6 | 0.0197 | 0.3% | 16.5 | 0.1004 | 1.4% | 16.6 | 0.0376 | 0.5% | 16.1 | 0.1472 | 2.1% | 17.1 | 0.0681 | 0.9% |
| | | | | MTP | 12.2 | 0.0838 | 1.5% | 11.6 | 0.1972 | 3.8% | 13.8 | 0.1070 | 1.7% | 12.3 | 0.0491 | 0.9% | 11.1 | 0.0556 | 1.1% | 13.9 | 0.0839 | 1.3% | 13.5 | 0.0328 | 0.5% |
| | SC BMD | (mg HA/$cm^3$) | ICE-µCT | LTP | 1206 | 1.0170 | 0.2% | 1165 | 1.2080 | 0.2% | 1236 | 0.6126 | 0.1% | 1187 | 0.8015 | 0.2% | 1219 | 0.4631 | 0.1% | 1133 | 0.4359 | 0.1% | 1195 | 0.4437 | 0.1% |
| | | | | MTP | 1362 | 2.2210 | 0.4% | 1378 | 5.3950 | 0.9% | 1246 | 1.4950 | 0.3% | 1344 | 0.9753 | 0.2% | 1201 | 0.6305 | 0.1% | 1336 | 2.8010 | 0.5% | 1181 | 0.7551 | 0.1% |
| | SC BMC | (median grey level) | sc-XRM | LTP | 176.4 | 0.4000 | 0.5% | 174.8 | 0.4899 | 0.6% | 180.1 | 0.6782 | 0.8% | 185.1 | 0.4000 | 0.5% | 187.2 | 0.7348 | 0.9% | 179.7 | 0.3000 | 0.4% | 180.5 | 0.2236 | 0.3% |
| | | | | MTP | 178.6 | 0.2449 | 0.3% | 171.8 | 0.2000 | 0.3% | 173.8 | 0.2000 | 0.3% | 189.2 | 0.2000 | 0.2% | 192.2 | 0.2000 | 0.2% | 182.2 | 0.2000 | 0.2% | 185.2 | 0.5831 | 0.7% |

**Joint phenotyping pipeline data from 50 unselected mouse lines.**

| | SD from population mean, or percentile | | | Tissue compartment: | Articular cartilage | | | | | | | | Subchondral bone | | | | | | | | | |
|---|---|---|---|---|---|---|---|---|---|---|---|---|---|---|---|---|---|---|---|---|---|---|
| | | | | Parameter: | ACV | | Median AC Th | | Maximum AC Th | | AC Damage Area | | SC BV/TV | | SC Tb.Th | | SC Tb.N | | SC BMD | | SC BMC | |
| | | | | Unit: | ($mm^3$) | | (mm) | | (mm) | | (%) | | (%) | | (mm) | | | | (mg HA/$cm^3$) | | (median grey level) | |
| | | | | Technique: | ICE-μCT | | ICE-μCT | | ICE-μCT | | JSR | | ICE-μCT | | ICE-μCT | | ICE-μCT | | ICE-μCT | | scXRM | |
| | | | | Reference range: | mean ± 2SD | 2.5–97.5% | 2.5–97.5% | 2.5–97.5% | mean ± 2SD | 2.5–97.5% | 2.5–97.5% | 2.5–97.5% | mean ± 2SD | 2.5–97.5% | mean ± 2SD | mean ± 2SD | mean ± 2SD | 2.5-97.5% | mean ± 2SD | 2.5–97.5% | mean ± 2SD | mean ± 2SD |
| Line | Knockout Allele | Genotype | Samples (n) | Gene | LTP | MTP | LTP | MTP | LTP | MTP | LTP | MTP | LTP | MTP | LTP | MTP | LTP | MTP | LTP | MTP | LTP | MTP |
| 1 | 4932431P20Rik<em1(IMPC)Wtsi> | HOM | 6 | 4932431P20Rik | -0.92 | 38.2% | 10.3% | 14.6% | -0.08 | 9.2% | 91.0% | 57.7% | -0.29 | 11.7% | -0.58 | -1.19 | 0.16 | 84.2% | 0.04 | 48.2% | -0.79 | -0.60 |
| 2 | A430078G23Rik<tm1a(KOMP)Wtsi> | HOM | 3 | Arhgef18 | 0.06 | 84.1% | 48.8% | 86.7% | 0.32 | 80.5% | 96.3% | 50.2% | -0.39 | 23.8% | -0.31 | -0.68 | -0.20 | 72.5% | -0.96 | 6.5% | -0.13 | 0.44 |
| **3** | **Arhgap30<tm1a(EUCOMM)Wtsi>** | **HOM** | **3** | **Arhgap30** | **0.13** | **85.6%** | **18.1%** | **66.3%** | **-1.10** | **80.5%** | **98.0%** | **83.0%** | **0.62** | **39.5%** | **1.03** | **0.17** | **0.57** | **42.4%** | **-1.60** | **11.5%** | **0.43** | **-0.39** |
| **4** | **Arrdc5<tm1b(EUCOMM)Wtsi>** | **HOM** | **4** | **Arrdc5** | 0.22 | 9.9% | 39.1% | <0% | -0.30 | 5.1% | 97.4% | 79.2% | 0.41 | 48.0% | 0.33 | 0.23 | 0.16 | 40.4% | 0.85 | 50.2% | 0.21 | -1.09 |
| **5** | **Bhlhe40<tm1b(KOMP)Wtsi>** | **HOM** | **3** | **Bhlhe40** | -1.96 | 4.9% | <0% | **1.6%** | -3.06 | <0% | 91.2% | 81.8% | 0.86 | 47.9% | 0.62 | -0.35 | 0.56 | 68.3% | -0.46 | 74.0% | 0.74 | -0.03 |
| **6** | **Ccdc6<em1(IMPC)Wtsi>** | **HET** | **4** | **Ccdc6** | -0.17 | 30.6% | 49.5% | 14.5% | -0.21 | 15.5% | 89.3% | **98.8%** | 0.35 | 44.4% | -0.14 | 0.01 | 0.90 | 47.8% | 0.70 | 51.4% | 0.12 | -0.21 |
| 7 | Cfap53<em1(IMPC)Wtsi> | HET | 6 | Cfap53 | -0.11 | 58.0% | 27.7% | 14.8% | -0.48 | 26.5% | 91.7% | 48.3% | 0.69 | 37.1% | 0.53 | -0.42 | 0.27 | 63.3% | 0.87 | 33.9% | 0.43 | 0.03 |
| **8** | **Chka<tm2a(KOMP)Wtsi>** | **HET** | **4** | **Chka** | 0.75 | 72.5% | 73.2% | 67.0% | 0.63 | 60.8% | **97%** | 83.5% | 0.57 | 26.2% | 0.52 | -0.09 | 0.16 | 46.5% | 0.90 | 42.1% | 1.05 | 0.60 |
| 9 | ENSMUSG00000065619<(tm1)Brd> | HOM | 3 | Clust6N1 | 0.49 | 60.7% | 59.7% | 44.8% | 0.68 | 36.3% | 92.2% | 66.3% | -0.28 | 38.9% | -0.31 | -0.31 | 0.40 | 65.7% | -0.69 | 43.9% | 0.12 | -0.63 |
| **10** | **ClusterXN1<tm1(Brd)>** | **HEMI** | **3** | **ClusterXN1** | **-0.69** | **76.2%** | **27.0%** | **33.3%** | **-0.30** | **25.7%** | **>100%** | **71.2%** | **0.32** | **27.8%** | **0.23** | **-0.66** | **0.81** | **73.1%** | **-1.50** | **2.90%** | **-0.38** | **-0.21** |
| **11** | **Col4a2<em1(IMPC)Wtsi>** | **HET** | **3** | **Col4a2** | -0.54 | 64.3% | 39.2% | 77.5% | -0.21 | 61.0% | **>100%** | 44.5% | -0.36 | 74.0% | 0.01 | 0.57 | -0.22 | 29.7% | 0.30 | 60.8% | 0.12 | 0.56 |
| **12** | **Cpgi81<tm1.1(NCC)WCS>** | **HOM** | **5** | **Cpgi81** | **-0.45** | **70.6%** | **18.6%** | **53.6%** | **-0.52** | **61.1%** | **96.70%** | **93.2%** | **0.05** | **29.4%** | **-0.50** | **-0.81** | **0.56** | **77.5%** | **-0.68** | **38.3%** | **-1.16** | **-0.65** |
| 13 | Dctn4<em1(IMPC)Wtsi> | HET | 6 | Dctn4 | -0.65 | 38.2% | 38.7% | 24.7% | -0.03 | 15.6% | 89.4% | 75.1% | -0.70 | 28.3% | -0.68 | -0.43 | -0.19 | 58.2% | -0.12 | 58.7% | -0.76 | 0.12 |
| 14 | Deptor<tm1b(EUCOMM)Wtsi> | HOM | 3 | Deptor | -0.35 | 39.6% | 7.9% | 24.0% | -1.28 | 52.4% | 58.2% | 8.6% | -0.60 | 37.3% | -0.59 | -0.17 | 0.34 | 47.9% | -0.72 | 47.7% | 0.62 | -0.74 |
| 15 | Dnah17<tm1e(KOMP)Wtsi> | HOM | 3 | Dnah17 | -0.38 | 78.7% | 27.2% | 53.9% | -0.30 | 36.6% | 81.6% | 42.4% | -0.27 | 18.2% | -0.27 | -0.96 | -0.10 | 69.6% | 0.59 | 15.7% | -0.40 | -0.06 |
| 16 | Washc2<tm2b(KOMP)Wtsi> | HET | 4 | Fam21 | 1.03 | 38.5% | 79.7% | 53.4% | 0.83 | 80.6% | 90.9% | 43.6% | 0.80 | 73.2% | 0.63 | 0.51 | -0.17 | 27.5% | 1.15 | 50.9% | 1.29 | 0.20 |
| **17** | **Fgfbp1<em1(IMPC)Wtsi>** | **HOM** | **4** | **Fgfbp1** | **-0.46** | **32.5%** | **49.0%** | **14.3%** | **-0.03** | **52.9%** | **81.8%** | **24.7%** | **0.42** | **60.6%** | **-0.54** | **-0.12** | **1.74** | **48.4%** | **0.84** | **18.3%** | **-2.08** | **-1.04** |
| 18 | Gm6578<tm1a(KOMP)Wtsi> | HOM | 3 | Gm6578 | 0.69 | 62.3% | 60.0% | 54.1% | 0.50 | 60.7% | 78.8% | 69.5% | 0.70 | 44.2% | 1.28 | 0.04 | -0.52 | 44.9% | 0.08 | 49.4% | 1.18 | 0.56 |
| 19 | Gpbp1l1<tm1a(EUCOMM)Wtsi> | HOM | 3 | Gpbp1l1 | 1.15 | 84.1% | 92.4% | 87.1% | 1.57 | 95.0% | 72.5% | 25.6% | -0.29 | 29.1% | 0.25 | -0.02 | -0.71 | 46.5% | -0.68 | 34.0% | 0.05 | 0.74 |
| **20** | **Grsf1<tm1b(EUCOMM)Wtsi>** | **HOM** | **3** | **Grsf1** | **0.04** | **90.5%** | **48.8%** | **87.1%** | **0.59** | **94.7%** | **76.5%** | **95.5%** | **1.49** | **50.4%** | **1.43** | **0.20** | **0.06** | **41.9%** | **0.41** | **52.0%** | **-0.45** | **-0.63** |
| **21** | **Gsdme<tm1b(KOMP)Wtsi>** | **HOM** | **4** | **Gsdme** | -0.25 | 53.9% | 27.7% | 45.0% | 0.10 | 53.2% | 97.5% | 60.6% | 0.59 | 43.5% | 0.19 | -0.26 | 0.31 | 58.7% | 0.68 | 66.9% | 0.16 | -0.15 |
| 22 | Hao2<em1(IMPC)Wtsi> | HOM | 3 | Hao2 | -1.23 | 55.7% | 7.9% | 54.1% | -1.28 | 52.8% | 88.1% | 42.0% | -0.42 | 33.7% | -0.51 | -0.46 | -0.66 | 64.3% | 0.24 | 51.4% | -0.88 | 0.09 |
| **23** | **Herc1<em2(IMPC)Wtsi>** | **HOM** | **4** | **Herc1** | -1.27 | 36.7% | **5.4%** | 6.6% | -1.24 | 9.6% | 73.4% | 12.3% | 0.01 | 17.4% | -0.55 | -0.96 | 0.61 | 72.2% | 0.12 | 9.9% | -0.26 | 0.43 |
| 24 | Hmgxb3<tm1a(EUCOMM)Wtsi> | HET | 4 | Hmgxb3 | -0.32 | 54.0% | 27.2% | 33.4% | -0.44 | 36.1% | 56.1% | 24.9% | -0.82 | 43.5% | -0.92 | 0.01 | -0.11 | 59.3% | -0.58 | 76.9% | -0.40 | -0.73 |
| **25** | **Hpf1<em1(IMPC)Wtsi>** | **HOM** | **5** | **Hpf1** | **-0.67** | **11.5%** | **27.7%** | **6.3%** | **0.18** | **15.4%** | **81.5%** | **62.2%** | **0.40** | **82.8%** | **0.13** | **1.12** | **0.82** | **13.2%** | **0.69** | **47.9%** | **0.49** | **0.99** |
| **26** | **Josd1<em1(IMPC)Wtsi>** | **HOM** | **6** | **Josd1** | -0.34 | 37.6% | **10.5%** | **14.1%** | **-1.24** | 15.5% | 97.6% | 77.5% | 0.76 | 60.7% | 0.39 | 0.49 | 0.43 | 38.5% | 0.91 | 43.1% | -0.07 | 0.20 |
| 27 | Lpxn<tm1b(EUCOMM)Hmgu> | HOM | 3 | Lpxn | -0.13 | 22.1% | 39.6% | 33.6% | -0.39 | 36.3% | 82.2% | 48.0% | -0.40 | 47.2% | -0.08 | -0.06 | -0.71 | 56.8% | -0.89 | 64.7% | 0.43 | 0.09 |
| 28 | Lrrc8d<tm1a(EUCOMM)Wtsi> | HOM | 3 | Lrrc8d | 0.62 | 38.1% | 60.0% | 14.8% | 0.05 | 36.3% | 81.4% | 11.4% | 0.29 | 61.1% | 0.60 | 0.39 | -0.75 | 39.0% | -0.71 | 73.7% | 0.55 | -0.03 |
| 29 | Mbd1<em1(IMPC)Wtsi> | HOM | 4 | Mbd1 | 1.10 | 71.4% | 72.7% | 54.2% | 0.50 | 53.2% | 96.0% | 50.9% | -0.15 | 37.2% | -0.10 | -0.12 | -0.04 | 57.0% | 0.49 | 43.0% | -0.49 | 0.12 |
| **30** | **Mkrn2<em1(IMPC)Wtsi>** | **HOM** | **4** | **Mkrn2** | -0.85 | 47.0% | 18.0% | 33.4% | -1.44 | 25.9% | 97.6% | 81.8% | 0.79 | 69.2% | 0.27 | 0.25 | -0.03 | 42.3% | 0.39 | 52.0% | 0.40 | 0.47 |
| 31 | Mroh9<tm1a(EUCOMM)Wtsi> | HOM | 3 | Mroh9 | 0.43 | 37.4% | 49.5% | 33.0% | -0.48 | 35.9% | 64.7% | 26.0% | -0.24 | 55.4% | -0.63 | 0.23 | 0.47 | 46.8% | -0.64 | 58.0% | -0.51 | 0.74 |
| **32** | **Nebl<tm1b(EUCOMM)Wtsi>** | **HOM** | **4** | **Nebl** | **0.06** | **46.7%** | **72.5%** | **45.0%** | **1.23** | **35.8%** | **91.8%** | **74.1%** | **-0.09** | **53.4%** | **-0.38** | **0.66** | **0.63** | **30.3%** | **0.77** | **33.4%** | **-0.40** | **0.52** |
| 33 | Ostn<em1(IMPC)Wtsi> | HOM | 3 | Ostn | -1.15 | 23.7% | 17.8% | 14.1% | -1.28 | 26.4% | 70.4% | 26.6% | -0.86 | 40.1% | -0.96 | -0.08 | -0.26 | 46.8% | 0.26 | 14.0% | -1.38 | -0.74 |
| **34** | **Pitx1<em1(IMPC)Wtsi>** | **HET** | **7** | **Pitx1** | -4.91 | <0% | <0% | **6.6%** | -4.5 | **2.5%** | >100% | >100% | **0.73** | **73.5%** | **0.99** | **1.05** | **-0.42** | **20.4%** | **0.13** | **76.6%** | **-0.93** | **-0.18** |
| 35 | Plet1<em1(IMPC)Wtsi> | HOM | 3 | Plet1 | 0.43 | 27.2% | 39.6% | 6.9% | 0.14 | 26.0% | 73.0% | 28.6% | 0.28 | 95.7% | 0.63 | 1.18 | -0.22 | 12.9% | -0.22 | 77.1% | 1.18 | 1.04 |
| **36** | **Rab17<tm1a(KOMP)Wtsi>** | **HOM** | **3** | **Rab17** | -0.79 | 7.7% | 18.4% | **<0%** | -1.19 | 5.6% | 97.0% | 49.2% | 0.80 | 67.5% | 0.41 | 0.21 | -0.06 | 39.2% | 1.24 | 57.9% | 0.05 | 0.32 |
| 37 | Rbm33<tm1b(EUCOMM)Wtsi> | HET | 3 | Rbm33 | 1.26 | 70.4% | 59.7% | 44.8% | -0.03 | 36.3% | 94.1% | 55.9% | 0.94 | 31.8% | 1.07 | -0.32 | -0.04 | 64.8% | -0.34 | 43.9% | 1.43 | 1.39 |
| **38** | **Rnf114<em1(IMPC)Wtsi>** | **HOM** | **3** | **Rnf114** | **-0.22** | **74.9%** | **27.7%** | **54.4%** | **-0.21** | **71.2%** | **97.9%** | **25.3%** | **-0.35** | **78.4%** | **-0.12** | **0.97** | **0.42** | **22.4%** | **-1.36** | **74.1%** | **0.49** | **0.98** |
| **39** | **Scaf11<em1(IMPC)Wtsi>** | **HOM** | **5** | **Scaf11** | -0.44 | 17.6% | 18.4% | **4.1%** | -0.73 | **5.1%** | 88.5% | 44.8% | 0.68 | 41.4% | 0.20 | 0.01 | 0.48 | 44.9% | 0.81 | 47.6% | 0.34 | -0.19 |
| **40** | **Sh3bp4<tm1a(EUCOMM)Wtsi>** | **HOM** | **5** | **Sh3bp4** | **-1.70** | 36.7% | **5.5%** | 44.7% | **-1.74** | 35.8% | 97.5% | 71.3% | 0.29 | 38.8% | -0.58 | -0.23 | 1.53 | 64.4% | 0.87 | 49.6% | -1.05 | -0.01 |
| **41** | **Slamf9<tm1b(EUCOMM)Wtsi>** | **HOM** | **4** | **Slamf9** | 0.11 | 36.9% | 59.6% | 33.4% | 0.30 | 36.3% | **97.4%** | 76.4% | 0.99 | 63.2% | 0.67 | 0.81 | 0.35 | 22.2% | -0.07 | 51.6% | 0.30 | 0.43 |
| **42** | **Smg9<tm1b(EUCOMM)Wtsi>** | **HET** | **3** | **Smg9** | -1.05 | 54.0% | <0% | 14.5% | -1.99 | 4.9% | 40.7% | 21.2% | 0.24 | 40.6% | 0.35 | -0.50 | 0.01 | 69.7% | 0.32 | 57.3% | -0.26 | 0.09 |
| 43 | Stau2<tm1a(EUCOMM)Wtsi> | HOM | 4 | Stau2 | -0.83 | 70.9% | 18.0% | 53.9% | -0.97 | 36.3% | 47.3% | 57.9% | 0.66 | 18.4% | 0.14 | -1.02 | 1.00 | 79.3% | 0.82 | 17.9% | -0.07 | 0.38 |
| 44 | Timeless<tm1b(EUCOMM)Hmgu> | HET | 5 | Timeless | 0.02 | 22.7% | 48.7% | 14.4% | -0.57 | 26.7% | 91.2% | 41.6% | 0.22 | 43.4% | 0.00 | 0.43 | -0.06 | 39.8% | -0.31 | 75.3% | -0.30 | -0.90 |
| 45 | Tmem37<tm1a(EUCOMM)Wtsi> | HOM | 4 | Tmem37 | -0.60 | 54.0% | 10.3% | 45.0% | -0.97 | 60.8% | 73.6% | 68.7% | 0.10 | 62.6% | -0.04 | 0.37 | -0.10 | 38.8% | 0.28 | 57.3% | 0.72 | 0.07 |
| **46** | **Unk<tm1a(KOMP)Wtsi>** | **HOM** | **4** | **Unk** | **-1.82** | 28.2% | 7.4% | 14.8% | -1.50 | 2.2% | 82.3% | **>100%** | 0.34 | 25.6% | -0.33 | -0.98 | 1.19 | 84.4% | 0.75 | 6.4% | -1.20 | -0.77 |
| 47 | Upb1<tm1a(EUCOMM)Wtsi> | HOM | 3 | Upb1 | -0.73 | 80.8% | 38.9% | 67.3% | -0.30 | 80.5% | 89.3% | 57.5% | 0.22 | 25.9% | -0.46 | -0.91 | 1.77 | 80.2% | -0.41 | 32.8% | 0.24 | -0.98 |
| 48 | Vamp3<tm2b(EUCOMM)Wtsi> | HOM | 3 | Vamp3 | -0.42 | 61.7% | 26.7% | 67.3% | -0.21 | 70.9% | 56.0% | 36.7% | -1.45 | 15.3% | -1.51 | -0.55 | 0.11 | 55.2% | -1.86 | 33.9% | -1.76 | -0.51 |
| 49 | Wac<tm1b(EUCOMM)Wtsi> | HET | 3 | Wac | 1.99 | 87.0% | 73.2% | 77.8% | 0.77 | 71.2% | 97.0% | 52.0% | 0.16 | 71.9% | 0.24 | 0.53 | -0.06 | 37.7% | -0.87 | 69.8% | 1.36 | 1.51 |
| **50** | **Zfp341<tm1a(KOMP)Wtsi>** | **HOM** | **4** | **Zfp341** | **-0.21** | **36.8%** | **18.3%** | **24.5%** | **-0.97** | **36.3%** | >100% | **96.8%** | **0.97** | **76.7%** | **0.08** | **0.38** | **1.48** | **39.4%** | **1.27** | **56.4%** | **-0.31** | **-0.60** |

| | Ref. range | Wilcoxon | Mahalanobis |
|---|---|---|---|
| **Normal** | | | |
| **Ref. range** | | | |
| **Wilcoxon** | | | |
| **Mahalanobis** | | | |
| **Ref. range + Mahalanobis** | | | |
| **Wilcoxon + Mahalanobis** | | | |
| **Ref.range + Wilcoxon** | | | |
| **Ref. range + Mahalanobis + Wilcoxon** | | | |

**Statistical analysis of joint phenotyping pipeline data from 50 unselected mouse lines.**

P-values (Wilcoxon sum rank test)

| Significance | |
|---|---|
| n.s. | |
| * | p<0.00568 |
| ** | P<0.001 |
| *** | P<0.0001 |

| | | | | Tissue compartment: | Articular cartilage | | | | | | | | Subchondral bone | | | | | | | | | |
|---|---|---|---|---|---|---|---|---|---|---|---|---|---|---|---|---|---|---|---|---|---|---|
| | | | | Parameter: | ACV | | Median AC Th | | Maximum AC Th | | AC Damage Area | | SC BV/TV | | SC Tb.Th | | SC Tb.N | | SC BMD | | SC BMC | |
| | | | | Unit: | ($mm^3$) | | (mm) | | (mm) | | (%) | | (%) | | (mm) | | | | (mg.HA/$cm^3$) | | (median grey level) | |
| | | | | Technique: | ICE-µCT | | ICE-µCT | | ICE-µCT | | JSR | | ICE-µCT | | ICE-µCT | | ICE-µCT | | ICE-µCT | | scXRM | |
| | | | | Reference range: | mean ± 2SD | 2.5–97.5% | 2.5–97.5% | 2.5–97.5% | mean ± 2SD | 2.5–97.5% | 2.5–97.5% | 2.5–97.5% | mean ± 2SD | 2.5–97.5% | mean ± 2SD | mean ± 2SD | mean ± 2SD | 2.5-97.5% | mean ± 2SD | 2.5–97.5% | mean ± 2SD | mean ± 2SD |
| Line | Knockout Allele | Genotype | Samples (n) | Gene | LTP | MTP | LTP | MTP | LTP | MTP | LTP | MTP | LTP | MTP | LTP | MTP | LTP | MTP | LTP | MTP | LTP | MTP |
| 1 | 4932431P20Rik<em1(IMPC)Wtsi> | HOM | 6 | 4932431P20Rik | 0.02538 | 0.46863 | 0.01593 | 0.02143 | 0.95623 | 0.01311 | 0.01975 | 0.81091 | 0.54747 | 0.00693 | 0.17807 | 0.00612 | 0.57932 | 0.01257 | 0.93462 | 0.91290 | 0.03038 | 0.28529 |
| 2 | A430078G23Rik<tm1a(KOMP)Wtsi> | HOM | 3 | A430078G23Rik | 0.95308 | 0.04758 | 0.88244 | 0.04772 | 0.60868 | 0.12901 | 0.08263 | 0.88306 | 0.37221 | 0.41571 | 0.64488 | 0.32198 | 0.76089 | 0.35652 | 0.09549 | 0.04440 | 0.65126 | 0.54896 |
| 3 | Arhgap30<tm1a(EUCOMM)Wtsi> | HOM | 3 | Arhgap30 | 0.83683 | 0.06379 | 0.08282 | 0.58118 | 0.09622 | 0.17377 | 0.01240 | 0.08981 | 0.28514 | 0.92968 | 0.16083 | 0.61701 | 0.68012 | 0.49234 | 0.06106 | 0.11438 | 0.47318 | 0.70155 |
| **4** | **Arrdc5<tm1b(EUCOMM)Wtsi>** | **HOM** | **4** | **Arrdc5** | 0.68494 | 0.02797 | 0.46502 | **0.00225** | 0.45014 | 0.02905 | 0.01459 | 0.25389 | 0.39332 | 0.75450 | 0.36134 | 0.63601 | 0.83907 | 0.41219 | 0.12816 | 0.69743 | 0.70942 | 0.02952 |
| **5** | **Bhlhe40<tm1b(KOMP)Wtsi>** | **HOM** | **3** | **Bhlhe40** | 0.02119 | 0.02874 | **0.00469** | 0.01016 | **0.00347** | **0.00498** | 0.16085 | 0.12368 | 0.13354 | 0.91410 | 0.31720 | 0.42704 | 0.56247 | 0.38804 | 0.55627 | 0.17286 | 0.27971 | 0.88281 |
| 6 | Ccdc6<em1(IMPC)Wtsi> | HET | 4 | Ccdc6 | 0.76860 | 0.20581 | 0.79771 | 0.08477 | 0.73780 | 0.11019 | 0.12854 | 0.10997 | 0.48017 | 1.00000 | 0.76862 | 0.91410 | 0.37698 | 0.94526 | 0.23928 | 0.79119 | 0.96082 | 0.58214 |
| 7 | Cfap53<em1(IMPC)Wtsi> | HET | 6 | Cfap53 | 0.71200 | 0.55655 | 0.33622 | 0.04354 | 0.30336 | 0.19647 | 0.18706 | 0.84286 | 0.09946 | 0.62743 | 0.23425 | 0.38158 | 0.72196 | 0.50716 | 0.12910 | 0.13436 | 0.32717 | 0.86399 |
| **8** | **Chka<tm2a(KOMP)Wtsi>** | **HET** | **4** | **Chka** | 0.14136 | 0.15314 | 0.26228 | 0.22155 | 0.46046 | 0.36790 | **0.00270** | 0.04893 | 0.24692 | 0.52617 | 0.28310 | 0.89912 | 0.78006 | 0.78026 | 0.07315 | 0.29853 | 0.02650 | 0.30153 |
| 9 | ENSMUSG00000065619<(tm1)Brd> | HOM | 3 | Clust6N1 | 0.31244 | 0.54967 | 0.62913 | 0.70811 | 0.33962 | 0.59441 | 0.92968 | 0.36180 | 0.61700 | 0.56290 | 0.56288 | 0.63787 | 0.47968 | 0.50477 | 0.90632 | 0.63785 | 0.75318 | 0.19798 |
| **10** | **ClusterXN1<tm1(Brd)>** | **HEMI** | **3** | **ClusterXN1** | 0.13865 | 0.53022 | 0.20711 | 0.58129 | 0.41386 | 0.50904 | 0.00660 | 0.30319 | 0.58966 | 0.25126 | 0.57619 | 0.20239 | 0.22728 | 0.17275 | 0.01909 | **0.00519** | 0.51660 | 0.93735 |
| 11 | Col4a2<em1(IMPC)Wtsi> | HET | 3 | Col4a2 | 0.28069 | 0.38811 | 0.58101 | 0.15314 | 0.67927 | 0.47770 | 0.01173 | 0.42704 | 0.41571 | 0.14400 | 0.96871 | 0.29407 | 0.58246 | 0.23918 | 0.73881 | 0.90632 | 0.66540 | 0.35059 |
| **12** | **Cpgi81<tm1.1(NCC)WCS>** | **HOM** | **5** | **Cpgi81** | 0.22002 | 0.92805 | 0.04507 | 0.50606 | 0.24179 | 0.86789 | **0.00435** | 0.10255 | 0.95800 | 0.25593 | 0.23751 | 0.05897 | 0.16578 | 0.07823 | 0.20892 | 0.38690 | 0.02024 | 0.23656 |
| 13 | Dctn4<em1(IMPC)Wtsi> | HET | 6 | Dctn4 | 0.07661 | 0.40037 | 0.24851 | 0.26594 | 0.78371 | 0.16517 | 0.04668 | 0.52495 | 0.18030 | 0.30842 | 0.11908 | 0.32831 | 0.85876 | 0.62249 | 0.81089 | 0.69174 | 0.07946 | 0.90187 |
| 14 | Deptor<tm1b(EUCOMM)Wtsi> | HOM | 3 | Deptor | 0.43271 | 0.65188 | 0.02343 | 0.58799 | 0.05748 | 0.55429 | 0.61701 | 0.01240 | 0.29407 | 0.48629 | 0.27638 | 0.65902 | 0.44975 | 0.96088 | 0.17595 | 0.48628 | 0.39803 | 0.12292 |
| 15 | Dnah17<tm1e(KOMP)Wtsi> | HOM | 3 | Dnah17 | 0.37936 | 0.06661 | 0.28063 | 0.74697 | 0.61664 | 0.68320 | 0.22683 | 0.37486 | 0.44184 | 0.04167 | 0.59440 | 0.04083 | 0.81930 | 0.18164 | 0.20485 | 0.02798 | 0.37842 | 0.76041 |
| 16 | Washc2<tm2b(KOMP)Wtsi> | HET | 4 | Fam21 | 0.03682 | 0.52060 | 0.10649 | 1.00000 | 0.11078 | 0.05597 | 0.00816 | 0.29855 | 0.07183 | 0.12608 | 0.17094 | 0.31454 | 0.76712 | 0.17888 | 0.04891 | 0.69119 | 0.02125 | 0.79287 |
| **17** | **Fgfbp1<em1(IMPC)Wtsi>** | **HOM** | **4** | **Fgfbp1** | 0.26816 | 0.17894 | 0.77927 | 0.02002 | 0.95264 | 0.94583 | 0.17901 | 0.04893 | 0.36136 | 0.40276 | 0.25037 | 0.75450 | **0.00413** | 0.95955 | 0.10644 | 0.04338 | 0.03880 | 0.14292 |
| 18 | Gm6578<tm1a(KOMP)Wtsi> | HOM | 3 | Gm6578 | 0.13605 | 0.43269 | 0.47180 | 0.67903 | 0.41388 | 0.66449 | 0.38282 | 0.32199 | 0.39906 | 0.96090 | 0.13868 | 0.89856 | 0.35613 | 0.71666 | 1.00000 | 0.64488 | 0.04607 | 0.37122 |
| 19 | Gpbp1l1<tm1a(EUCOMM)Wtsi> | HOM | 3 | Gpbp1l1 | 0.04543 | 0.30312 | 0.02112 | 0.32462 | 0.23355 | 0.26535 | 0.68766 | 0.11897 | 0.65902 | 0.43854 | 0.73144 | 0.89856 | 0.13568 | 0.77607 | 0.15791 | 0.37747 | 0.95298 | 0.34548 |
| 20 | Grsf1<tm1b(EUCOMM)Wtsi> | HOM | 3 | Grsf1 | 0.89855 | 0.05970 | 0.90584 | 0.23716 | 0.55475 | 0.15869 | 0.15507 | 0.03023 | 0.01311 | 0.92189 | 0.01860 | 0.85220 | 0.90622 | 0.64478 | 0.65900 | 0.91410 | 0.48538 | 0.31626 |
| **21** | **Gsdme<tm1b(KOMP)Wtsi>** | **HOM** | **4** | **Gsdme** | 0.67257 | 0.87906 | 0.45472 | 0.78584 | 0.76006 | 0.93904 | **0.00355** | 0.59442 | 0.21409 | 0.84587 | 0.64810 | 0.65420 | 0.41665 | 0.77378 | 0.28688 | 0.27558 | 0.71568 | 0.87217 |
| 22 | Hao2<em1(IMPC)Wtsi> | HOM | 3 | Hao2 | 0.04983 | 0.78361 | 0.03329 | 0.64335 | 0.02817 | 0.98427 | 0.43854 | 0.16085 | 0.45022 | 0.41011 | 0.39904 | 0.36180 | 0.51706 | 0.45600 | 0.82919 | 0.80634 | 0.40915 | 0.97648 |
| **23** | **Herc1<em2(IMPC)Wtsi>** | **HOM** | **4** | **Herc1 (T1696A)** | 0.01878 | 0.27182 | **0.00321** | 0.02003 | 0.01324 | 0.02905 | 0.89912 | 0.01070 | 0.98651 | 0.05505 | 0.21408 | 0.05090 | 0.18974 | 0.14828 | 0.90581 | 0.01424 | 0.45612 | 0.66581 |
| 24 | Hmgxb3<tm1a(EUCOMM)Wtsi> | HET | 4 | Hmgxb3 | 0.37027 | 0.74805 | 0.23440 | 0.38634 | 0.39618 | 0.48047 | 1.00000 | 0.07183 | 0.17361 | 0.66645 | 0.09933 | 0.50434 | 0.89231 | 0.48817 | 0.33953 | 0.05721 | 0.32576 | 0.10214 |
| 25 | Hpf1<em1(IMPC)Wtsi> | HOM | 5 | Hpf1 | 0.11581 | 0.01387 | 0.29338 | 0.01376 | 1.00000 | 0.14430 | 0.05897 | 0.64628 | 0.30978 | 0.01331 | 0.63550 | 0.02968 | 0.19006 | 0.01415 | 0.09485 | 0.77495 | 0.25800 | 0.03542 |
| **26** | **Josd1<em1(IMPC)Wtsi>** | **HOM** | **6** | **Josd1** | 0.33850 | 0.41587 | **0.00344** | **0.00487** | **0.00362** | 0.02465 | **0.00563** | 0.91833 | 0.05389 | 0.34901 | 0.31825 | 0.41204 | 0.16681 | 0.36675 | 0.04372 | 0.30194 | 0.88022 | 0.69613 |
| 27 | Lpxn<tm1b(EUCOMM)Hmgu> | HOM | 3 | Lpxn | 0.85218 | 0.07269 | 0.72270 | 0.52834 | 0.43101 | 0.52807 | 0.73145 | 0.77613 | 0.49866 | 0.89856 | 0.82156 | 0.96090 | 0.16040 | 0.79871 | 0.27206 | 0.62391 | 0.30676 | 1.00000 |
| 28 | Lrrc8d<tm1a(EUCOMM)Wtsi> | HOM | 3 | Lrrc8d | 0.41006 | 0.55623 | 0.61522 | 0.11729 | 0.98429 | 0.58758 | 0.18234 | 0.02473 | 0.63088 | 0.25946 | 0.25125 | 0.45022 | 0.14627 | 0.41558 | 0.14942 | 0.28957 | 0.33555 | 0.97648 |
| 29 | Mbd1<em1(IMPC)Wtsi> | HOM | 4 | Mbd1 | 0.04165 | 0.16310 | 0.36333 | 0.58671 | 0.70262 | 0.87175 | 0.02012 | 0.85913 | 0.81293 | 0.42203 | 0.91922 | 0.72896 | 0.79310 | 0.84582 | 0.36133 | 0.53724 | 0.28196 | 0.87882 |
| **30** | **Mkrn2<em1(IMPC)Wtsi>** | **HOM** | **4** | **Mkrn2** | 0.05944 | 0.81947 | 0.03098 | 0.36812 | 0.00968 | 0.09761 | **0.00228** | 0.25389 | 0.15320 | 0.20189 | 0.66643 | 0.57122 | 0.95276 | 0.56538 | 0.63598 | 0.84587 | 0.39226 | 0.36480 |
| 31 | Mroh9<tm1a(EUCOMM)Wtsi> | HOM | 3 | Mroh9 | 0.39355 | 0.49237 | 0.79771 | 0.30086 | 0.26590 | 0.65026 | 0.70215 | 0.13869 | 0.73144 | 0.71674 | 0.25944 | 0.68765 | 0.24669 | 0.86756 | 0.41008 | 0.73143 | 0.29294 | 0.23057 |
| **32** | **Nebl<tm1b(EUCOMM)Wtsi>** | **HOM** | **4** | **Nebl** | 0.85911 | 0.79981 | 0.70234 | 0.67740 | 0.75361 | 0.40026 | **0.00489** | 0.09260 | 0.81950 | 0.64205 | 0.45192 | 0.81950 | 0.10071 | 0.54274 | 0.21406 | 0.27932 | 0.38758 | 0.26726 |
| 33 | Ostn<em1(IMPC)Wtsi> | HOM | 3 | Ostn | 0.03672 | 0.12124 | 0.10186 | 0.07300 | 0.10648 | 0.57411 | 0.25946 | 0.12854 | 0.12368 | 0.71674 | 0.07591 | 0.89856 | 0.58247 | 0.85985 | 0.73881 | 0.04985 | 0.12053 | 0.21209 |
| **34** | **Pitx1<em1(IMPC)Wtsi>** | **HET** | **7** | **Pitx1** | **0.00001** | **0.00008** | **0.00001** | **0.00307** | **0.00001** | **0.00319** | **0.00014** | **0.00001** | 0.15637 | 0.06402 | 0.04316 | 0.02617 | 0.49984 | 0.03424 | 0.73371 | 0.02334 | 0.09427 | 0.82027 |
| 35 | Plet1<em1(IMPC)Wtsi> | HOM | 3 | Plet1 | 0.36692 | 0.15216 | 0.74496 | 0.11056 | 0.65062 | 0.20356 | 0.25946 | 0.19217 | 0.65902 | 0.00765 | 0.23156 | 0.02736 | 0.64450 | 0.03018 | 0.70212 | 0.09550 | 0.03140 | 0.05291 |
| 36 | Rab17<tm1a(KOMP)Wtsi> | HOM | 3 | Rab17 | 0.16674 | 0.22023 | 0.08106 | 0.12934 | 0.03969 | 0.11482 | 0.13354 | 0.73145 | 0.22030 | 0.29407 | 0.42133 | 0.74625 | 0.93740 | 0.46794 | 0.15221 | 0.91410 | 0.81356 | 0.66547 |
| 37 | Rbm33<tm1b(EUCOMM)Wtsi> | HET | 3 | Rbm33 | 0.14940 | 0.42697 | 0.66453 | 0.73025 | 0.83625 | 0.70067 | 0.01240 | 0.85220 | 0.06966 | 0.63088 | 0.04872 | 0.68045 | 0.96085 | 0.58955 | 0.54318 | 0.44434 | 0.14585 | 0.02774 |
| 38 | Rnf114<em1(IMPC)Wtsi> | HOM | 3 | Rnf114 | 0.59641 | 0.16372 | 0.32430 | 0.52832 | 0.60183 | 0.48405 | 0.02876 | 0.13354 | 0.53673 | 0.10780 | 0.78364 | 0.31249 | 0.47967 | 0.20930 | 0.03946 | 0.25944 | 0.54885 | 0.08203 |
| **39** | **Scaf11<em1(IMPC)Wtsi>** | **HOM** | **5** | **Scaf11** | 0.24047 | 0.02260 | 0.03362 | **0.00053** | 0.07345 | **0.00179** | 0.03083 | 0.37063 | 0.17327 | 0.59322 | 0.73493 | 0.98799 | 0.42029 | 0.66794 | 0.11935 | 0.48410 | 0.37771 | 0.44642 |
| **40** | **Sh3bp4<tm1a(EUCOMM)Wtsi>** | **HOM** | **5** | **Sh3bp4** | **0.00311** | 0.39099 | **0.00081** | 0.54035 | **0.00133** | 0.30369 | **0.00282** | 0.23160 | 0.49357 | 0.53728 | 0.24050 | 0.37872 | 0.01022 | 0.37858 | 0.04144 | 0.42078 | 0.23647 | 0.94590 |
| **41** | **Slamf9<tm1b(EUCOMM)Wtsi>** | **HOM** | **4** | **Slamf9** | 0.86576 | 0.46209 | 0.62220 | 0.39576 | 0.62863 | 0.54062 | **0.00136** | 0.59442 | 0.06418 | 0.24009 | 0.17359 | 0.18176 | 0.91242 | 0.14829 | 0.82606 | 0.87908 | 0.47153 | 0.56470 |
| **42** | **Smg9<tm1b(EUCOMM)Wtsi>** | **HET** | **3** | **Smg9** | 0.04040 | 0.85218 | **0.00377** | 0.07952 | 0.00568 | 0.04537 | 0.61701 | 0.08616 | 0.63088 | 0.61701 | 0.60325 | 0.32680 | 0.99217 | 0.23529 | 0.43851 | 0.94527 | 0.61622 | 0.96081 |
| 43 | Stau2<tm1a(EUCOMM)Wtsi> | HOM | 4 | Stau2 | 0.05831 | 0.20785 | 0.03903 | 0.76635 | 0.04005 | 0.54062 | 0.58277 | 0.55979 | 0.14371 | 0.03250 | 0.83926 | 0.02985 | 0.06637 | 0.04696 | 0.06060 | 0.06791 | 0.93249 | 0.38761 |
| 44 | Timeless<tm1b(EUCOMM)Hmgu> | HET | 5 | Timeless | 0.99400 | 0.04295 | 0.89172 | 0.03126 | 0.15563 | 0.18035 | 0.00944 | 0.31700 | 0.65712 | 0.87446 | 0.95800 | 0.58285 | 0.81534 | 0.40777 | 0.66255 | 0.08354 | 0.40693 | 0.09877 |
| 45 | Tmem37<tm1a(EUCOMM)Wtsi> | HOM | 4 | Tmem37 | 0.13463 | 0.88574 | 0.02089 | 0.69612 | 0.06079 | 0.49103 | 0.32275 | 0.27560 | 0.91251 | 0.27560 | 0.95282 | 0.39332 | 0.91914 | 0.33942 | 0.50431 | 0.91921 | 0.09191 | 0.97297 |
| **46** | **Unk<tm1a(KOMP)Wtsi>** | **HOM** | **4** | **Unk** | **0.00215** | 0.13461 | 0.00983 | 0.08324 | 0.01966 | **0.00295** | 0.03250 | 0.93937 | 0.46732 | 0.13467 | 0.48827 | 0.05722 | 0.01954 | 0.03679 | 0.09759 | 0.01266 | 0.01405 | 0.06488 |
| 47 | Upb1<tm1a(EUCOMM)Wtsi> | HOM | 3 | Upb1 | 0.18880 | 0.08975 | 0.47786 | 0.20025 | 0.60881 | 0.10388 | 0.21301 | 0.80634 | 0.72408 | 0.16085 | 0.38280 | 0.07922 | 0.01413 | 0.09542 | 0.29857 | 0.21299 | 0.56868 | 0.05923 |
| 48 | Vamp3<tm2b(EUCOMM)Wtsi> | HOM | 3 | Vamp3 | 0.31244 | 0.44428 | 0.13927 | 0.20025 | 0.60183 | 0.20705 | 0.91410 | 0.42704 | 0.01206 | 0.32198 | 0.01502 | 0.43854 | 0.89846 | 0.85216 | 0.01461 | 0.34150 | 0.00846 | 0.49779 |
| **49** | **Wac<tm1b(EUCOMM)Wtsi>** | **HET** | **3** | **Wac** | 0.01672 | 0.09942 | 0.17067 | 0.34940 | 0.42524 | 0.80539 | 0.04873 | 0.33166 | 0.89081 | 0.14400 | 0.62392 | 0.27640 | 0.84436 | 0.36164 | 0.20236 | 0.27207 | 0.04826 | 0.03637 |
| **50** | **Zfp341<tm1a(KOMP)Wtsi>** | **HOM** | **4** | **Zfp341** | 0.58853 | 0.27182 | 0.12838 | 0.18249 | 0.10154 | 0.54631 | **0.00144** | 0.13467 | 0.08021 | 0.07183 | 0.87908 | 0.38402 | 0.01784 | 0.38851 | 0.02680 | 0.89243 | 0.81252 | 0.22918 |

Spearman correlation matrix for statistically significant correlation coefficients (R-values, P<0.05) between joint phenotyping parameters.

| R-values (*P<0.05) | LTP ACV (mm3) | MTP ACV (mm3) | LTP Median AC Th (mm) | MTP Median AC Th (mm) | LTP Maximum AC Th (mm) | MTP Maximum AC Th (mm) | LTP AC Damage Area (%) | MTP AC Damage Area (%) | LTP SC BV/TV (%) | MTP SC BV/TV (%) | LTP SC Tb.Th (mm) | MTP SC Tb.Th (mm) | LTP SC Tb.N | MTP SC Tb.N | LTP SC BMD (mg HA/$cm^3$) | MTP SC BMD (mg HA/$cm^3$) | LTP SC BMC | MTP SC BMC |
|---|---|---|---|---|---|---|---|---|---|---|---|---|---|---|---|---|---|---|
| LTP ACV (mm3) | | | | | | | | | | | | | | | | | | |
| MTP ACV (mm3) | 0.54 | | | | | | | | | | | | | | | | | |
| LTP Median AC Th (mm) | 0.81 | 0.42 | | | | | | | | | | | | | | | | |
| MTP Median AC Th (mm) | 0.29 | 0.74 | 0.34 | | | | | | | | | | | | | | | |
| LTP Maximum AC Th (mm) | 0.63 | 0.40 | 0.72 | 0.33 | | | | | | | | | | | | | | |
| MTP Maximum AC Th (mm) | 0.45 | 0.73 | 0.45 | 0.80 | 0.43 | | | | | | | | | | | | | |
| LTP AC Damage Area (%) | | 0.25 | | 0.25 | | 0.21 | | | | | | | | | | | | |
| MTP AC Damage Area (%) | | | | | | | | | | | | | | | | | | |
| LTP SC BV/TV (%) | 0.27 | | | | | | 0.22 | | | | | | | | | | | |
| MTP SC BV/TV (%) | 0.37 | | 0.24 | | | | 0.21 | | 0.56 | | | | | | | | | |
| LTP SC Tb.Th (mm) | 0.43 | 0.22 | 0.26 | | 0.20 | | 0.25 | | 0.87 | 0.65 | | | | | | | | |
| MTP SC Tb.Th (mm) | 0.46 | | 0.29 | | | | 0.26 | | 0.61 | 0.96 | 0.72 | | | | | | | |
| LTP SC Tb.N | -0.22 | | -0.26 | | -0.23 | | | | 0.32 | | | | | | | | | |
| MTP SC Tb.N | -0.47 | | -0.28 | | | | -0.26 | | -0.62 | -0.94 | -0.73 | -0.98 | | | | | | |
| LTP SC BMD (mg HA/$cm^3$) | | | | | | | | | 0.74 | 0.27 | 0.66 | 0.33 | 0.24 | -0.38 | | | | |
| MTP SC BMD (mg HA/$cm^3$) | | | | | | | | -0.23 | | 0.48 | | 0.39 | -0.20 | -0.31 | -0.35 | | | |
| LTP SC BMC | 0.40 | 0.26 | 0.21 | | | 0.22 | 0.32 | | 0.68 | 0.55 | 0.75 | 0.63 | | -0.64 | 0.36 | | | |
| MTP SC BMC | 0.48 | 0.42 | 0.31 | | 0.22 | 0.30 | 0.21 | 0.23 | 0.43 | 0.60 | 0.55 | 0.63 | | -0.64 | 0.28 | | 0.51 | |

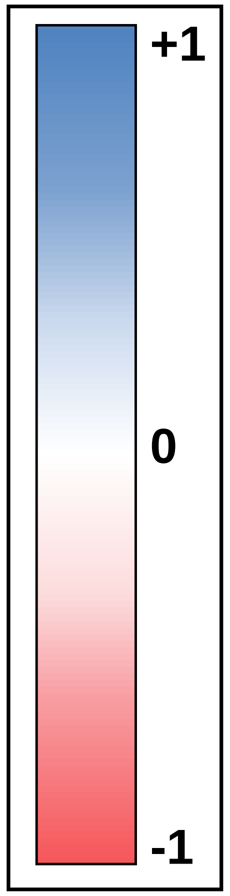

**Prioritization analysis of 25 mouse lines with abnormal joint phenotypes.**

| Gene nomenclature | | | | Mouse | | | Human | | |
|---|---|---|---|---|---|---|---|---|---|
| Line | Knockout Allele | Genotype | (n) | Mouse Gene | ENSEMBL ID | Gene Name | Human Gene | ENSEMBL ID | OVERALL SCORE /21 |
| 1 | Arhgap30<tm1a(EUCOMM)Wtsi> | HOM | 3 | **Arhgap30** | ENSMUSG00000048865 | Rho GTPase activating protein 30 | **ARHGAP30** | ENSG00000186517 | 4 |
| 2 | Arrdc5<tm1b(EUCOMM)Wtsi> | HOM | 4 | **Arrdc5** | ENSMUSG00000073380 | Arrestin domain containing 5 | **ARRDC5** | ENSG00000205784 | 3 |
| 3 | Bhlhe40<tm1b(KOMP)Wtsi> | HOM | 3 | **Bhlhe40** | ENSMUSG00000030103 | Basic helix-loop-helix family, member e4 | **BHLHE40** | ENSG00000134107 | 10 |
| 4 | Ccdc6<em1(IMPC)Wtsi> | HET | 3 | **Ccdc6** | ENSMUSG00000048701 | Coiled-coil domain containing 6 | **CCDC6** | ENSG00000108091 | 6 |
| 5 | Chka<tm2a(KOMP)Wtsi> | HET | 4 | **Chka** | ENSMUSG00000024843 | Choline kinase alpha | **CHKA** | ENSG00000110721 | 6 |
| 6 | ClusterXN1<tm1(Brd)> | HEMI | 3 | **ClusterXN1** | ENSMUSG00000065613 El | Mir92-2, Mir20b, Mir363, Mir19b-2, Mir1 | - | - | 6 |
| 7 | Col4a2<em1(IMPC)Wtsi> | HET | 3 | **Col4a2** | ENSMUSG00000031503 | Collagen, type IV, alpha 2 | **COL4A2** | ENSG00000134871 | 7 |
| 8 | Cpgi81<tm1.1(NCC)WCS> | HOM | 5 | **Cpgi81** | - | CpG island 81 | - | - | 4 |
| 9 | Fgfbp1<em1(IMPC)Wtsi> | HOM | 4 | **Fgfbp1** | ENSMUSG00000048373 | Fibroblast growth factor binding protein | **FGFBP1** | ENSG00000137440 | 5 |
| 10 | Grsf1<tm1b(EUCOMM)Wtsi> | HOM | 3 | **Grsf1** | ENSMUSG00000044221 | G-rich RNA sequence binding factor 1 | **GRSF1** | ENSG00000132463 | 4 |
| 11 | Gsdme<tm1b(KOMP)Wtsi> | HOM | 4 | **Gsdme** | ENSMUSG00000029821 | Gasdermin E | **GSDME** | ENSG00000105928 | 5 |
| 12 | Herc1<em2(IMPC)Wtsi> | HOM | 4 | **Herc1** | ENSMUSG00000038664 | HECT and RLD domain containing E3 u | **HERC1** | ENSG00000103657 | 7 |
| 13 | Hpf1<em1(IMPC)Wtsi> | HOM | 5 | **Hpf1** | ENSMUSG00000038005 | histone PARylation factor 1 | **HPF1** | ENSG00000056050 | 4 |
| 14 | Josd1<em1(IMPC)Wtsi> | HOM | 6 | **Josd1** | ENSMUSG00000022426 | Josephin domain containing 1 | **JOSD1** | ENSG00000100221 | 5 |
| 15 | Mkrn2<em1(IMPC)Wtsi> | HOM | 4 | **Mkrn2** | ENSMUSG00000000439 | Makorin, ring finger protein, 2 | **MKRN2** | ENSG00000075975 | 4 |
| 16 | Nebl<tm1b(EUCOMM)Wtsi> | HOM | 4 | **Nebl** | ENSMUSG00000053702 | Nebulette | **NEBL** | ENSG00000078114 | 5 |
| 17 | Pitx1<em1(IMPC)Wtsi> | HET | 7 | **Pitx1** | ENSMUSG00000021506 | Paired-like homeodomain transcription f | **PITX1** | ENSG00000069011 | 16 |
| 18 | Rab17<tm1a(KOMP)Wtsi> | HOM | 3 | **Rab17** | ENSMUSG00000026304 | RAB17, member RAS oncogene family | **RAB17** | ENSG00000124839 | 3 |
| 19 | Rnf114<em1(IMPC)Wtsi> | HOM | 3 | **Rnf114** | ENSMUSG00000006418 | Ring finger protein 114 | **RNF114** | ENSG00000124226 | 7 |
| 20 | Scaf11<em1(IMPC)Wtsi> | HOM | 5 | **Scaf11** | ENSMUSG00000033228 | SR-related CTD-associated factor 11 | **SCAF11** | ENSG00000139218 | 6 |
| 21 | Sh3bp4<tm1a(EUCOMM)Wtsi> | HOM | 5 | **Sh3bp4** | ENSMUSG00000036206 | SH3-domain binding protein 4 | **SH3BP4** | ENSG00000130147 | 8 |
| 22 | Slamf9<tm1b(EUCOMM)Wtsi> | HOM | 4 | **Slamf9** | ENSMUSG00000026548 | SLAM family member 9 | **SLAMF9** | ENSG00000162723 | 4 |
| 23 | Smg9<tm1b(EUCOMM)Wtsi> | HET | 3 | **Smg9** | ENSMUSG00000002210 | Smg-9 homolog, nonsense mediated m | **SMG9** | ENSG00000105771 | 7 |
| 24 | Unk<tm1a(KOMP)Wtsi> | HOM | 4 | **Unk** | ENSMUSG00000020770 | Unkempt family zinc finger | **UNK** | ENSG00000132478 | 8 |
| 25 | Zfp341<tm1a(KOMP)Wtsi> | HOM | 4 | **Zfp341** | ENSMUSG00000059842 | Zinc finger protein 341 | **ZNF341** | ENSG00000131061 | 6 |

| | OBCD Joint phenotypes | | | | | | |
|---|---|---|---|---|---|---|---|
| | Statistical analysis | | | Articular cartilage | | Subchondral bone | OBCD Joint |
| Line | Ref. Range | Wilcoxon test | Mahalanobis | Morphology | Damage | Morphology | Score /6 |
| 1 | 1 | 0 | 1 | 0 | 1 | 0 | 3 |
| 2 | 1 | 1 | 0 | 1 | 0 | 0 | 3 |
| 3 | 1 | 1 | 0 | 1 | 0 | 0 | 3 |
| 4 | 1 | 0 | 0 | 0 | 1 | 0 | 2 |
| 5 | 0 | 1 | 0 | 0 | 1 | 0 | 2 |
| 6 | 1 | 1 | 1 | 0 | 1 | 1 | 5 |
| 7 | 1 | 0 | 0 | 0 | 1 | 0 | 2 |
| 8 | 0 | 1 | 1 | 0 | 1 | 0 | 3 |
| 9 | 1 | 1 | 1 | 0 | 0 | 1 | 4 |
| 10 | 0 | 0 | 1 | 0 | 0 | 0 | 1 |
| 11 | 1 | 1 | 0 | 0 | 1 | 0 | 3 |
| 12 | 0 | 1 | 0 | 1 | 0 | 0 | 2 |
| 13 | 0 | 0 | 1 | 0 | 0 | 0 | 1 |
| 14 | 1 | 1 | 0 | 1 | 1 | 0 | 4 |
| 15 | 1 | 1 | 0 | 0 | 1 | 0 | 3 |
| 16 | 0 | 1 | 1 | 0 | 1 | 0 | 3 |
| 17 | 1 | 1 | 1 | 1 | 1 | 0 | 5 |
| 18 | 1 | 0 | 0 | 1 | 0 | 0 | 2 |
| 19 | 1 | 0 | 1 | 0 | 1 | 0 | 3 |
| 20 | 0 | 1 | 0 | 1 | 0 | 0 | 2 |
| 21 | 1 | 1 | 0 | 1 | 1 | 0 | 4 |
| 22 | 0 | 1 | 0 | 0 | 1 | 0 | 2 |
| 23 | 1 | 1 | 0 | 1 | 0 | 0 | 3 |
| 24 | 1 | 1 | 0 | 1 | 1 | 0 | 4 |
| 25 | 1 | 1 | 1 | 0 | 1 | 0 | 4 |

| | Other KO mouse phenotypes | | | | | |
|---|---|---|---|---|---|---|
| Line | OBCD Bone Phenotype | IMPRESS Skeletal | DMDD Skeletal | PubMed skeletal | MGI Skeletal | Phenotype Score /4 |
| 1 | pending | 0 | | 0 | 0 | 0 |
| 2 | pending | 0 | | 0 | 0 | 0 |
| 3 | 1 | 1 | | 0 | 1 | 3 |
| 4 | 1 | 0 | | 0 | 0 | 1 |
| 5 | 1 | 0 | 0 | 1 | 0 | 2 |
| 6 | 1 | 0 | | 0 | | 1 |
| 7 | 0 | 0 | 1 | 0 | 0 | 1 |
| 8 | 1 | 0 | | 0 | | 1 |
| 9 | pending | 0 | | 0 | 0 | 0 |
| 10 | 1 | 0 | | 0 | 0 | 1 |
| 11 | pending | 0 | | 0 | 0 | 0 |
| 12 | 1 | 1 | | 0 | 0 | 2 |
| 13 | pending | 1 | | 0 | 1 | 2 |
| 14 | 0 | 0 | | 0 | 0 | 0 |
| 15 | pending | 0 | | 0 | 0 | 0 |
| 16 | 0 | 0 | | 0 | 0 | 0 |
| 17 | 1 | 1 | 0 | 1 | 1 | 4 |
| 18 | 0 | 0 | | 0 | 0 | 0 |
| 19 | 1 | 0 | | 1 | 0 | 2 |
| 20 | 1 | 1 | | 0 | 1 | 3 |
| 21 | 1 | 0 | | 0 | 0 | 1 |
| 22 | 1 | 0 | | 0 | 0 | 1 |
| 23 | 1 | 0 | 1 | 0 | 1 | 3 |
| 24 | pending | 1 | 0 | 0 | 1 | 2 |
| 25 | pending | 0 | | 0 | 0 | 0 |

| | Gene expression | | | | | | | | | | | | | | |
|---|---|---|---|---|---|---|---|---|---|---|---|---|---|---|---|
| | | | | Chondrocytes | | | Osteoblasts | | Osteocytes | | Osteoclasts | | | | |
| Line | MGI Skeletal | BioGPS Bone | Score | Cells 2019 | SkeletalVis | Score | BioGPS | SkeletalVis | RNAseq | Score | BioGPS | RNAseq | SkeletalVis | Score | Expression Score /4 |
| 1 | | 0.5 | 0.5 | 0 | 0 | 0.0 | 0 | 0.3 | 0 | 0.3 | 0.3 | 0 | 0 | 0.3 | 1.2 |
| 2 | | 0 | 0 | 0 | 0 | 0 | 0 | 0 | 0 | 0 | 0 | 0 | 0 | 0 | 0 |
| 3 | 0.5 | 0 | 0.5 | 0.5 | 0.5 | 1 | 0.3 | 0.3 | 0.3 | 0.3 | 0.3 | 0.3 | 0.3 | 1 | 2.8 |
| 4 | | 0 | 0 | 0.5 | | 0.5 | 0 | | 0 | 0 | 0 | 0 | | 0 | 0.5 |
| 5 | 0.5 | 0 | 0.5 | 0 | 0.5 | 0.5 | 0 | 0 | 0 | 0 | 0 | 0.3 | 0 | 0.3 | 1.3 |
| 6 | | | 0 | | | 0 | | | | 0 | | | | 0 | 0 |
| 7 | 0 | 0 | 0 | 0.5 | 0.5 | 1.0 | 0.3 | 0.3 | 0.3 | 1 | 0 | 0 | 0 | 0 | 2 |
| 8 | | | 0 | | | 0 | | | | 0 | | | | 0 | 0 |
| 9 | 0 | 0 | 0 | 0 | 0 | 0.0 | 0 | 0.3 | 0 | 0.3 | 0 | 0 | 0 | 0 | 0.3 |
| 10 | 0 | 0 | 0 | 0.5 | 0.5 | 1 | 0 | 0 | 0 | 0 | 0 | 0.3 | 0 | 0.3 | 1.3 |
| 11 | 0 | 0.5 | 0.5 | 0 | 0 | 0.0 | 0.3 | 0 | 0 | 0.3 | 0.3 | 0.3 | 0 | 0.7 | 1.5 |
| 12 | 0.5 | 0 | 0.5 | 0.5 | 0.5 | 1 | 0 | 0 | 0.3 | 0.3 | 0 | 0 | 0 | 0 | 1.8 |
| 13 | | 0.5 | 0.5 | | 0.5 | 1 | 0 | 0 | 0 | 0.0 | 0.3 | 0 | 0 | 0.3 | 1.3 |
| 14 | | 0 | 0 | 0.5 | 0 | 0.5 | 0 | 0 | 0 | 0 | 0 | 0.3 | 0 | 0.3 | 1 |
| 15 | 0 | 0 | 0 | 0.5 | 0.5 | 1 | 0 | 0 | 0 | 0 | 0 | 0.3 | 0 | 0.3 | 1.3 |
| 16 | 0 | 0 | 0 | | | 0 | 0 | | 0.3 | 0.3 | 0 | | | 0 | 0.3 |
| 17 | 0.5 | 0 | 0.5 | 0.5 | 0.5 | 1 | 0 | 0.3 | 0.3 | 0.7 | 0 | 0 | 0 | 0 | 2.2 |
| 18 | 0 | 0 | 0 | 0 | 0.5 | 0.5 | 0 | 0 | 0 | 0 | 0 | 0 | 0 | 0 | 0.5 |
| 19 | 0 | 0 | 0 | 0.5 | 0.5 | 1 | 0 | 0.3 | 0 | 0.3 | 0 | 0 | 0 | 0 | 1.3 |
| 20 | 0 | 0 | 0 | 0.5 | 0.5 | 1 | 0 | 0 | 0 | 0 | 0 | 0 | 0 | 0 | 1 |
| 21 | 0 | 0 | 0 | 0.5 | 0.5 | 1 | 0.3 | 0.3 | 0.3 | 1 | 0 | 0 | 0 | 0 | 2 |
| 22 | 0 | 0 | 0 | 0 | 0 | 0 | 0.3 | 0.3 | 0 | 0.7 | 0.3 | 0 | 0 | 0.3 | 1 |
| 23 | | 0 | 0 | 0.5 | 0.5 | 1 | 0 | 0 | 0 | 0.0 | 0 | 0.3 | 0 | 0.3 | 1.3 |
| 24 | 0 | 0 | 0 | 0.5 | 0 | 0.5 | 0 | 0.3 | 0 | 0.3 | 0 | 0.3 | 0 | 0.3 | 1.2 |
| 25 | 0 | 0 | 0 | 0 | 0.5 | 0.5 | 0 | 0.3 | 0 | 0.3 | 0 | 0 | 0 | 0 | 1 |

| | Human disease | | | | | Literature search (Date 01/07/2019) | | |
|---|---|---|---|---|---|---|---|---|
| | Monogenic disease | | Polygenic disease | | | | | |
| | MGI | OMIM | GWAS Catalogue | | Disease | | | Literature |
| Line | Human-Mouse | Disease | Arthritis | Bone | Score /4 | PubMed | Google Scholar | Score /2 |
| 1 | 0 | 0 | 0 | 0 | 0 | 0 | 0 | 0 |
| 2 | 0 | 0 | 0 | 0 | 0 | 0 | 0 | 0 |
| 3 | 0 | 0 | 0 | 0 | 0 | 1 | 0.5 | 1.5 |
| 4 | 0 | 1 | 0 | 1 | 2 | 0 | 0.5 | 0.5 |
| 5 | 0 | 0 | | | 0 | 0.5 | 0.5 | 1 |
| 6 | | | | | 0 | 0 | 0 | 0 |
| 7 | 0 | 0 | 0 | 0 | 0 | 0.5 | 1 | 1.5 |
| 8 | | | | | 0 | 0 | 0 | 0 |
| 9 | 0 | 0 | | | 0 | 0.5 | 0.5 | 1 |
| 10 | 0 | 0 | | | 0 | 0 | 0.5 | 0.5 |
| 11 | 0 | 0 | 0 | 0 | 0 | 0 | 0 | 0 |
| 12 | 0 | 0 | 0 | 1 | 1 | 0 | 0.5 | 0.5 |
| 13 | 0 | 0 | | | 0 | 0 | 0 | 0 |
| 14 | 0 | 0 | 0 | 0 | 0 | 0 | 0 | 0 |
| 15 | 0 | 0 | 0 | 0 | 0 | 0 | 0 | 0 |
| 16 | 0 | 1 | 0 | 0 | 1 | 0.5 | 0.5 | 1 |
| 17 | 1 | 1 | 0 | 1 | 3 | 0.5 | 1 | 1.5 |
| 18 | 0 | 0 | 0 | 0 | 0 | 0 | 0.5 | 0.5 |
| 19 | 0 | 0 | 0 | 0 | 0 | 0.5 | 0.5 | 1 |
| 20 | 0 | 0 | 0 | 0 | 0 | 0 | 0 | 0 |
| 21 | 0 | 0 | 0 | 1 | 1 | 0 | 0 | 0 |
| 22 | 0 | 0 | 0 | 0 | 0 | 0 | 0 | 0 |
| 23 | 0 | 0 | 0 | 0 | 0 | 0 | 0 | 0 |
| 24 | 0 | 0 | 0 | 0 | 0 | 0 | 0.5 | 0.5 |
| 25 | 0 | 0 | 0 | 1 | 1 | 0 | 0 | 0 |

**Animal species, strain, source, sex and unique identifier or Research Resource Identifier (RRID).**

| Mouse strain | Source | Unique identifier | Sex | Age |
|---|---|---|---|---|
| Mouse: Wild-type C57BL/6 | Charles River Labs | Strain: 027 | Male | 22 weeks |
| Mouse: Wild-type 129/SV/C57BL/6J | (Gauthier et al., 1999) | N/A | Male | 16 & 52 weeks |
| Mouse: *Dio2* $^{\textit{Thr92Thr}}$ | (Jo et al., 2019) | N/A | Male | 16 weeks |
| Mouse: *Dio2* $^{\textit{Ala92Ala}}$ | (Jo et al., 2019) | N/A | Male | 16 weeks |
| Mouse: Wild-type C57BL/6N Taconic;C57BL/6N from strains detailed below | IMPC | See following rows | Male | 16 weeks |
| Mouse: 4932431P20Rik-/-: 4932431P20Rik$^{\text{em1(IMPC)Wtsi/em1(IMPC)Wtsi}}$ | IMPC | RRID:MGI:6261757 | Male | 16 weeks |
| Mouse: A430078G23Rik-/-: A430078G23Rik$^{\text{tm1a(KOMP)Wtsi/tm1a(KOMP)Wtsi}}$ | IMPC | RRID:IMSR_KOMP:CSD77453-1a-Wtsi | Male | 16 weeks |
| Mouse: Arhgap30-/-: Arhgap30$^{\text{tm1a(EUCOMM)Wtsi/tm1a(EUCOMM)Wtsi}}$ | IMPC | RRID:MGI:6261895 | Male | 16 weeks |
| Mouse: Arrdc5-/-: Arrdc5$^{\text{tm1b(EUCOMM)Wtsi/tm1b(EUCOMM)Wtsi}}$ | IMPC | RRID:MGI:6261913 | Male | 16 weeks |
| Mouse: Bhlhe40-/-: Bhlhe40$^{\text{tm1b(KOMP)Wtsi/tm1b(KOMP)Wtsi}}$ | IMPC | RRID:MGI:6261982 | Male | 16 weeks |
| Mouse: Ccdc6+/-: Ccdc6$^{\text{em1(IMPC)Wtsi/em1(IMPC)Wtsi}}$ | IMPC | RRID:MGI:6262052 | Male | 16 weeks |
| Mouse: Cfap53+/-: Cfap53$^{\text{em1(IMPC)Wtsi/em1(IMPC)Wtsi}}$ | IMPC | RRID:MGI:6262116 | Male | 16 weeks |
| Mouse: Chka+/-: Chka$^{\text{tm2a(KOMP)Wtsi/WT}}$ | IMPC | RRID:IMSR_KOMP:CSD44635-1a-Wts | Male | 16 weeks |
| Mouse: Clust6N1(Mir183)-/-: Clust6N1$^{\text{(tm1)Brd/(tm1)Brd}}$ | IMPC | N/A | Male | 16 weeks |
| Mouse: ClusterXN1 (Mir92-2, Mir20b, Mir363, Mir19b-2, Mir18b, Mir106a)+/-: ClusterXN1$^{\text{tm1(Brd)/WT}}$ | IMPC | N/A | Male | 16 weeks |
| Mouse: Col4a2+/-: Col4a2$^{\text{em1(IMPC)Wtsi/WT}}$ | IMPC | RRID:MGI:6161265 | Male | 16 weeks |
| Mouse: Cpgi81-/-: Cpgi81$^{\text{tm1.1(NCC)WCS/tm1.1(NCC)WCS}}$ | IMPC | RRID:IMSR_EM:1226 | Male | 16 weeks |
| Mouse: Dctn4+/-: Dctn4$^{\text{em1(IMPC)Wtsi/WT}}$ | IMPC | RRID:MGI:6262256 | Male | 16 weeks |
| Mouse: Deptor-/-: Deptor$^{\text{tm1b(EUCOMM)Wtsi/tm1b(EUCOMM)Wtsi}}$ | IMPC | RRID:IMSR_EM:11080 | Male | 16 weeks |
| Mouse: Dnah17-/-: Dnah17$^{\text{tm1e(KOMP)Wtsi/tm1e(KOMP)Wtsi}}$ | IMPC | RRID:MGI:6262291 | Male | 16 weeks |
| Mouse: Washc2+/-: Fam21$^{\text{tm2b(KOMP)Wtsi/WT}}$ | IMPC | RRID:MGI:6263870 | Male | 16 weeks |
| Mouse: Fgfbp1-/-: Fgfbp1$^{\text{em1(IMPC)Wtsi/em1(IMPC)Wtsi}}$ | IMPC | RRID:IMSR_EM:11893 | Male | 16 weeks |
| Mouse: Gm6578-/-: Gm6578$^{\text{tm1a(KOMP)Wtsi/tm1a(KOMP)Wtsi}}$ | IMPC | RRID:MGI:6262538 | Male | 16 weeks |
| Mouse: Gpbp1l1-/-: Gpbp1l1$^{\text{tm1a(EUCOMM)Wtsi/tm1a(EUCOMM)Wtsi}}$ | IMPC | RRID:IMSR_EM:09065 | Male | 16 weeks |
| Mouse: Grsf1-/-: Grsf1$^{\text{tm1b(EUCOMM)Wtsi/tm1b(EUCOMM)Wtsi}}$ | IMPC | RRID:MGI:6262583 | Male | 16 weeks |
| Mouse: Gsdme-/-: Gsdme$^{\text{tm1b(KOMP)Wtsi/tm1b(KOMP)Wtsi}}$ | IMPC | RRID:IMSR_EM:09924 | Male | 16 weeks |
| Mouse: Hao2-/-: Hao2$^{\text{em1(IMPC)Wtsi/em1(IMPC)Wtsi}}$ | IMPC | RRID:IMSR_KOMP:CSD31359-1a-Wtsi | Male | 16 weeks |
| Mouse: Herc1-/-: T1696A$^{\text{em2(IMPC)Wtsi/em2(IMPC)Wtsi}}$ | IMPC | RRID:MGI:6262614 | Male | 16 weeks |
| Mouse: Hmgxb3+/-: Hmgxb3$^{\text{tm1a(EUCOMM)Wtsi/WT}}$ | IMPC | RRID:MGI:6102815 | Male | 16 weeks |
| Mouse: Hpf1-/-: 2700029M09Rik$^{\text{em1(IMPC)Wtsi/em1(IMPC)Wtsi}}$ | IMPC | RRID:MGI:6262645 | Male | 16 weeks |
| Mouse: Josd1-/-: Josd1$^{\text{em1(IMPC)Wtsi/em1(IMPC)Wtsi}}$ | IMPC | RRID:MGI:6262720 | Male | 16 weeks |
| Mouse: Lpxn-/-: Lpxn$^{\text{tm1b(EUCOMM)Hmgu/tm1b(EUCOMM)Hmgu}}$ | IMPC | RRID:IMSR_EM:09745 | Male | 16 weeks |
| Mouse: Lrrc8d-/-: Lrrc8d$^{\text{tm1a(EUCOMM)Wtsi/tm1a(EUCOMM)Wtsi}}$ | IMPC | RRID:MGI:6262826 | Male | 16 weeks |
| Mouse: Mbd1-/-: Mbd1$^{\text{em1(IMPC)Wtsi/em1(IMPC)Wtsi}}$ | IMPC | RRID:MGI:6262848 | Male | 16 weeks |
| Mouse: Mkrn2-/-: Mkrn2$^{\text{em1(IMPC)Wtsi/em1(IMPC)Wtsi}}$ | IMPC | RRID:MGI:6262889 | Male | 16 weeks |
| Mouse: Mroh9-/-: Mroh9$^{\text{tm1a(EUCOMM)Wtsi/tm1a(EUCOMM)Wtsi}}$ | IMPC | RRID:MGI:6262914 | Male | 16 weeks |
| Mouse: Nebl-/-: Nebl$^{\text{tm1b(EUCOMM)Wtsi/tm1b(EUCOMM)Wtsi}}$ | IMPC | RRID:IMSR_EM:09926 | Male | 16 weeks |
| Mouse: Ostn-/-: Ostn$^{\text{em1(IMPC)Wtsi/em1(IMPC)Wtsi}}$ | IMPC | RRID:IMSR_EM:10628 | Male | 16 weeks |
| Mouse: Pitx1+/-: Pitx1$^{\text{em1(IMPC)Wtsi/WT}}$ | IMPC | RRID:MGI:6263167 | Male | 16 weeks |
| Mouse: Plet1-/-: Plet1$^{\text{em1(IMPC)Wtsi/em1(IMPC)Wtsi}}$ | IMPC | RRID:MGI:6263184 | Male | 16 weeks |
| Mouse: Rab17-/-: Rab17$^{\text{tm1a(KOMP)Wtsi/tm1a(KOMP)Wtsi}}$ | IMPC | RRID:IMSR_KOMP:CSD79710-1a-Wtsi | Male | 16 weeks |
| Mouse: Rbm33+/-: Rbm33$^{\text{tm1b(EUCOMM)Wtsi/WT}}$ | IMPC | RRID:MGI:6263339 | Male | 16 weeks |
| Mouse: Rnf114-/-: Rnf114$^{\text{em1(IMPC)Wtsi/em1(IMPC)Wtsi}}$ | IMPC | RRID:IMSR_EM:10264 | Male | 16 weeks |
| Mouse: Scaf11-/-: Scaf11$^{\text{em1(IMPC)Wtsi/em1(IMPC)Wtsi}}$ | IMPC | RRID:MGI:6263429 | Male | 16 weeks |
| Mouse: Sh3bp4-/-: Sh3bp4$^{\text{tm1a(EUCOMM)Wtsi/tm1a(EUCOMM)Wtsi}}$ | IMPC | RRID:MGI:6263464 | Male | 16 weeks |
| Mouse: Slamf9-/-: Slamf9$^{\text{tm1b(EUCOMM)Wtsi/tm1b(EUCOMM)Wtsi}}$ | IMPC | RRID:IMSR_EM:0961 | Male | 16 weeks |
| Mouse: Smg9+/-: Smg9$^{\text{tm1b(EUCOMM)Wtsi/WT}}$ | IMPC | RRID:MGI:5883885 | Male | 16 weeks |
| Mouse: Stau2-/-: Stau2$^{\text{tm1a(EUCOMM)Wtsi/tm1a(EUCOMM)Wtsi}}$ | IMPC | RRID:MGI:6263584 | Male | 16 weeks |
| Mouse: Timeless+/-: Timeless$^{\text{tm1b(EUCOMM)Hmgu/WT}}$ | IMPC | RRID:IMSR_EM:09834 | Male | 16 weeks |
| Mouse: Tmem37-/-: Tmem37$^{\text{tm1a(EUCOMM)Wtsi/tm1a(EUCOMM)Wtsi}}$ | IMPC | RRID:IMSR_EM:11749 | Male | 16 weeks |
| Mouse: Unk-/-: Unk$^{\text{tm1a(KOMP)Wtsi/tm1a(KOMP)Wtsi}}$ | IMPC | RRID:MGI:5883888 | Male | 16 weeks |
| Mouse: Upb1-/-: Upb1$^{\text{tm1a(EUCOMM)Wtsi/tm1a(EUCOMM)Wtsi}}$ | IMPC | RRID:MGI:6263825 | Male | 16 weeks |
| Mouse: Vamp3-/-: Vamp3$^{\text{tm2b(EUCOMM)Wtsi/tm2b(EUCOMM)Wtsi}}$ | IMPC | RRID:IMSR_EM:09918 | Male | 16 weeks |
| Mouse: Wac+/-: Wac$^{\text{tm1b(EUCOMM)Wtsi/WT}}$ | IMPC | RRID:MGI:6263865 | Male | 16 weeks |
| Mouse: Zfp341-/-: Zfp341$^{\text{tm1a(KOMP)Wtsi/tm1a(KOMP)Wtsi}}$ | IMPC | RRID:MGI:6263912 | Male | 16 weeks |
| Mouse: Clic3+/-: Clic3$^{\text{em1(IMPC)Wtsi/WT}}$ | IMPC | RRID:IMSR_KOMP:CSD31844-1a-Wtsi | Male | 16 weeks |
| Mouse: Cpt1a-/-: Cpt1a$^{\text{tm1.2Wtsi/tm1.2Wtsi}}$ | IMPC | RRID:IMSR_EM:11842 | Male | 16 weeks |
| Mouse: Crip1-/-: Crip1$^{\text{em1(IMPC)Wtsi/em1(IMPC)Wtsi}}$ | IMPC | RRID:IMSR_KOMP:CSD89475-1e-Wtsi | Male | 16 weeks |
| Mouse: Htra3-/-: Htra3$^{\text{EM1/EM1}}$ | IMPC | RRID:IMSR_EM:11594 | Male | 16 weeks |
| Mouse: Matn4-/-: Matn4$^{\text{EM1/ EM1}}$ | IMPC | RRID:IMSR_EM:10571 | Male | 16 weeks |
| Mouse: Pdlim1-/-: Pdlim1$^{\text{em1(IMPC)Wtsi/ em1(IMPC)Wtsi}}$ | IMPC | RRID:MGI:6263129 | Male | 16 weeks |
| Mouse: Sqrdl+/-: Sqrdl$^{\text{em1(IMPC)Wtsi/WT}}$ | IMPC | RRID:MGI:6260869 | Male | 16 weeks |

Actual P-values for validation of new methods with surgical provocation of osteoarthritis by DMM surgery, Application 2 and Application 3.

| Tissue compartment: | Articular cartilage | | | | | | | |
|---|---|---|---|---|---|---|---|---|
| Parameter: | ACV | | Median AC Th | | Maximum AC Th | | AC Damage Area | |
| Unit: | (mm$^3$) | | (mm) | | (mm) | | (%) | |
| Technique: | ICE-µCT | | ICE-µCT | | ICE-µCT | | JSR | |
| Plateau | LTP | MTP | LTP | MTP | LTP | MTP | LTP | MTP |
| **Validation of methods by destabilisation of the medial meniscus (DMM) surgery** | | | | | | | | |
| Test: | T-test | Wilcox M-P SR | Wilcox M-P SR | Wilcox M-P SR | T-test | Wilcox M-P SR | Wilcox M-P SR | Wilcox M-P SR |
| Actual P-value (sham vs. DMM) | 0.185634186 | <0.001 | 0.022 | 0.079 | 0.250432301 | 0.017 | 0.011 | <0.001 |
| Annotation | ns | *** | * | ns | ns | * | * | *** |
| **Application 2: Age-related joint degeneration** | | | | | | | | |
| Test: | T-test | Wilcox RS | Wilcox RS | Wilcox RS | T-test | Wilcox RS | Wilcox RS | Wilcox RS |
| Actual P-value (4 months vs. 12 months) | 0.134192505 | 0.31 | 0.258 | 0.087 | 0.683908237 | 0.848 | 0.0022 | 0.026 |
| Annotation | ns | ns | ns | ns | ns | ns | ** | * |
| **Application 3: Mice with Dio2$^{Ala92Thr}$ polymorphisms** | | | | | | | | |
| Test: | T-test | Wilcox RS | Wilcox RS | Wilcox RS | T-test | Wilcox RS | Wilcox RS | Wilcox RS |
| Actual P-value (Dio2$^{Thr92}$ vs. Dio2$^{Ala92}$) | 0.819116125 | 0.019 | 0.701 | 0.006 | 0.238028137 | 0.088 | 0.043 | 0.016 |
| Annotation | ns | * | ns | ** | ns | ns | * | * |

| Tissue compartment: | Subchondral bone | | | | | | | | | |
|---|---|---|---|---|---|---|---|---|---|---|
| Parameter: | SC BV/TV | | SC Tb.Th | | SC Tb.N | | SC BMD | | SC BMC | |
| Unit: | (%) | | (mm) | | | | (mg.HA/cm$^3$) | | (median grey level) | |
| Technique: | ICE-µCT | | ICE-µCT | | ICE-µCT | | ICE-µCT | | scXRM | |
| Plateau | LTP | MTP | LTP | MTP | LTP | MTP | LTP | MTP | LTP | MTP |
| **Validation of methods by destabilisation of the medial meniscus (DMM) surgery** | | | | | | | | | | |
| Test: | T-test | Wilcox M-P SR | T-test | T-test | T-test | Wilcox M-P SR | T-test | Wilcox M-P SR | T-test | T-test |
| Actual P-value (sham vs. DMM) | 0.839865643 | <0.001 | 0.72881873 | 1.30518E-05 | 0.143962906 | 0.001 | 0.391493582 | 0.744 | 0.51033724 | 0.00047707 |
| Annotation | ns | *** | ns | *** | ns | *** | ns | ns | ns | *** |
| **Application 2: Age-related joint degeneration** | | | | | | | | | | |
| Test: | T-test | Wilcox RS | T-test | T-test | T-test | Wilcox RS | T-test | Wilcox RS | T-test | T-test |
| Actual P-value (4 months vs. 12 months) | 0.008239618 | 0.699 | 0.038538195 | 0.799467341 | 0.035389783 | 0.3095 | 0.151520392 | 0.2403 | 0.145885602 | 0.034329291 |
| Annotation | * | ns | * | ns | * | ns | ns | ns | ns | * |
| **Application 3: Mice with Dio2$^{Ala92Thr}$ polymorphisms** | | | | | | | | | | |
| Test: | T-test | Wilcox RS | T-test | T-test | T-test | Wilcox RS | T-test | Wilcox RS | T-test | T-test |
| Actual P-value (Dio2$^{Thr92}$ vs. Dio2$^{Ala92}$) | 0.672017104 | >0.999 | 0.87528521 | 0.796381673 | 0.070258368 | 0.15 | 0.624157343 | 0.604 | 0.185963192 | 0.31460005 |
| Annotation | ns | ns | ns | ns | ns | ns | ns | ns | ns | ns |

| | Histology sum OARSI score | | | | | Histology max. OARSI score | | | | |
|---|---|---|---|---|---|---|---|---|---|---|
| Plateau | MTP | MFC | LTP | LFC | Whole joint | MTP | MFC | LTP | LFC | Whole joint |
| **Validation of methods by destabilisation of the medial meniscus (DMM) surgery** | | | | | | | | | | |
| Test: | Wilcox M-P SR | Wilcox M-P SR | Wilcox M-P SR | Wilcox M-P SR | Wilcox M-P SR | Wilcox M-P SR | Wilcox M-P SR | Wilcox M-P SR | Wilcox M-P SR | Wilcox M-P SR |
| Actual P-value (sham vs. DMM) | 0.001 | 0.001 | 0.0469 | 0.2441 | 0.001 | 0.001 | 0.002 | 0.1094 | 0.2656 | 0.002 |
| Annotation | *** | *** | * | ns | *** | *** | ** | ns | ns | ** |
| **Application 2: Age-related joint degeneration** | | | | | | | | | | |
| Test: | | | Wilcox RS | | | | | Wilcox RS | | |
| Actual P-value (4 months vs. 12 months) | | | 0.065 | | | | | 0.024 | | |
| Annotation | | | ns | | | | | * | | |

| Key | |
|---|---|
| **P<0.05** | * |
| **P<0.01** | ** |
| **P<0.001** | *** |
| **Paired 2-tailed t-test** | T-test |
| **2-tailed Wilcoxon matched-pairs signed rank test** | Wilcox M-PSR |
| **2-tailed Wilcoxon rank sum test** | Wilcox RS |

***Supplementary Data 1***

#### ImageJ macros

All macros were generated using ImageJ1.44. Text between square brackets is descriptive and does not form part of the macro.

##### *Joint surface replication: quantification*

***Macro 1***
*[**Description:** This macro clears outside freehand selection of plateaux in all open images, saves as .tiff to selectable destination directory, appends 'plateau' to end of file name and closes images.]*

```
dir2 = getDirectory("Choose Destination Directory ");

setBatchMode(true);
imgArray = newArray(nImages);
for (i=0; i<nImages; i++) {
selectImage(i+1);
imgArray[i] = getTitle();
 }

 for (i=0; i< imgArray.length; i++) {
 selectImage(imgArray[i]);
 run("Clear Outside");
 saveAs("TIFF", dir2 +imgArray[i]+"_plateau");
 close();
 }
 setBatchMode(false);
```

***Macro 2***
*[**Description:** This macro removes bright outliers 4 pixels and below from all images in selectable source directory, saves as .tiff to selectable destination directory, appends '_RO ' to the end of the file name and closes images.]*

```
dir1 = getDirectory("Choose Source Directory ");
dir2 = getDirectory("Choose Destination Directory ");
list = getFileList(dir1);
setBatchMode(true);
for (i=0; i<list.length; i++) {
 showProgress(i+1, list.length);
 open(dir1+list[i]);
run("Remove Outliers...", "radius=4 threshold=50 which=Bright");
 saveAs("TIFF", dir2+list[i]+"_RO");
 close();
}
```

***Macro 3***
[**Description:** *This macro finds edges on images of plateaux in selectable source directory, saves as .tiff to selectable destination directory and appends 'edges' to the file name.]*

```
dir1 = getDirectory("Choose Source Directory ");
dir2 = getDirectory("Choose Destination Directory ");
list = getFileList(dir1);
setBatchMode(true);
for (i=0; i<list.length; i++) {
 showProgress(i+1, list.length);
 open(dir1+list[i]);
run("Find Edges");
 saveAs("TIFF", dir2+list[i]+"edges");
 close();
}
```

***Macro 4***
[**Description:** *This macro saves all open files into selectable destination directory, appends 'cleaned' to the title name, and closes images.]*

```
dir2 = getDirectory("Choose Destination Directory ");
setBatchMode(true);
imgArray = newArray(nImages);
for (i=0; i<nImages; i++) {
selectImage(i+1);
imgArray[i] = getTitle();
 }
 for (i=0; i< imgArray.length; i++) {
 selectImage(imgArray[i]);
saveAs("TIFF", dir2 +imgArray[i]+"_cleaned");
close();
}
 setBatchMode(false);
```

***Macro 5***
*[**Description:** This macro saves all open images as .tiff to selectable destination directory, appends 'Thresh ' to file name and closes images.]*

```
dir2 = getDirectory("Choose Destination Directory ");

setBatchMode(true);
imgArray = newArray(nImages);
for (i=0; i<nImages; i++) {
selectImage(i+1);
imgArray[i] = getTitle();
 }
 for (i=0; i< imgArray.length; i++) {
 selectImage(imgArray[i]);
saveAs("TIFF", dir2 +imgArray[i]+"_Thresh");
close();
}
 setBatchMode(false);
```

***Macro 6***
[**Description:** *This macro analyses all images in selectable source folder, selects all particles with size 20-infinity and circularity 0-0.5 and saves outlines as new image.tiff in selectable destination directory.]*

```
dir1 = getDirectory("Choose Source Directory ");
dir2 = getDirectory("Choose Destination Directory ");
list = getFileList(dir1);
setBatchMode(true);
for (i=0; i<list.length; i++) {
 showProgress(i+1, list.length);
 open(dir1+list[i]);
//run("Threshold...");
setAutoThreshold("Minimum dark");
setThreshold(253, 255);
run("Convert to Mask");
run("Analyze Particles...", "size=20-Infinity circularity=0.00-0.5 show=[Bare Outlines]
display summarize");
saveAs("Tiff", dir2+list[i]+"_bare outlines");
close();
}
```

#### *Subchondral X-ray Microradiography (scXRM)*

***Macro 7***
[**Description:** *This macro stretches pixel information between the pixel value for the lowest-density standard (plastic) and the highest density standard (steel), applies a 16-colour look-up table and saves as a new image.]*

```
Dialog.create("Set Standards");
Dialog.addNumber("Steel mode grey level", 779, 0, 5, "");
Dialog.addNumber("Plastic mode grey level", 8769, 0, 5, "");
Dialog.addNumber("Minimum grey level", 0, 0, 5, "");
Dialog.addNumber("Maximum grey level", 12031, 0, 5, "");
Dialog.show;

steel = Dialog.getNumber;
plastic = Dialog.getNumber;
low = Dialog.getNumber;
high = Dialog.getNumber;

strl = steel
strh = plastic

min=low+(high-plastic)
max=high-(steel-low)

setBatchMode(true);
setMinAndMax(min,max);
run("Invert");
```

```
original = getTitle();
width = getWidth();
height = getHeight();
setBatchMode(true);
pix = newArray(width*height);
for (y=0; y<height; y++){
    for (x=0; x<width; x++){
        pix[y*width+x] =   (getPixel(x,y)-min)*255/(max-min);
    }
}
close();
newImage(original+"_normalised"+" "+strl+"_"+strh+" 16colour.tif", "8-bit black", width,
height, 1);
selectWindow(original+"_normalised"+" "+strl+"_"+strh+" 16colour.tif");
for (y=0; y<height; y++){
    for (x=0; x<width; x++){
        setPixel(x,y, pix[y*width+x]);
    }
}

run("16_colors");
setBatchMode(false);
```
